# NAD metabolism modulates inflammation and mitochondria function in diabetic kidney disease

**DOI:** 10.1101/2021.12.05.471273

**Authors:** Komuraiah Myakala, Xiaoxin X Wang, Nataliia V. Shults, Bryce A. Jones, Xiaoping Yang, Avi Z Rosenberg, Brandon Ginley, Pinaki Sarder, Leonid Brodsky, Yura Jang, Chan Hyun Na, Yue Qi, Xu Zhang, Udayan Guha, Ci Wu, Shivani Bansal, Junfeng Ma, Amrita Cheema, Chris Albanese, Matthew D Hirschey, Teruhiko Yoshida, Jeffrey B. Kopp, Julia Panov, Moshe Levi

## Abstract

Diabetes mellitus is the leading cause of cardiovascular and renal disease in the United States. In spite of the beneficial interventions available for patients with diabetes, there remains a need for additional therapeutic targets and therapies in diabetic kidney disease (DKD). Inflammation and oxidative stress are increasingly recognized as important causes of renal diseases. Inflammation is closely associated with mitochondrial damage. The molecular connection between inflammation and mitochondrial metabolism remains to be elucidated. Recently, nicotinamide adenine nucleotide (NAD+) metabolism has been found to regulate immune function and inflammation. In the present studies we tested the hypothesis that enhancing NAD metabolism could prevent inflammation in and progression of DKD. We found that treatment of *db/db* mice with type 2 diabetes with nicotinamide riboside (NR) prevented several manifestations of kidney dysfunction (i.e., albuminuria, increased urinary kidney injury marker-1 (KIM1) excretion and pathologic changes). These effects were associated with decreased inflammation, at least in part via inhibiting the activation of the cyclic GMP-AMP synthase-stimulator of interferon genes (cGAS-STING) signaling pathway. An antagonist of the serum stimulator of interferon genes (STING) and whole-body STING deletion in diabetic mice showed similar renoprotection. Further analysis found that NR increased SIRT3 activity and improved mitochondrial function, which led to decreased mitochondrial DNA damage, a trigger for mitochondrial DNA leakage which activates the cGAS-STING pathway. Overall, these data show that NR supplementation boosted NAD metabolism to augment mitochondrial function, reducing inflammation and thereby preventing progression of diabetic kidney disease.

## INTRODUCTION

Diabetes mellitus is the leading cause of cardiovascular and renal disease in the United States ^1–5^. The National Diabetes Statistics Report in 2020 estimated that more than 34 million Americans, or 10.5% of the population, had diabetes (https://www.cdc.gov/diabetes/pdfs/data/statistics/national-diabetes-statistics-report.pdf). Further, as many as one in four Americans are expected to become diabetic by the year 2050 ^4, 6^.

In spite of the beneficial interventions implemented in patients with diabetes, including tight glucose control, stringent blood pressure control, angiotensin-converting enzyme inhibition (ACEI), angiotensin II receptor blockade (ARB), mineralocorticoid receptor antagonism, sodium glucose cotransport-2 (SGLT2) inhibition and glucagon-like receptor protein-1 (GLP-1) receptor agonism ^7–12^, the new therapeutic targets in DKD are emerging based on the further understanding of the mechanisms to cause progression and/or prevention of DKD.

Inflammation and oxidative stress are increasingly recognized as important causes of renal diseases ^13, 14^. In animal models of DKD, such as db/db mice or KKAy mice, and in diet induced obesity mice, there are increased renal inflammation and oxidative stress ^15, 16^. Inflammation is closely associated with mitochondria damage ^17, 18^. In kidneys from DKD, there is a wide variety of mitochondrial dysfunction reported ^15, 16, 19, 20^. The molecular connection between inflammation and mitochondrial metabolism remains to be elucidated. Recently, nicotinamide adenine nucleotide (NAD+) metabolism has been found to regulate immune function and inflammation ^21–25^.

In the present studies, we administered a NAD+ booster, nicotinamide riboside (NR) to db/db mice, a model of type 2 diabetes, to evaluate its effects in diabetic nephropathy. We found that long-term NR treatment improved DKD in the db/db mice via preventing the renal inflammation, at least in part by inhibiting the activation of the cyclic GMP-AMP synthase-stimulator of interferon genes (cGAS-STING) signaling pathway. This anti-inflammatory activity was associated with mitochondrial function restoring.

## RESULTS

### NR treatment improved murine diabetic kidney disease

We treated db/db mice with type 2 diabetes with NR for 20 weeks (**Figure 1A**). NR treatment did not affect body weight, kidney weight, or blood glucose, but serum cholesterol and triglyceride levels were decreased in NR treated db/db mice (**Table 1**). We found that NR treated db/db mice had a significant decrease in albuminuria (**Figure 1B**) and excretion of the urinary kidney injury marker Kim1 (**Figure 1C**).

**Figure 1.**
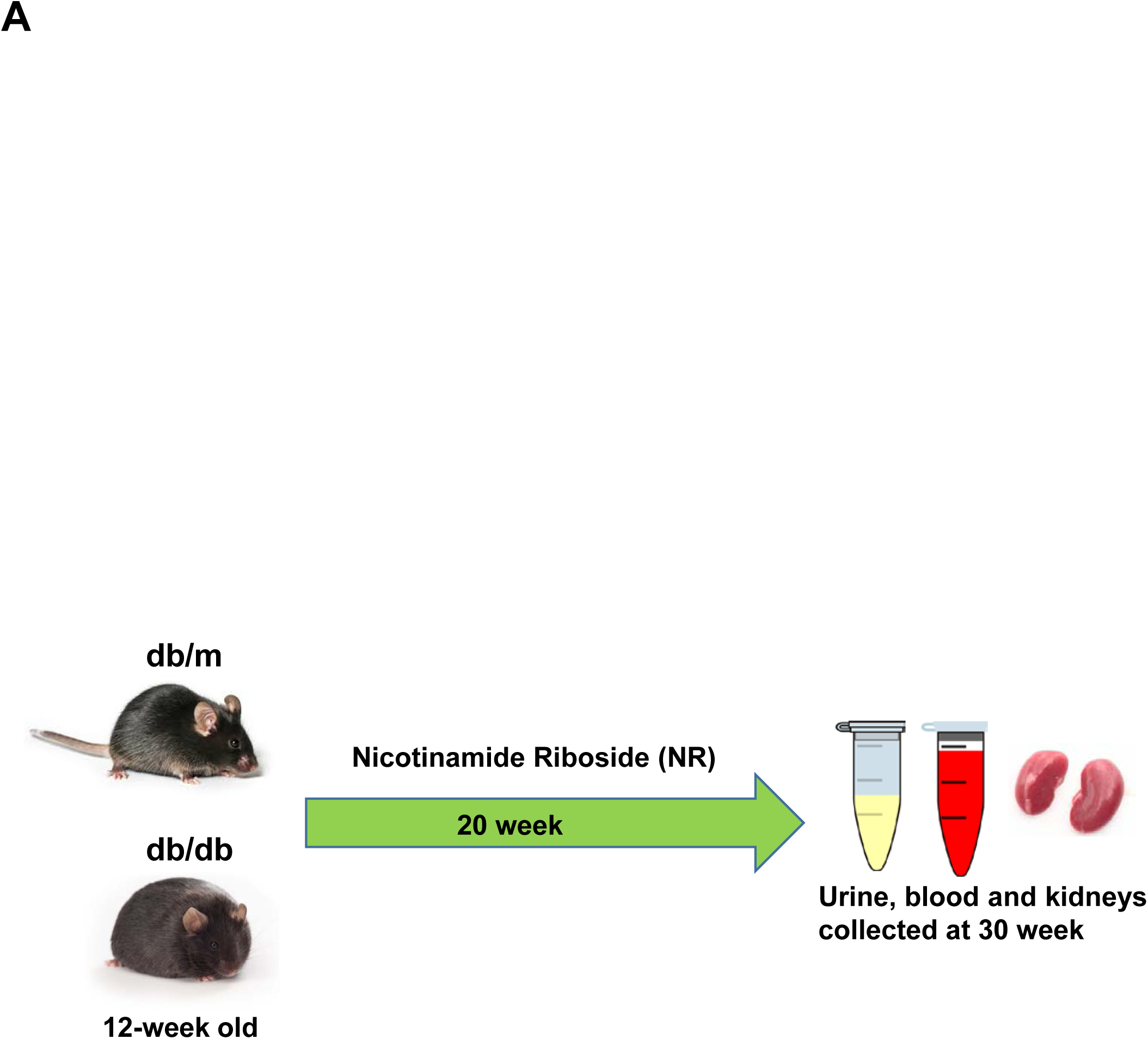
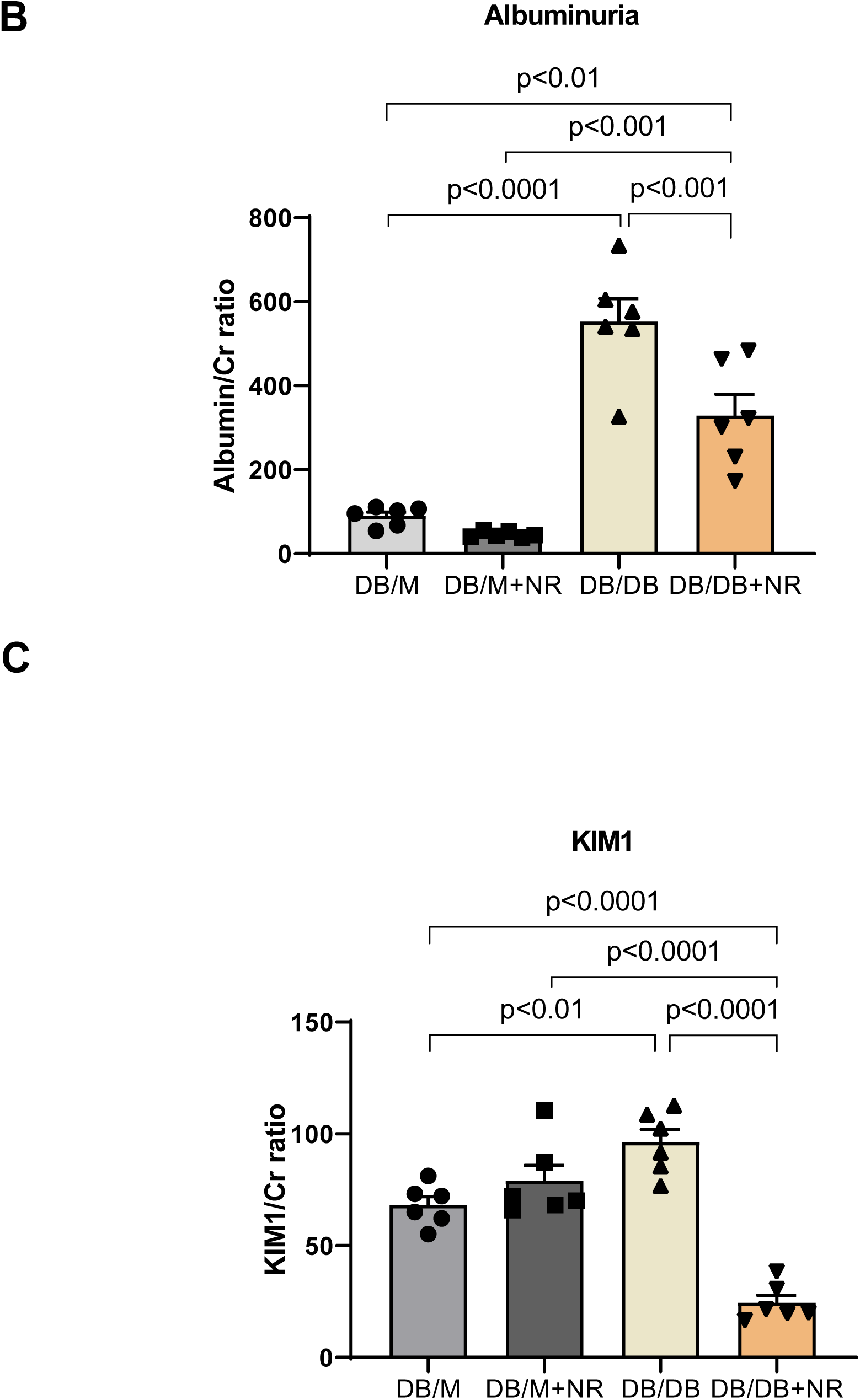
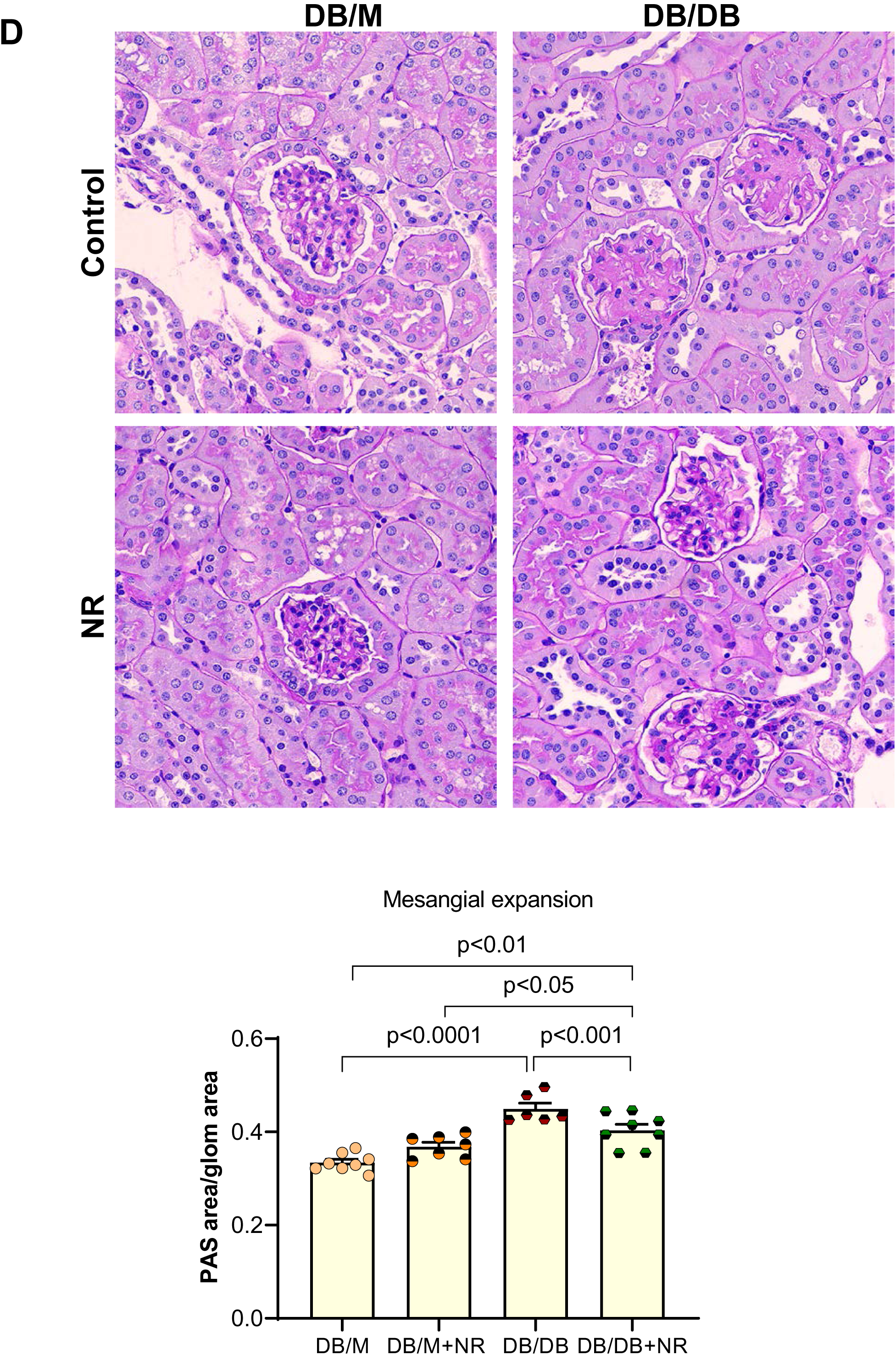
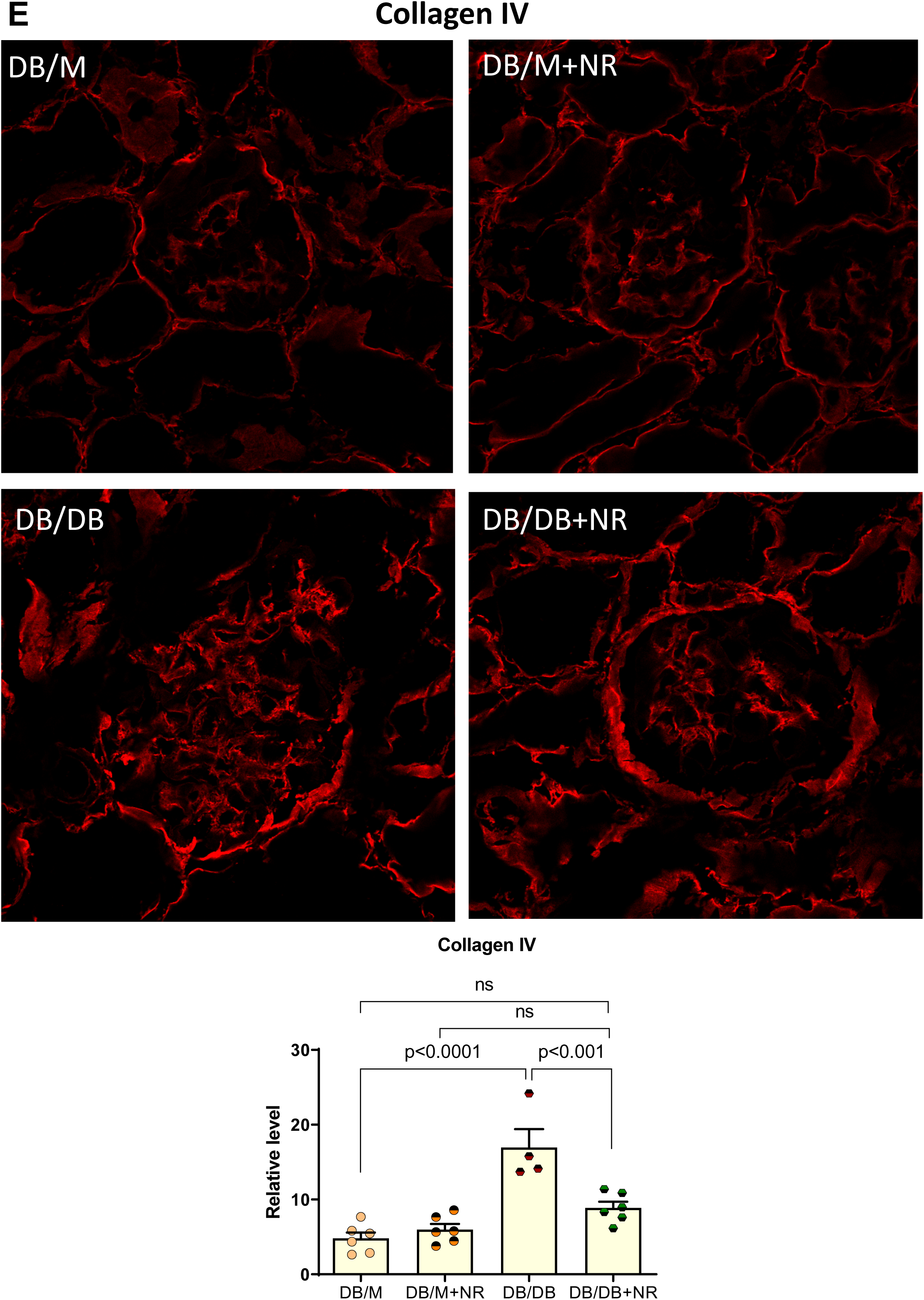
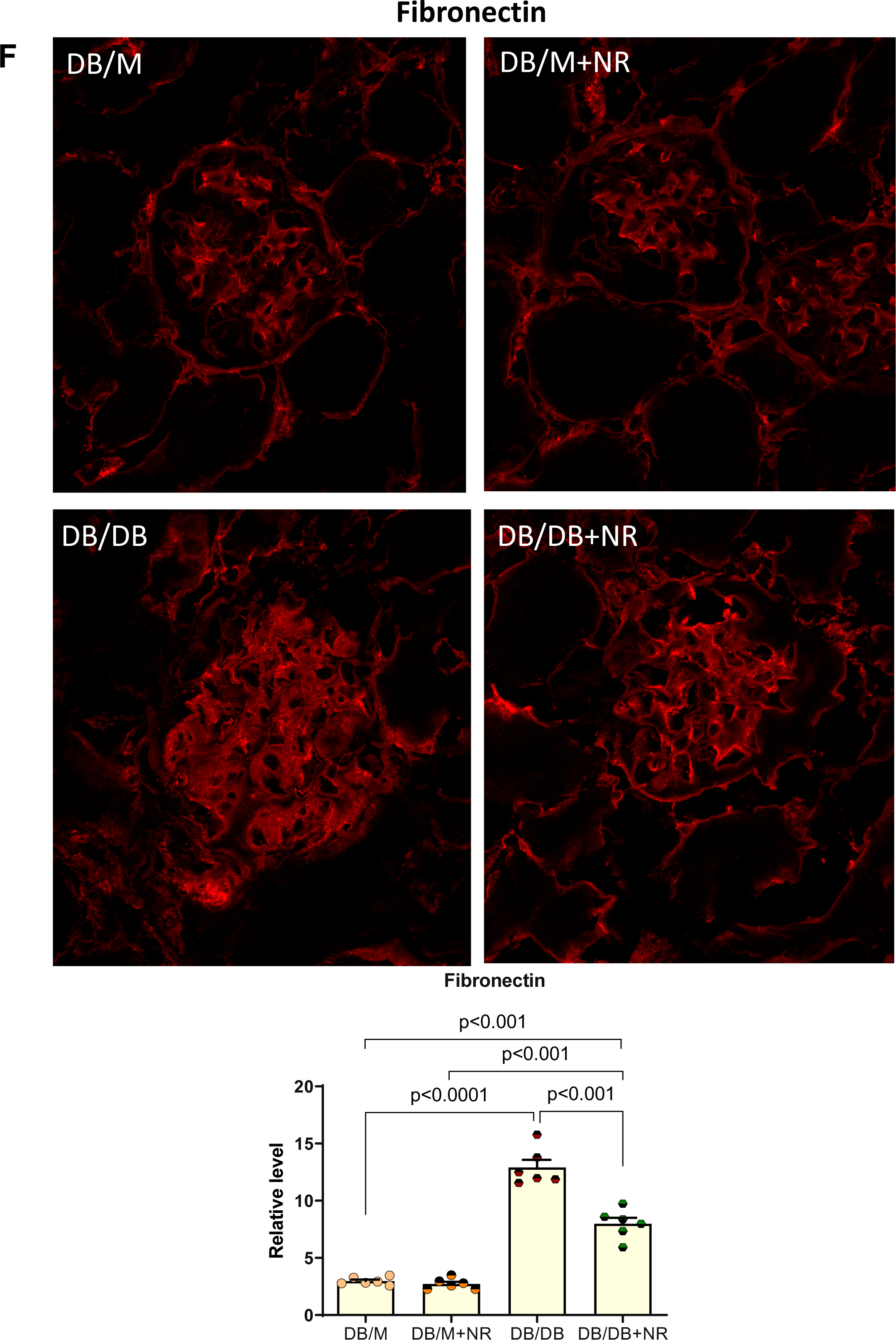
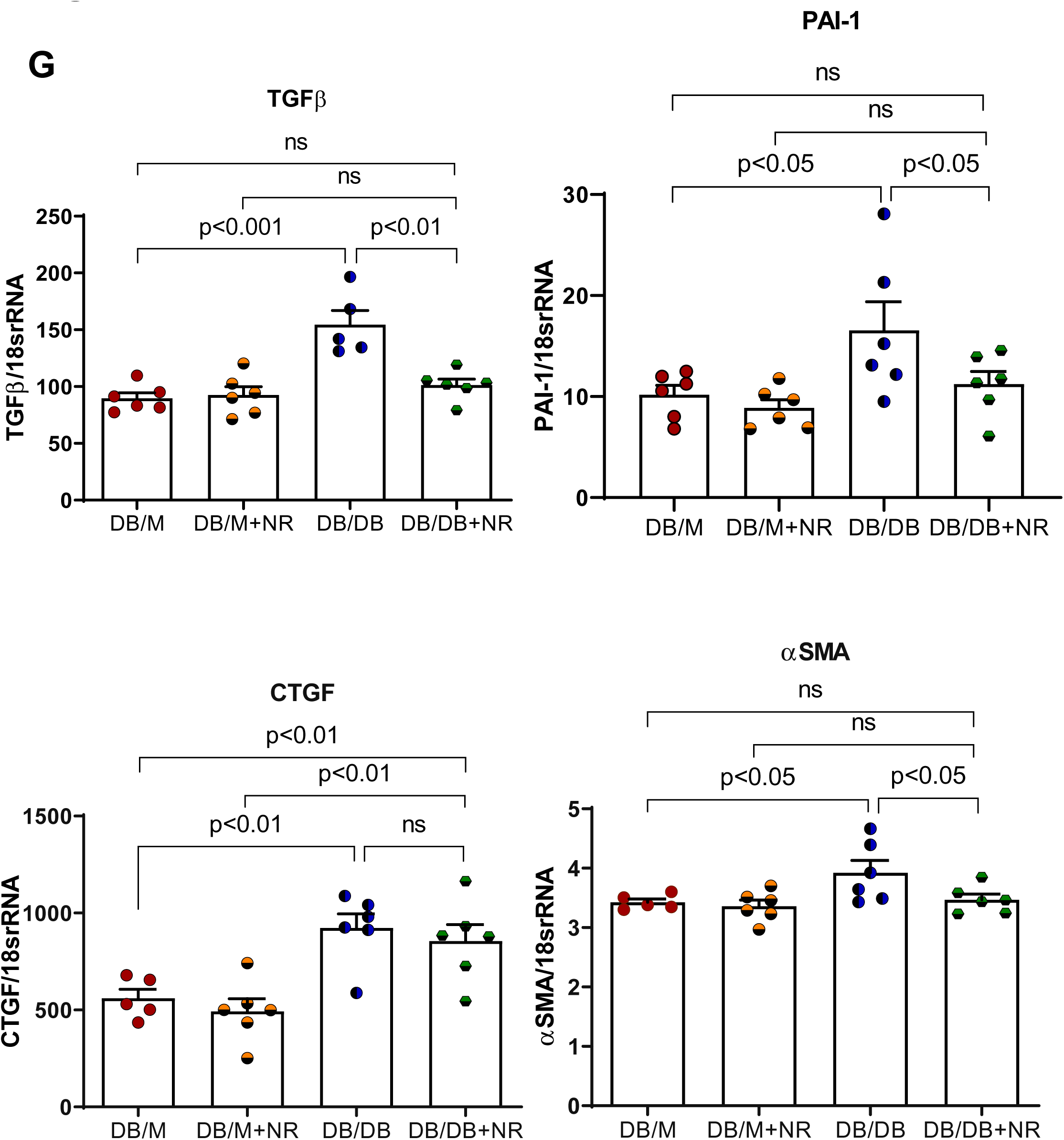
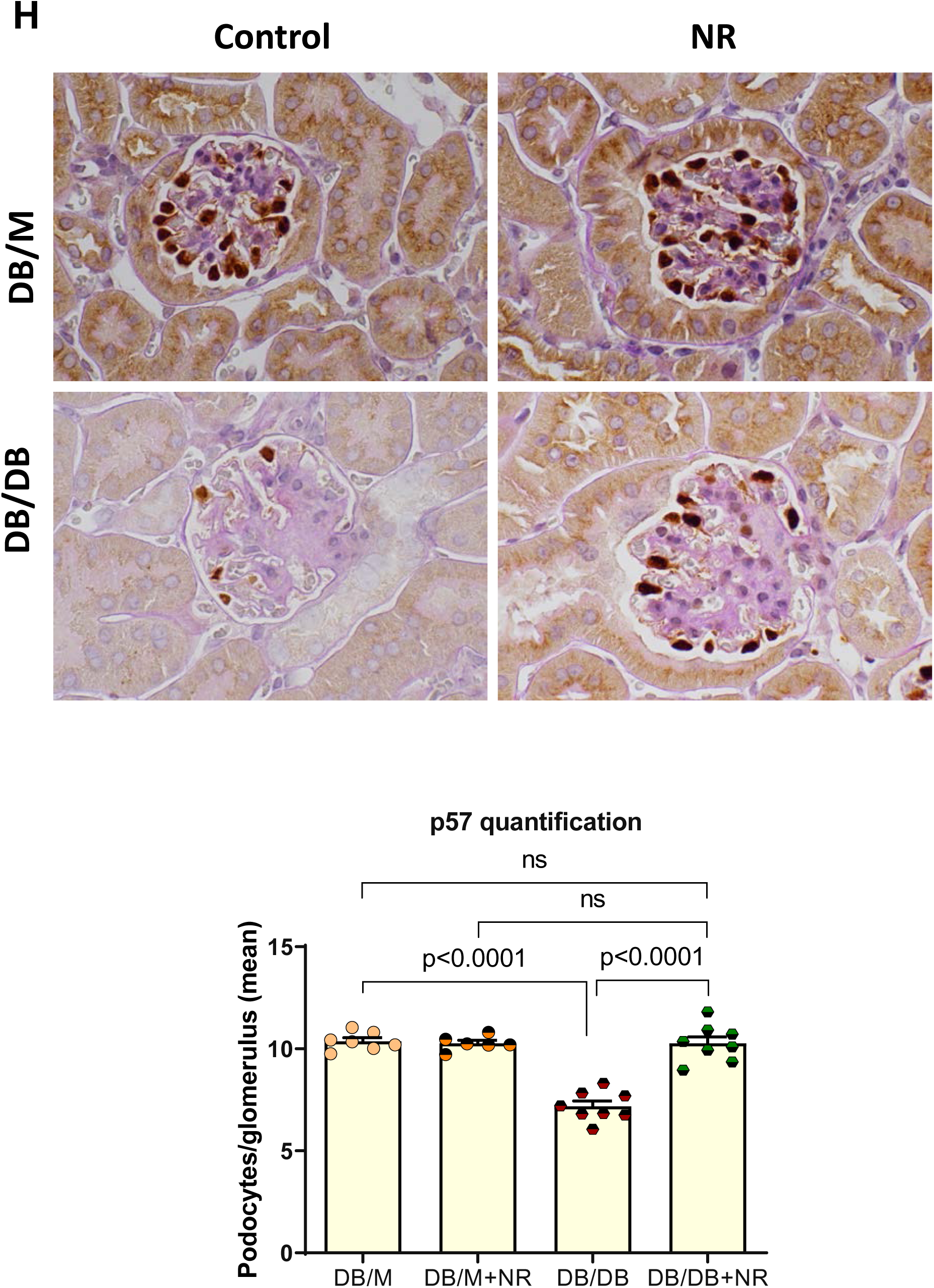
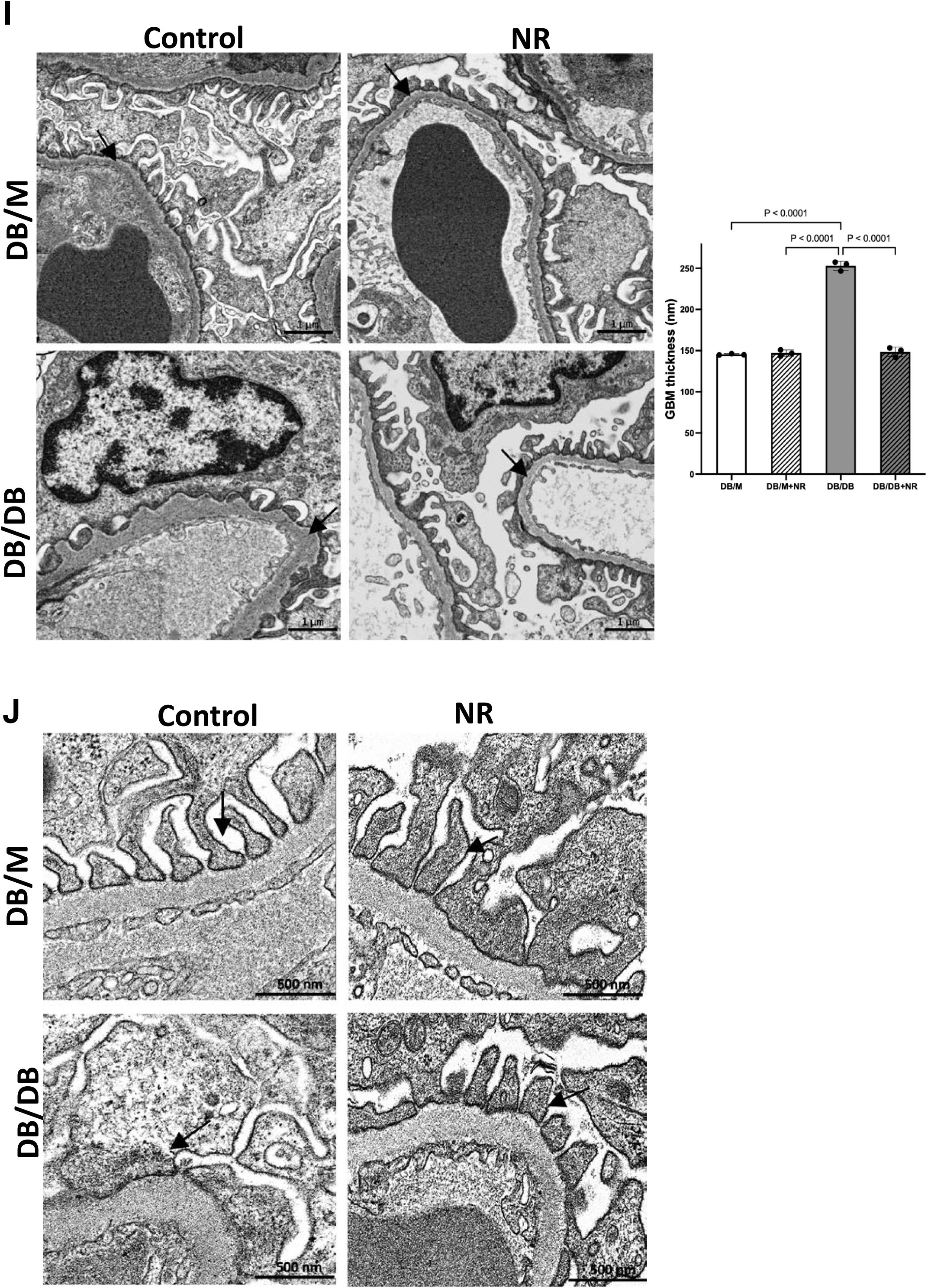
Effect of nicotinamide riboside (NR) treatment on diabetes induced podocyte and tubular dysfunction. A) Experimental scheme for the study with NR treatment. B) Urinary albumin and C) KIM1 normalized to creatinine ratio was measured in 24-hour urine. Albumin and KIM1 excretion were significantly increased in db/db compared to non-diabetic db/m mice. Dietary supplementation with NR significantly reduced the urinary excretion of albumin and KIM1. D) PAS stained images indicating, increased PAS positive area with mesangial matrix expansion, glomeruli size and pathological changes were suppressed upon NR treatment in db/db mice with unchanged glomerular size. E) & F) Immunofluorescence images indicating the collagen IV and fibronectin fluorescence signals were increased in db/db kidney and reduced significantly in NR treated mice. G) TGFβ, PAI1, CTGF and αSMA mRNA levels were higher in the db/db kidneys. TGFβ, PAI1, and αSMA mRNA levels were significantly decreased upon NR treatment. H) Immunochemistry images of podocyte positive marker p57 is markedly reduced in kidney of db/db mice and depletion was prevented upon NR treatment. p57 positive areas were quantified per each glomerulus of kidney section. n=6-8 per group, values presented as mean ± SEM with variance is calculated using one-way ANOVA. I) Representative SEM images of glomerular basement membrane (GBM) thickening with quantification showing an increased GBM thickness in the db/db kidney, which was normalized with NR treatment. Arrows indicate the GBM. Scale bars 1 um. Magnifications x 35,000. n=3 per group. J) TEM images demonstrate the normal structure of podocyte foot processes (arrows) in db/m controls and db/m treated with NR. db/db mice showed podocyte foot processes effacement (arrow) and reduction in filtration slits frequency. Scale bars 500 nm. Magnifications x 80,000. The graphs show the widening of podocyte foot processes, a reduction in filtration slit frequency in db/db and amelioration of those parameters in DB/DB with NR treatment. n=3 per group.

**Table 1.** Metabolic data for db-m and db-db mice. Data are means ± SE (N=6 mice in each group): ª p < 0.0001 vs. DB/M, ^b^ p < 0.001 vs. DB/M, ^c^ p < 0.05 vs. DB/M, ^d^ p < 0.05 vs. DB/DB.

|  | DB/M | DB/M + NR | DB/DB | DB/DB + NR |
| --- | --- | --- | --- | --- |
| Body weight (g) | 30.8±2.28 | 31.7±1.67 | 33.8±2.7 | 27.8±2.39 |
| Food intake (mouse/day) | 3.1±0.06 | 3.03±0.06 | 5.4±2.60<sup>a</sup> | 5.2±0.21 |
| Kidney weight (g) | 0.29±0.02 | 0.24±0.02 | 0.31±0.02 | 0.26±0.02 |
| Kidney weight/body weight ratio (%) | 0.90±0.11 | 0.77±0.1 | 0.98±0.11 | 0.94±0.1 |
| glucose (mg/dL) | 145.8±11 | 156.8±7.78 | 783.3±29<sup>a</sup> | 787.2±3.83 |
| Plasma TG (mg/dL) | 55.6±15.2 | 46.9±8.7 | 137.1±15.2<sup>a</sup> | 72.0±64.9<sup>b</sup> |
| Plasma TC (mg/dL) | 86.5±2.3 | 84.2±9.6 | 121±34.4<sup>c</sup> | 89.2±31.7<sup>d</sup> |

Data are means ± SE (N=6 mice in each group): <sup>a</sup> p < 0.0001 vs. DB/M, <sup>b</sup> p < 0.001 vs. DB/M, <sup>c</sup> p < 0.05 vs. DB/M, <sup>d</sup> p < 0.05 vs. DB/DB.

NR treatment reduced mesangial expansion in db/db mice, assessed on PAS staining (**Figure 1D**). Immunofluorescence microscopy showed increased extracellular matrix protein (collagen IV and fibronectin) deposition in the glomeruli of db/db kidneys indicating the presence of glomerulosclerosis, which was improved with NR treatment (**Figure 1E-F, Supplementary Figure S1**). RNA expression of profibrotic factors, including transforming growth factor (TGF)-β1, plasminogen activator inhibitor (PAI)-1 and connective tissue growth (CTGF), and α−smooth muscle actin (αSMA), were increased in db/db kidneys. Treatment with NR decreased mRNA abundance of TGFβ1, PAI-1 and αSMA, but not CTGF (**Figure 1G**).

We next examined podocyte loss with the podocyte nuclear marker p57. Immunohistochemistry analysis of p57 showed decreased podocyte numbers in db/db kidneys. This loss was prevented by NR treatment (**Figure 1H**).

Ultrastructural examination demonstrated irregular glomerular basement membrane (GBM) thickening in db/db mice, which was ameliorated with NR treatment (**Figure 1I**). Finally, podocyte foot processes effacement was notable in db/db mice and preserved with NR treatment (**Figure 1J**).

Taken together, we concluded that NR prevent effects in protecting podocyte and glomerular integrity and preventing tubulointerstitial injury in a model of diabetic kidney disease.

### NR treatment improved renal oxidative stress

We examined levels of thiobarbituric acid reactive substances (TBARS), which represent oxidation products of lipid. We found that urinary TBARS level, kidney NADPH oxidase 4 (NOX4) mRNA and kidney 4-hydroxynonenal (4-HNE) protein levels-markers of lipid peroxidation, were all increased in db/db mice. NR treatment decreased urinary TBARS level, and kidney NOX4 mRNA and 4-HNE protein levels in db/db mice (**Figures 2A-C**).

**Figure 2:**
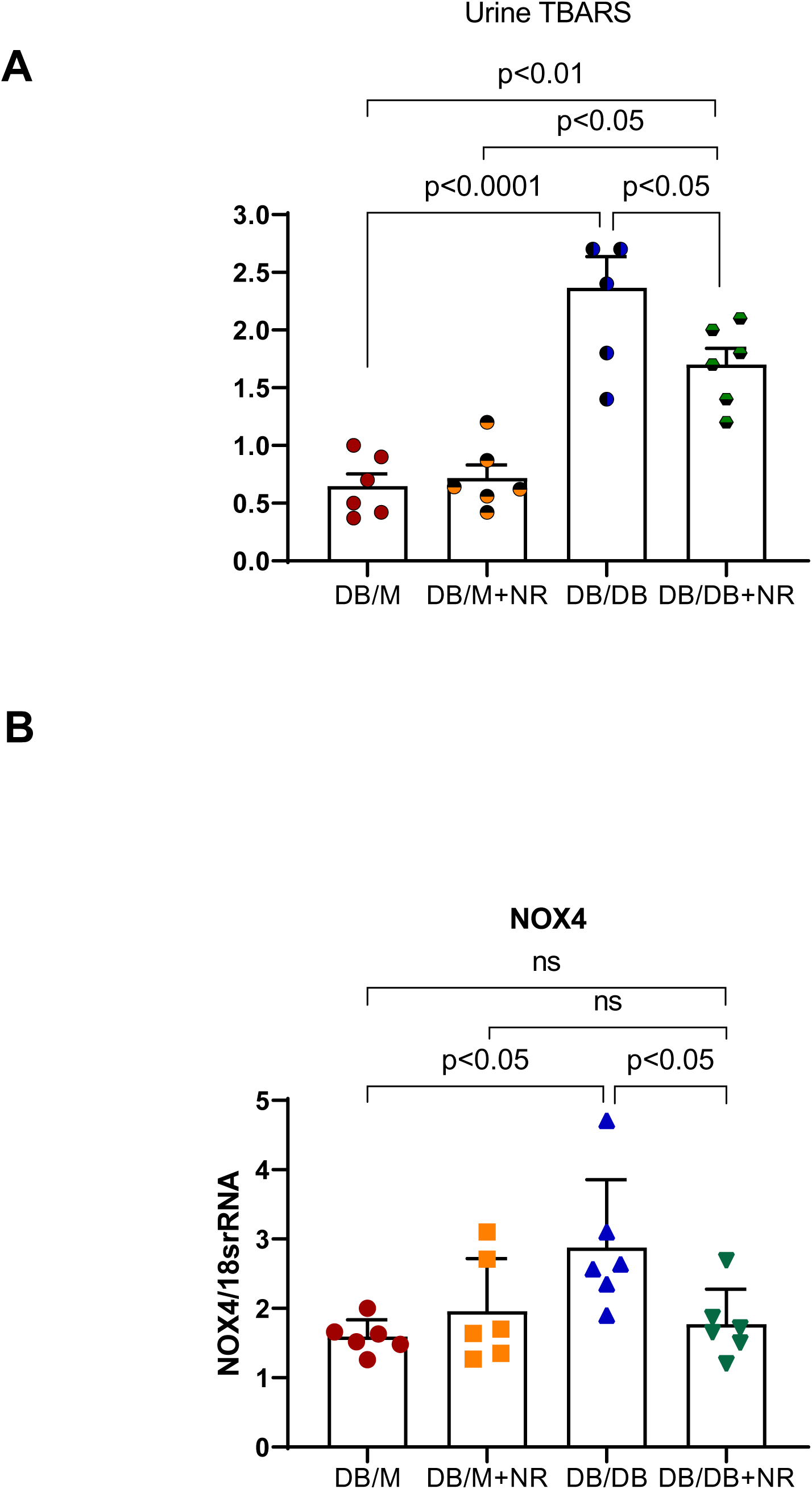
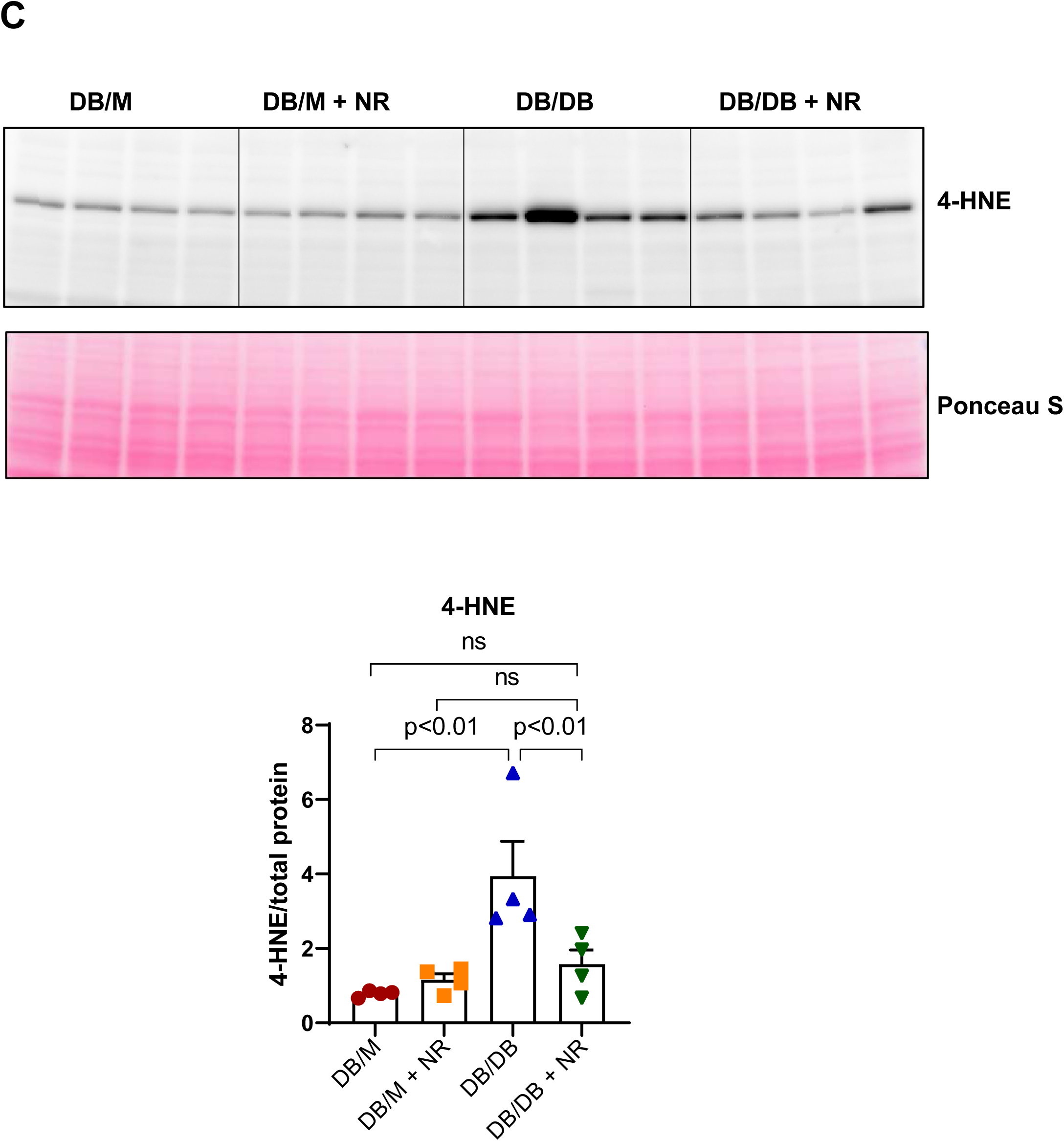
Effect of NR treatment on markers related to oxidative stress in diabetic mice. A) Urinary thiobarbituric acid reactive substances (TBARS), B) NADPH oxidase 4 (NOX4) mRNA levels were significantly higher in db/db mice and NR treatment prevented increases in TBARS and NOX4 mRNA. C) Western blot analysis showing 4-hydroxynonenal (4-HNE) protein expression levels were upregulated and abrogated upon NR treatment in kidney of db/db mice. n=6 per group, values presented as mean ± SEM with variance is calculated using one-way ANOVA.

### NR treatment decreased inflammation in diabetic kidneys

We found increased expression of monocyte chemoattractant protein (MCP)-1, tumor necrosis factor (TNF), interleukin (IL)-6, tissue inhibitor of metalloproteinase (TIMP)1 and CD68 mRNA in diabetic kidneys. Expression of toll-like receptor 2 (TLR2), a marker of innate immune pathway triggered by damage-associated molecular patterns, was also increased in db/db kidneys. NR treatment effectively prevented these increases (**Figure 3A**). Staining of CD45 and CD68, markers of leukocytes and macrophages, respectively, showed glomerular infiltration of immune cells in db/db kidney, which was prevented by NR treatment (**Figure 3B and 3C**). Thus we demonstrate the role of NR treatment on reducing inflammatory pathways in the db/db model.

**Figure 3:**
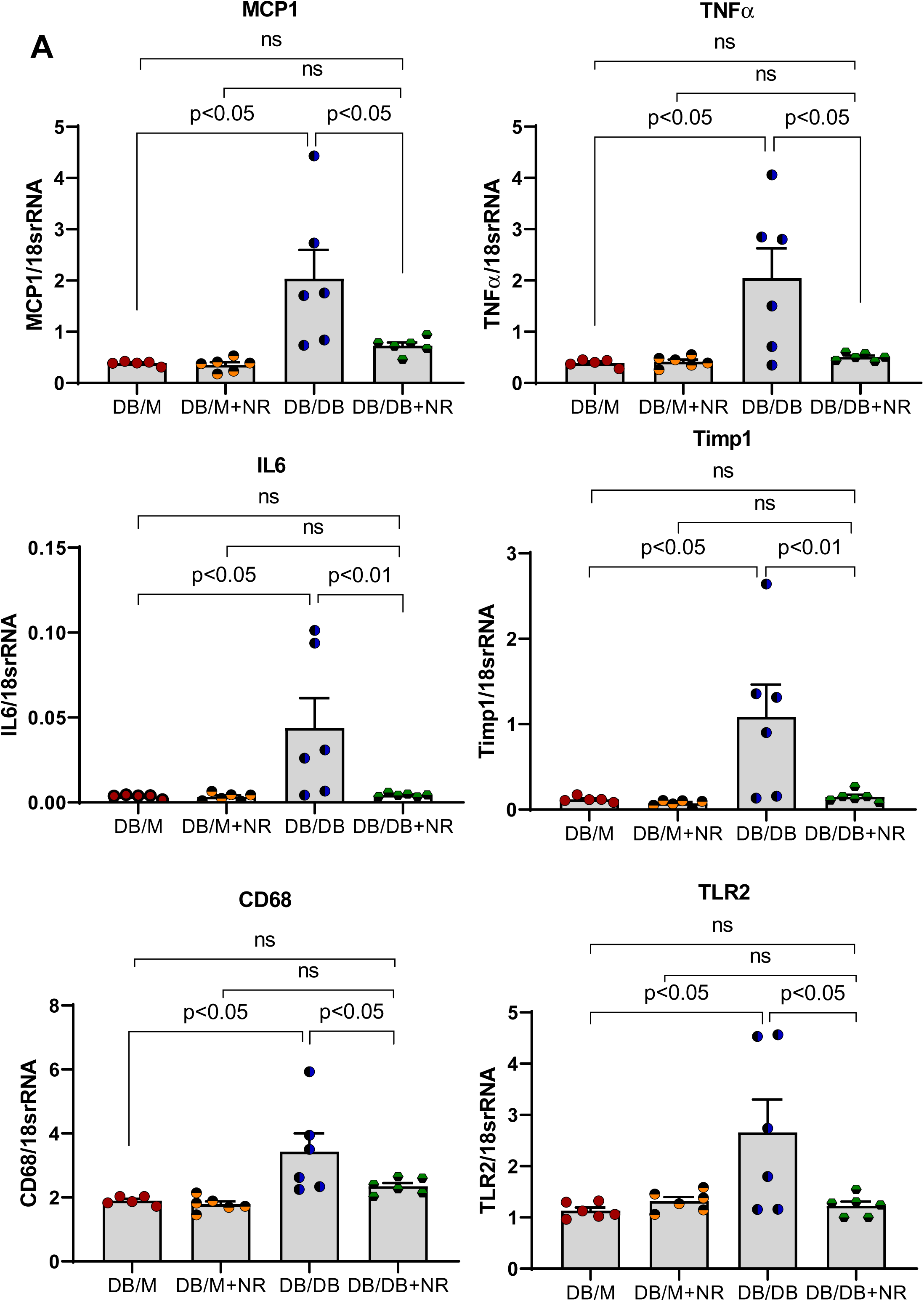
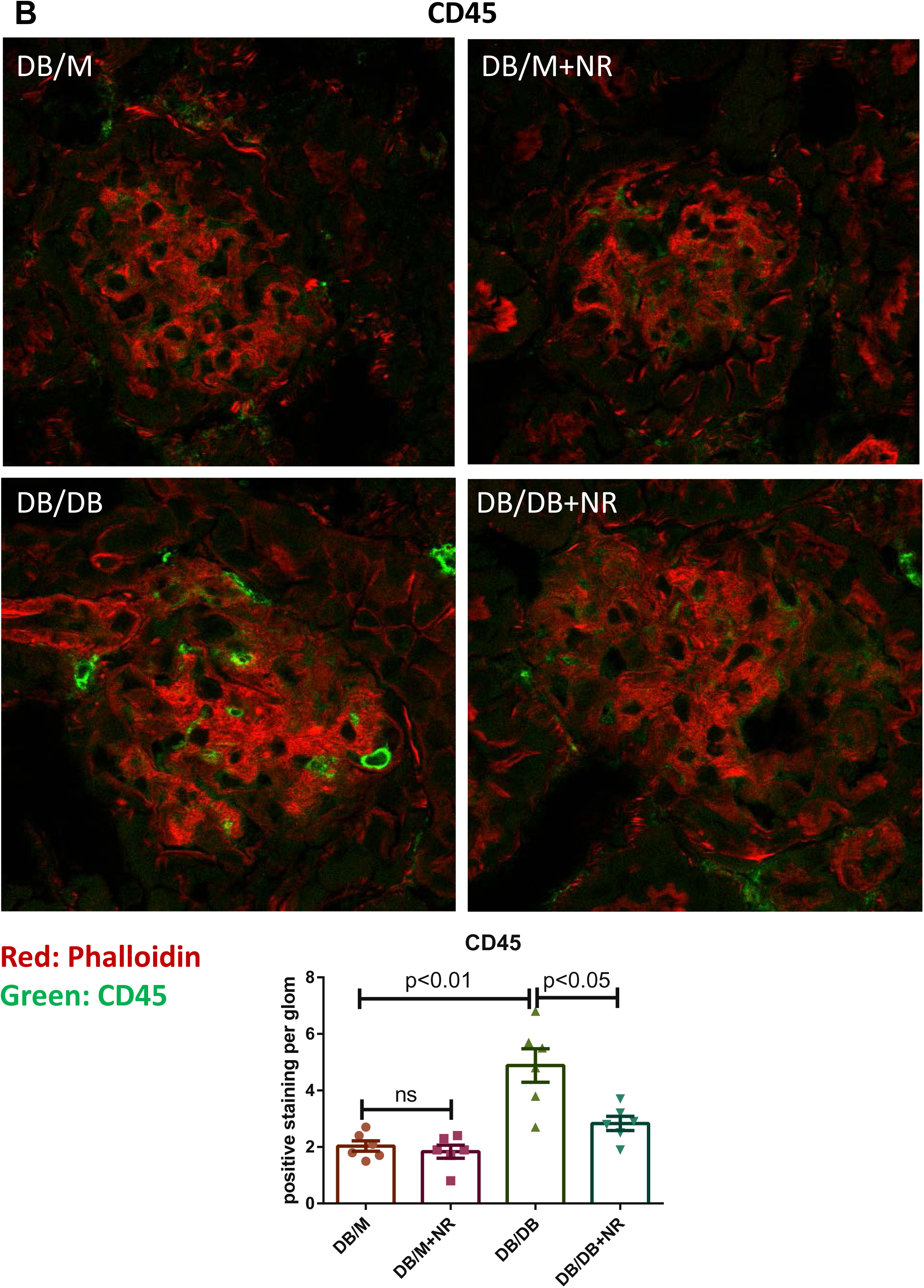
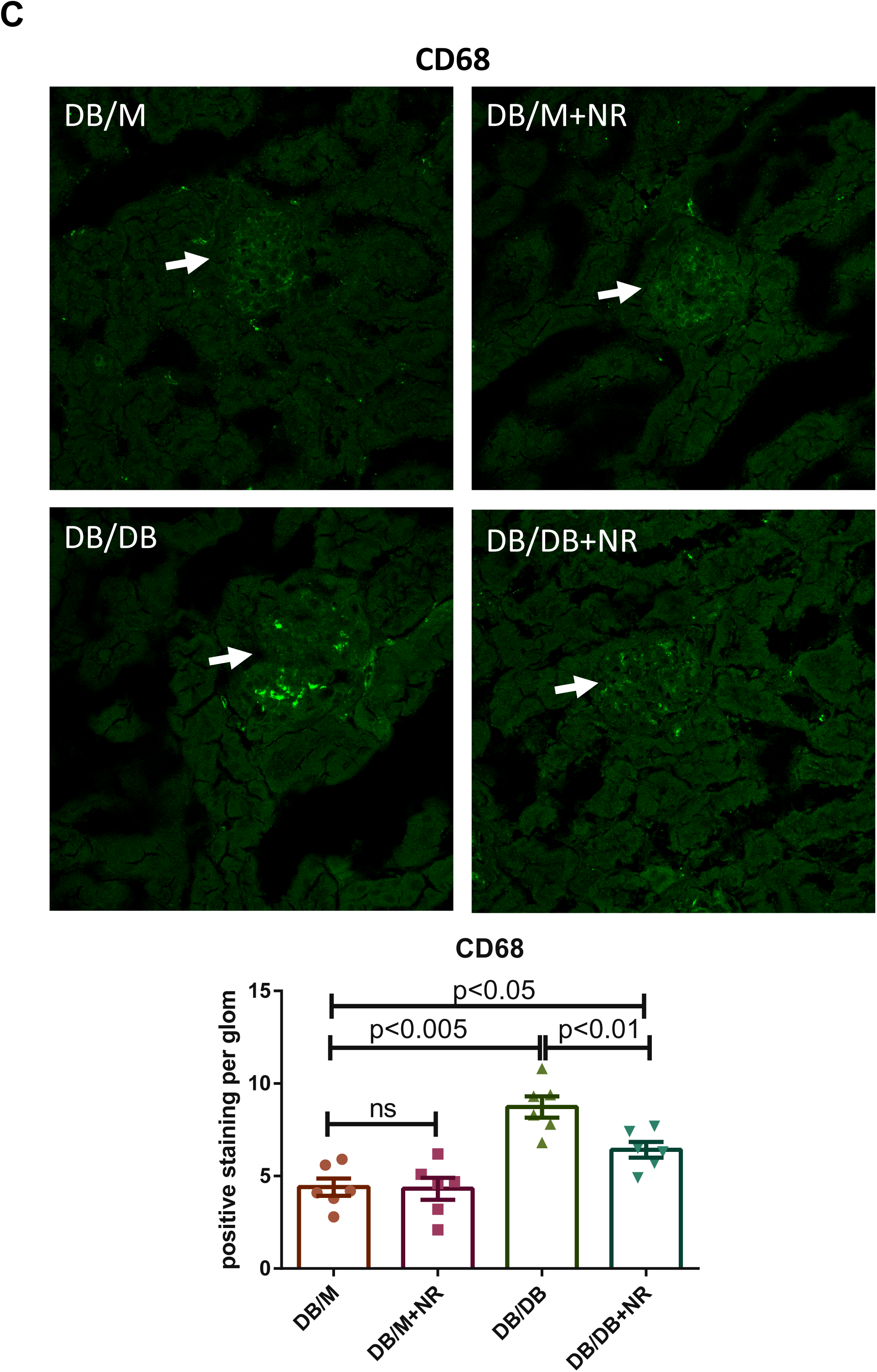
Effect of NR treatment on inflammation in the kidney of db/db mice. A) The inflammatory cytokine markers MCP-1, TNFα, IL-6, TIMP1 and CD68 transcript levels were significantly higher in db/db mice. Also TLR2 an innate immune response marker increased in db/db mice. NR treatment successfully prevented their increases. Immunofluorescence microscopic images B) CD45 stained in green, phalloidin as red and C) CD68 stained in green showing the infiltration of immune cells in the glomeruli of db/db kidney which was prevented in the NR treated db/db mice. n=5-6 per group, values presented as mean ± SEM with variance is calculated using one-way ANOVA.

### NR treatment decreased cGAS-STING activation in diabetic kidneys

To explore how NR treatment exerts anti-inflammatory effects, we analyzed bulk RNAseq and proteomics data. We found that NR treatment downregulated many of the inflammatory genes and proteins upregulated in db/db kidneys (**Figure 4 A-C, Supplementary Tables S1 and Figure S2**). Interestingly, the expression of interferon-induced transmembrane (IFITM) genes ^27^, members of the interferon stimulated gene (ISG) family, was upregulated in db/db kidneys and expression of these genes was reduced by NR treatment (**Figure 4D**). Aside from viral infections, ISG induction can be triggered by activated nucleic acid sensors, such as cGAS/STING, which respond to the leak of nuclear or mitochondrial DNA into the cytoplasm ^28–30^. In diabetic kidneys, we found marked increases in the cGAS mRNA, STING mRNA and STING protein levels. cGAS and STING levels were significantly decreased by treatment with NR (**Figure 4E**). Activation of STING activates downstream effectors TBK1 and IRF3, by promoting their phosphorylation. We detected increased phosphorylation of TBK1 **(Figure 4F)** and IRF3 **(Figure 4G)** in db/db mice that was significantly reduced by NR treatment, confirming modulation of STING pathway signaling in an NR-dependent manner (**Figure 4F-G**). We further examined the downstream response and we found that both STAT3 **(Figure 4H)** and NFκB **(Figure 4I)** were activated in db/db kidneys. Their activation as determined by the increased levels of phospho-STAT3 and phospho-p65, was reduced by NR treatment.

**Figure 4:**
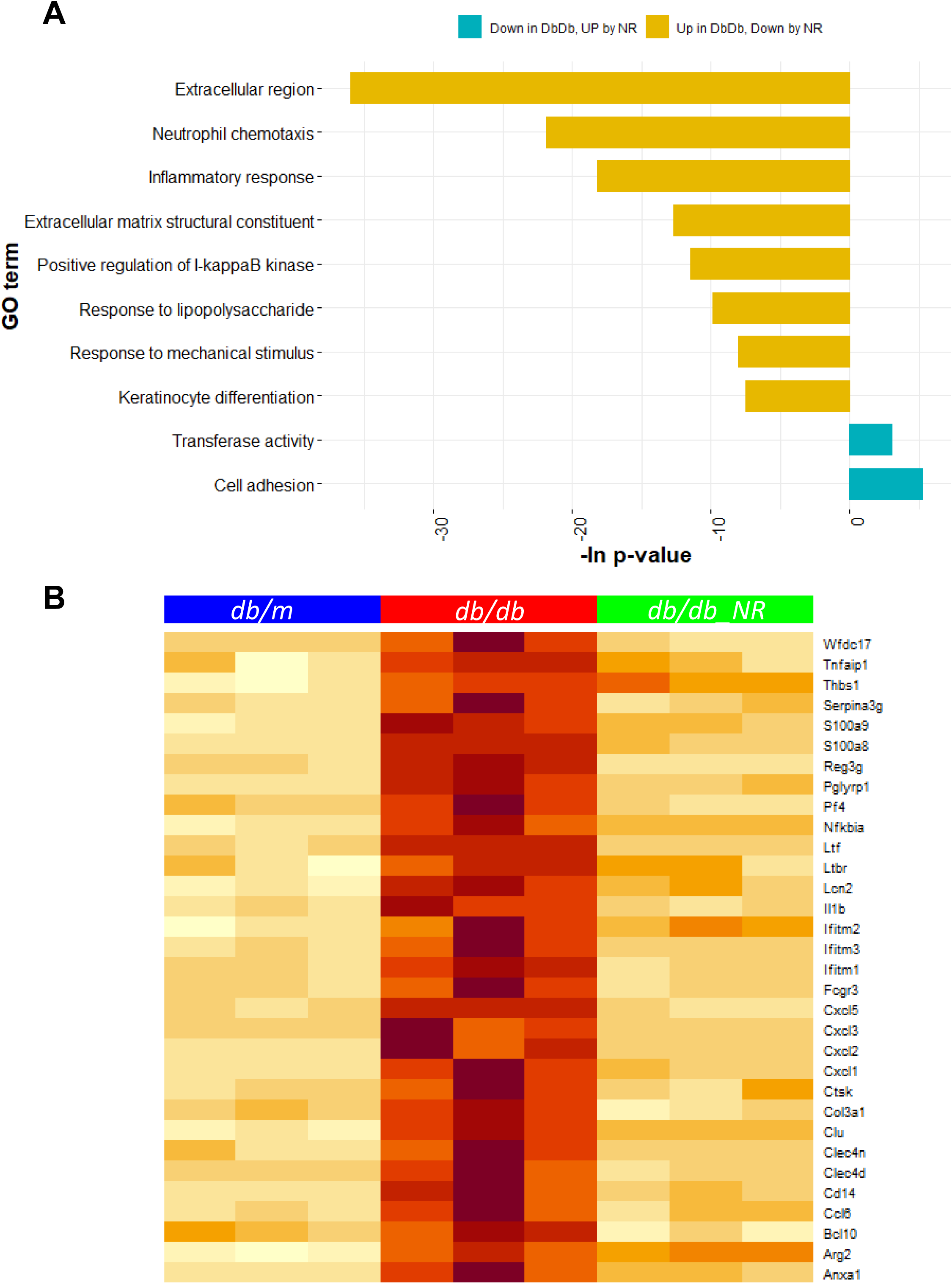
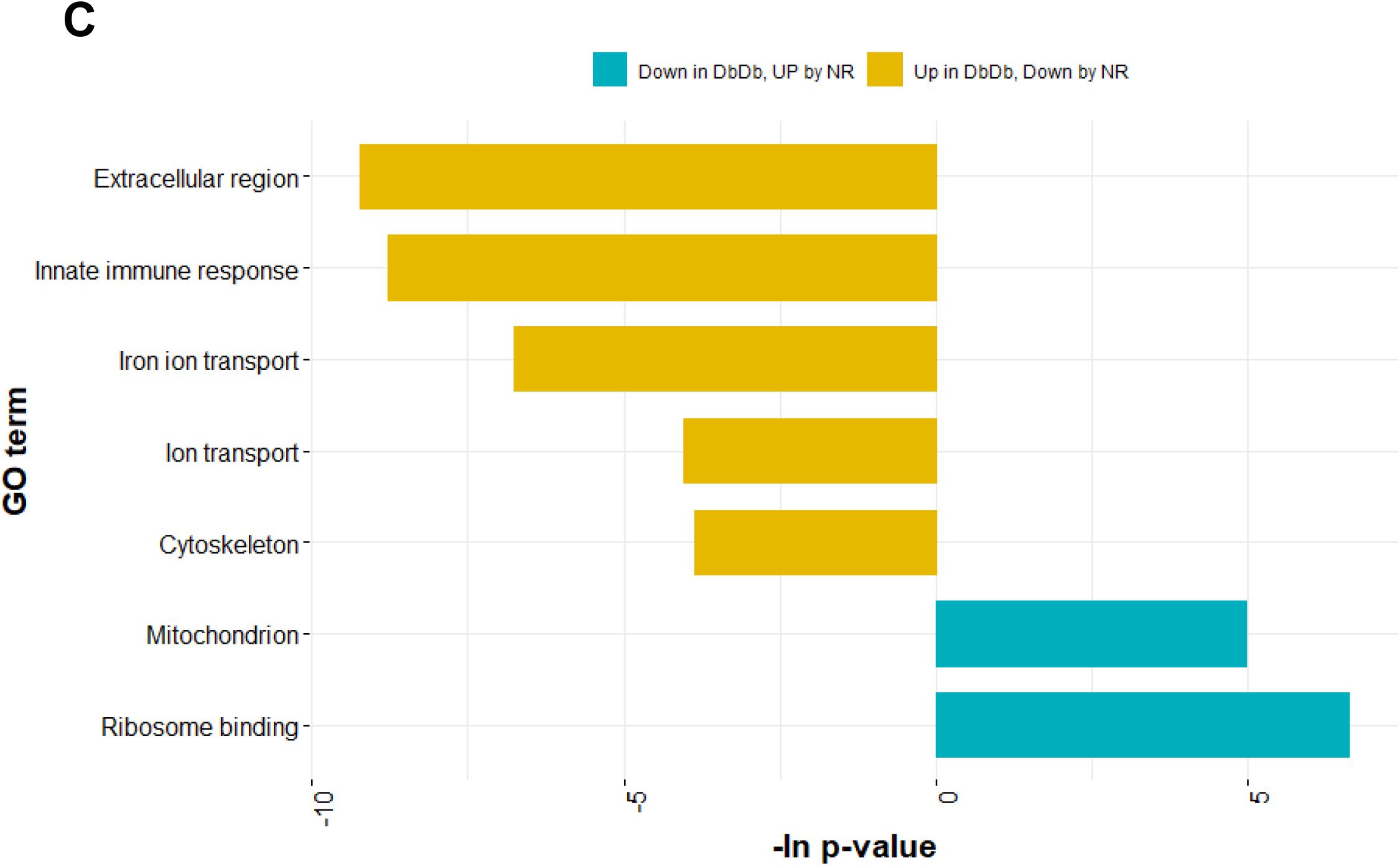
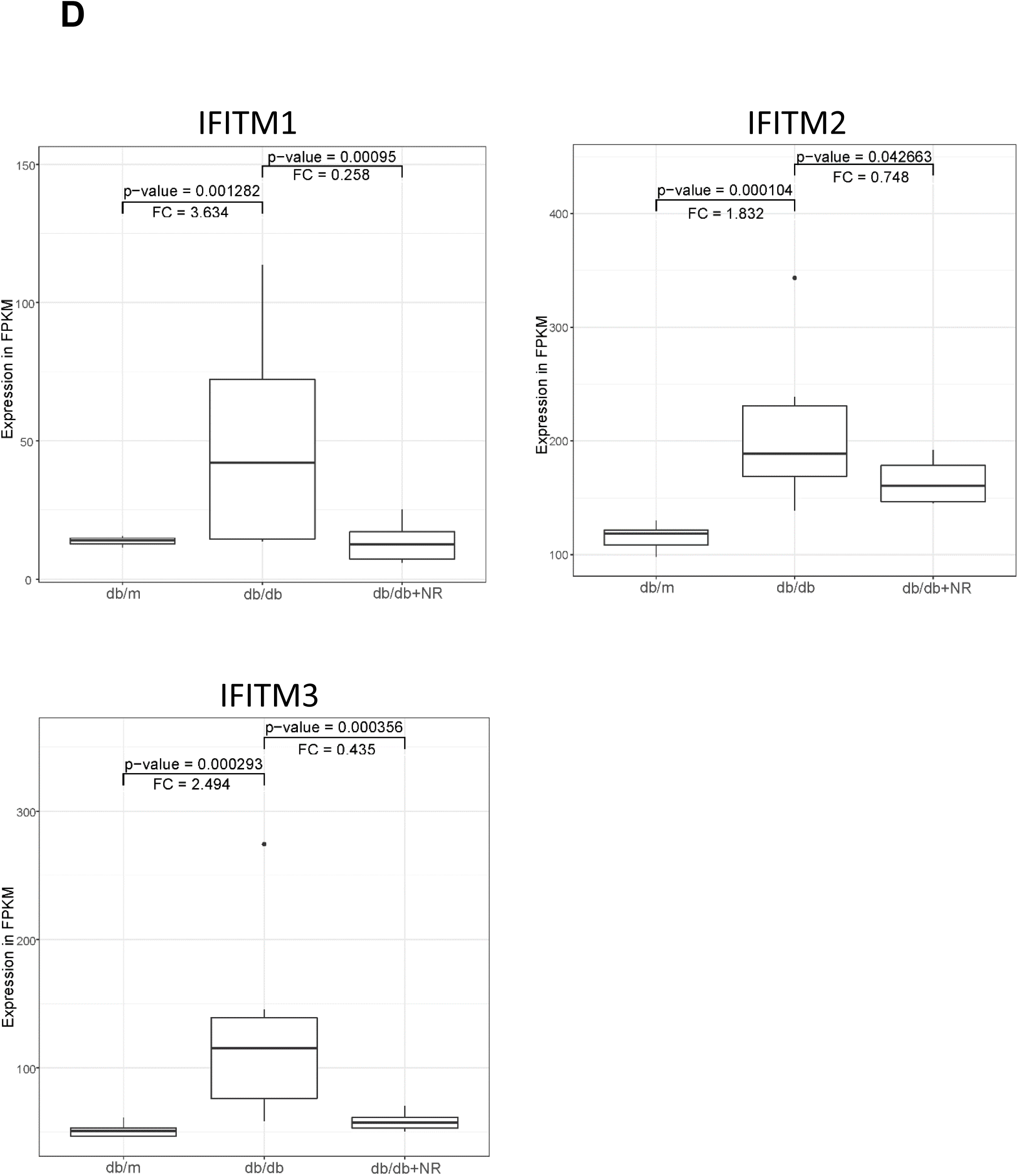
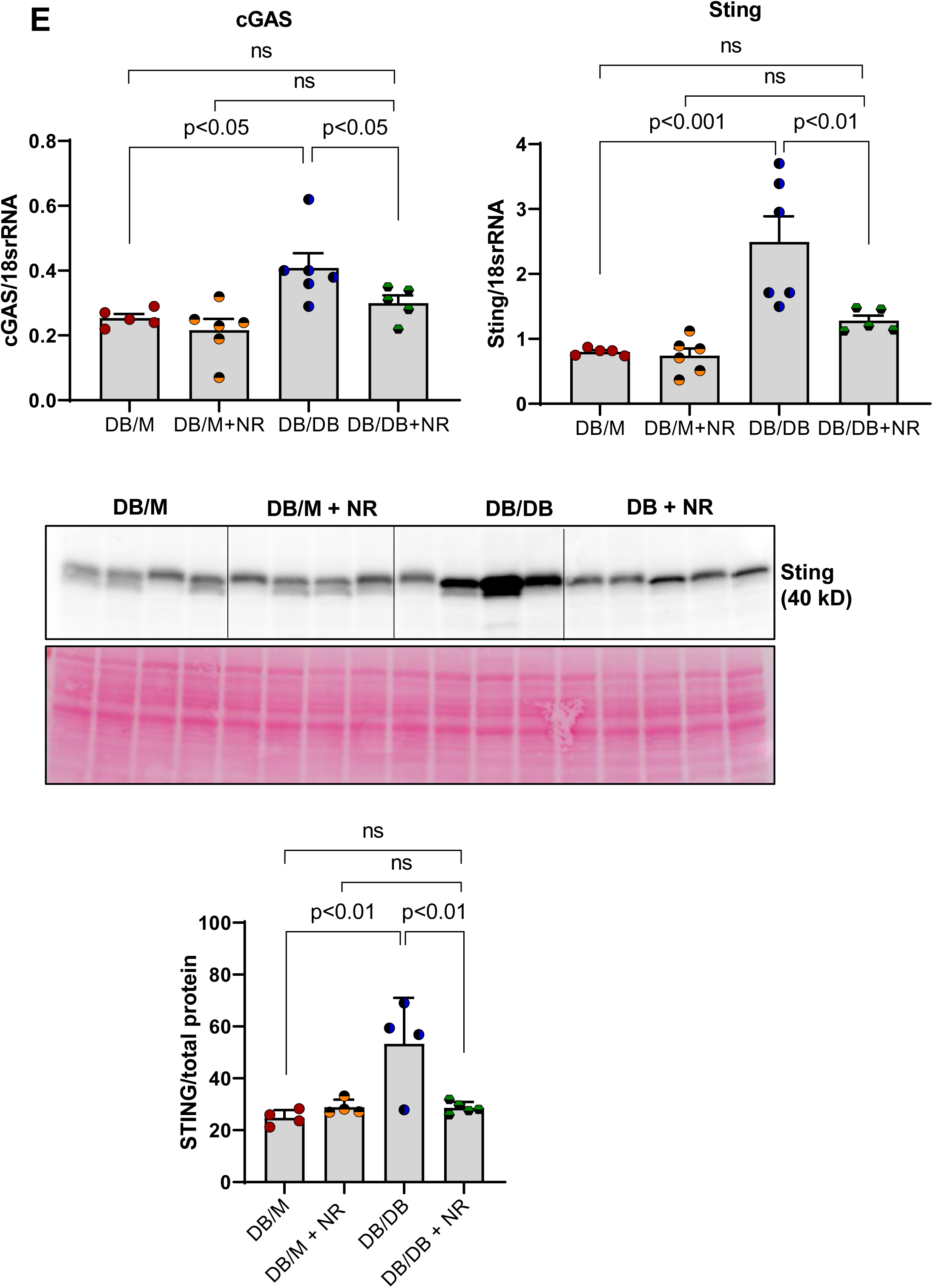
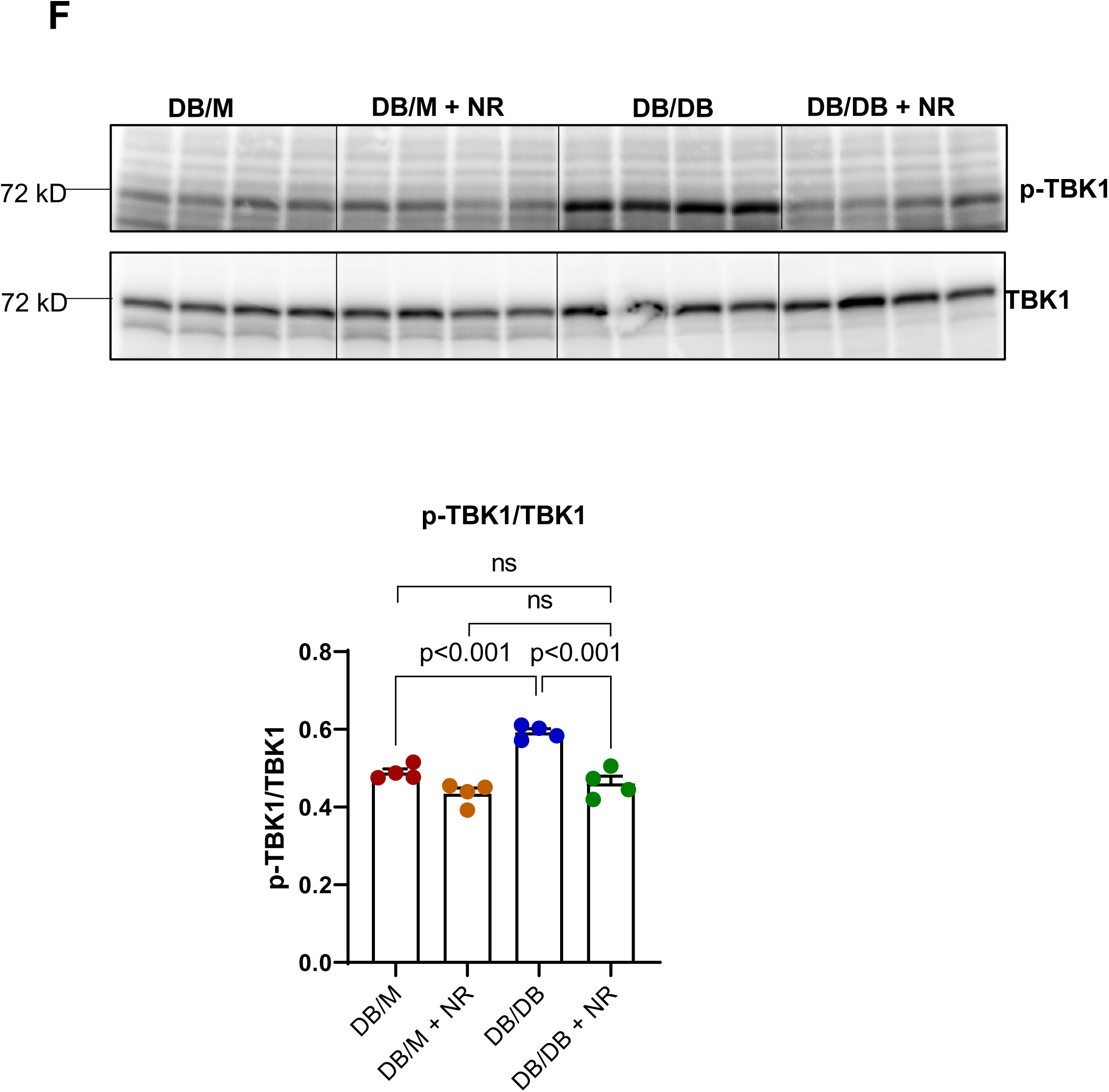
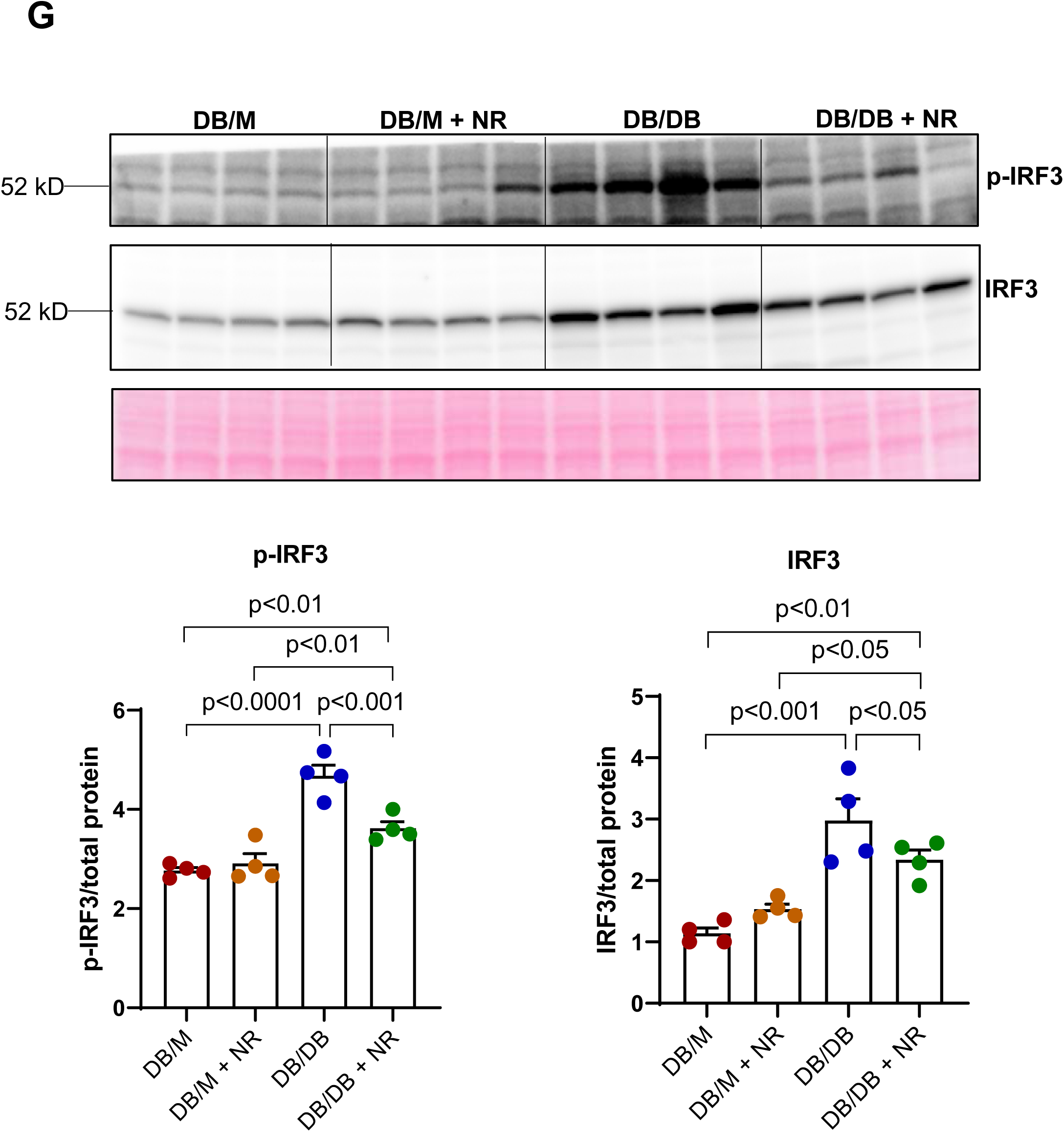
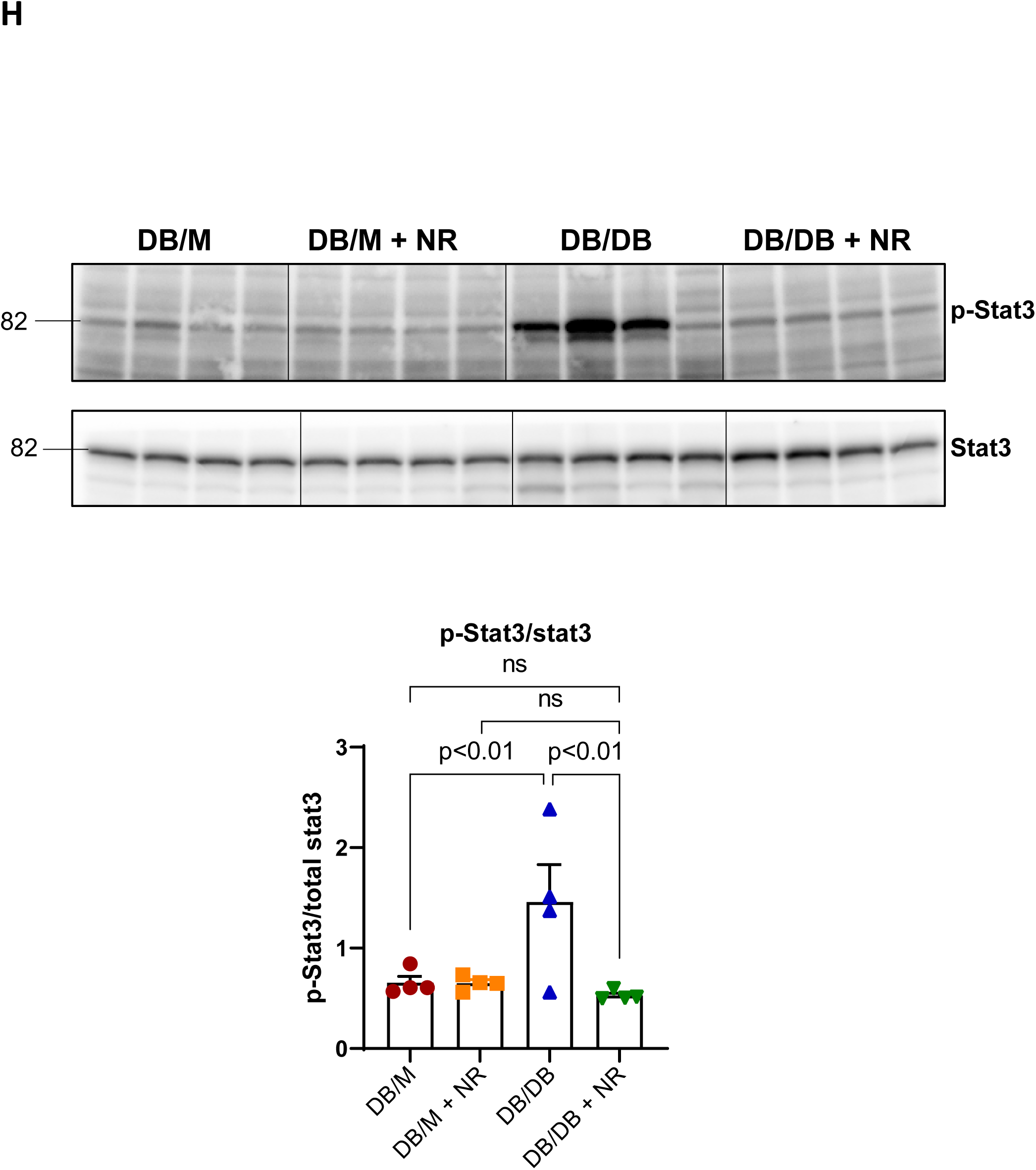
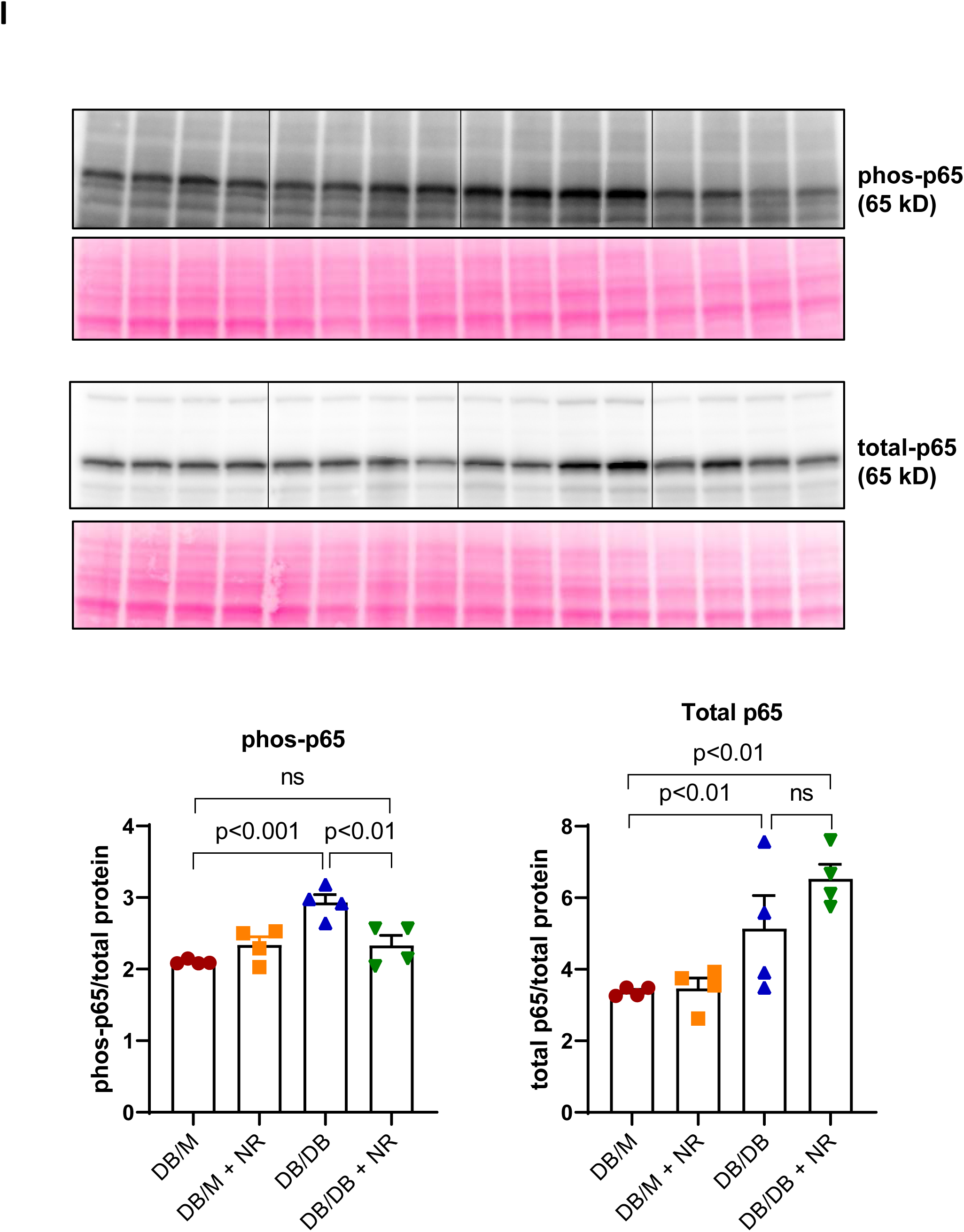
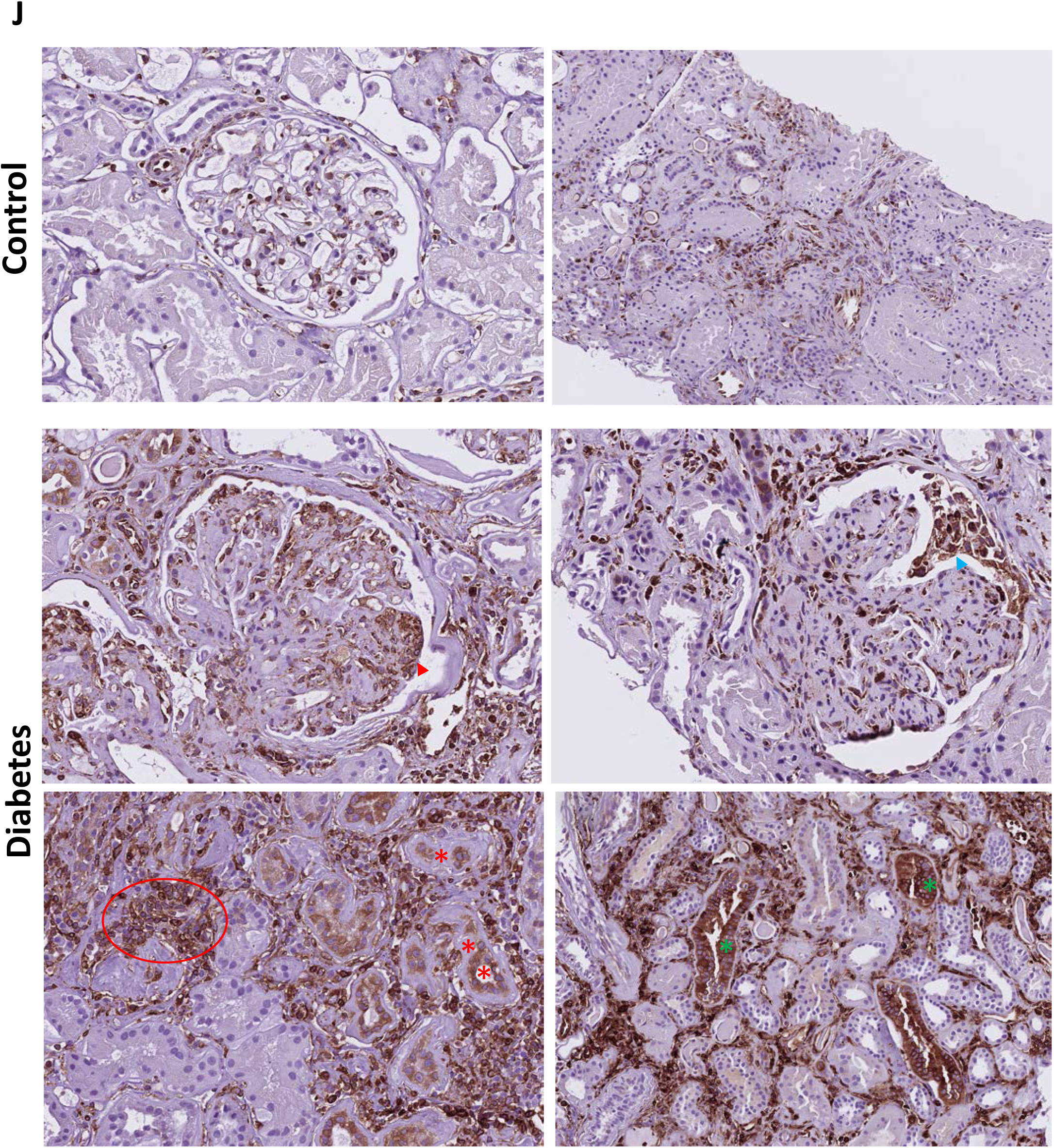
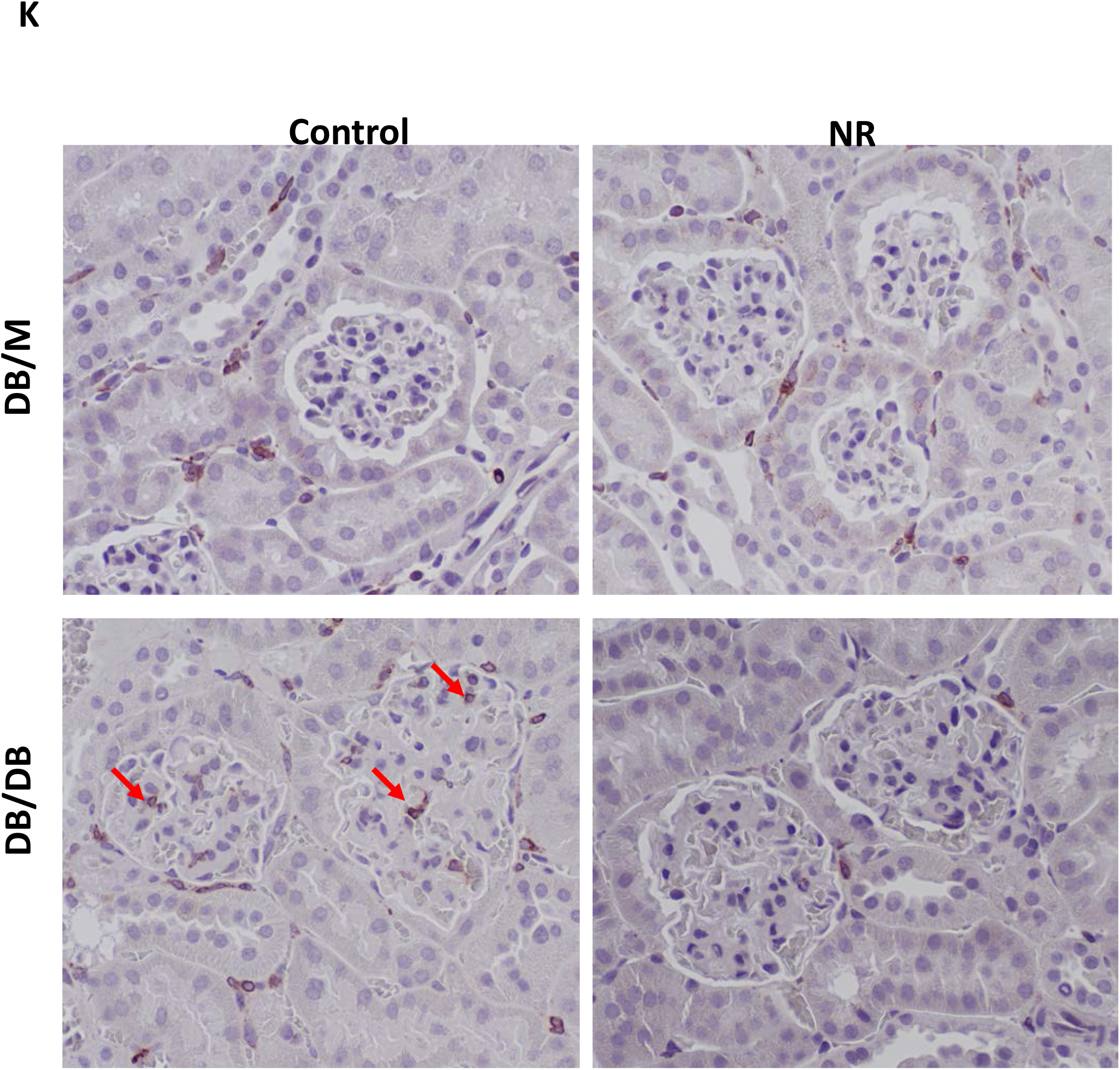
NR treatment inhibit the cGAS-STING activation in the kidney of db/db mice. A & B) Transcriptome and C) proteomics analysis of kidney indicating the upregulation of pathways related to immunity in the kidney of db/db mice compared to db/m controls. D) Transcriptomic analysis found IFITM genes which belong to interferon stimulated genes (ISG) was upregulated in db/db kidneys and inhibited by NR treatment. E) cGAS mRNA and Sting mRNA and proteins levels were increased in db/db mice and NR treatment decreased their levels. Downstream of cGAS-Sting singling proteins, active form of F) phosphorylated TBK1, G) phosphorylated IRF3, H) phosphorylated Stat3, I) phosphorylated p65 protein abundance were increased in db/db mice and normalized upon NR treatment, although total p65 levels were unchanged. Immunohistochemistry indicating J) Sting protein expression was increased in the kidney of humans with diabetes. K) Similar to humans, Sting expression levels were increased in glomeruli and interstitial kidney of db/db mice and reduced with NR treatment. The western blot images were quantified and normalized to the total protein or phosphorylated active form divided inactive total protein as loading control. n=5-6 per group, values presented as mean ± SEM with variance is calculated using one-way ANOVA.

To further evaluate STING activation in the kidneys, we performed the immunostaining of STING on kidney tissues. In non-diabetic human kidneys, STING expression is primarily limited to endothelial cells (glomerular, peritubular capillaries and larger vessels) and sparse interstitial inflammatory cells. By contrast, in diabetic renal parenchyma, an expanded pattern of STING staining is observed. (1) There is increased endovascular STING staining, enhanced inglomerular segments with prominent endothelium and endocapillary inflammatory cells including lymphocytes and monocyte/macrophages. (2) Prominent parietal epithelial cells showed increased STING expression. (3) By far, the most prominent compartment with STING expression are the increased influx of interstitial inflammatory cells. (4) Also noted is enhanced SITNG staining in atrophic tubule and tubules of the distal nephron (**Figure 4J**). the pattern of increased STING expression was noted, albeit not as broadly, and included primarily increased STING expression in intraglomerular inflammatory cells (**Figure 4K**). These data confirm the cellular compartments most notable for STING expression, with conservation of inflammatory cell STING expression in mouse and human.

### STING inhibition decreased inflammation in diabetic kidneys

To determine the role of the increased STING activity *per se* in mediating the inflammation in the diabetic kidney we treated db/m and db/db mice with the STING inhibitor C176 ^31–33^ **(Figure 5A)**. Treatment of db/db mice with C176 prevented the increases in IRF3 and phospho-IRF3 **(Figure 5B)**. C176 also prevented the increases in the inflammatory markers phospho-STAT3 protein and IL-1β mRNA levels **(Figure 5C)**.

**Figure 5:**
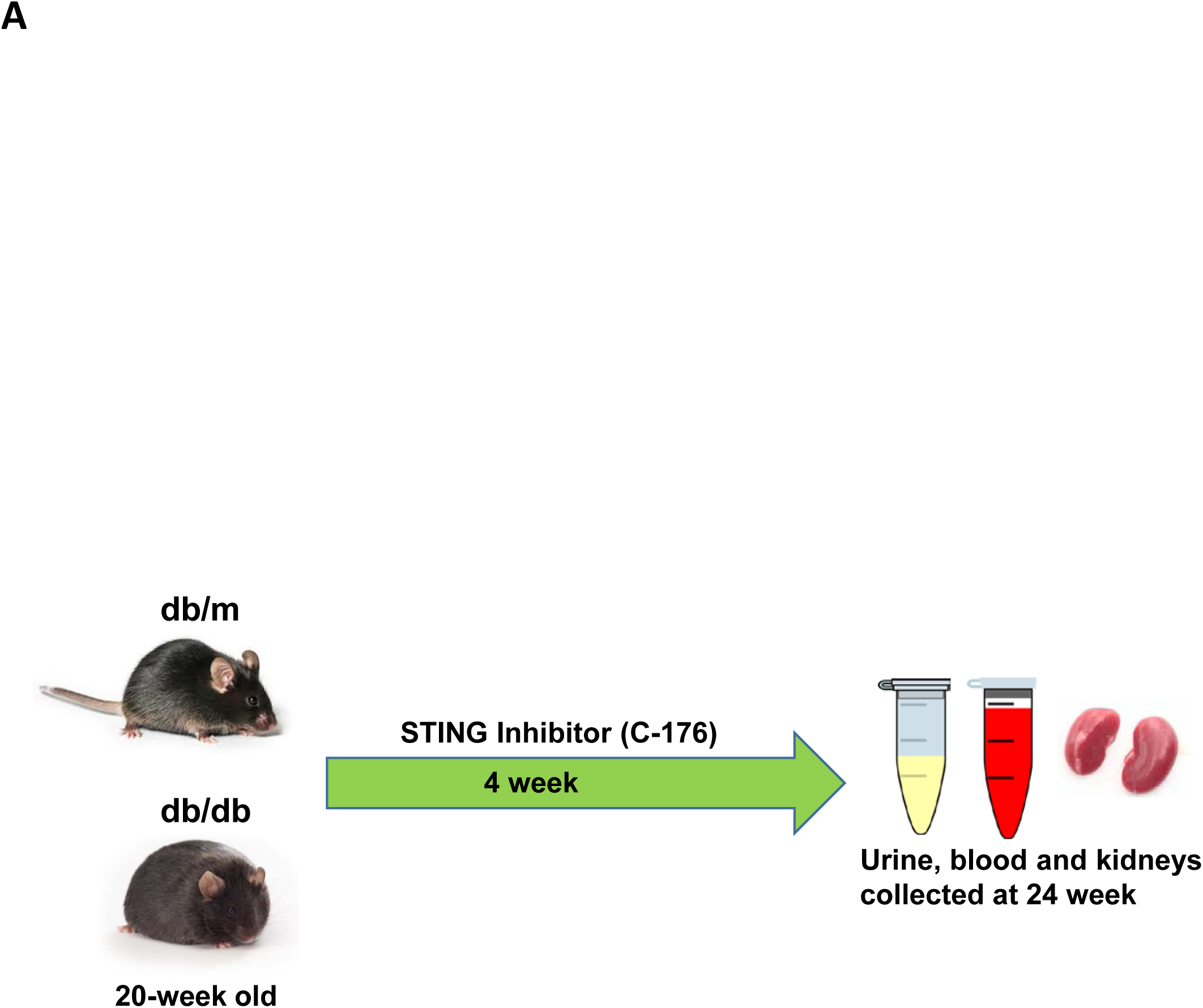
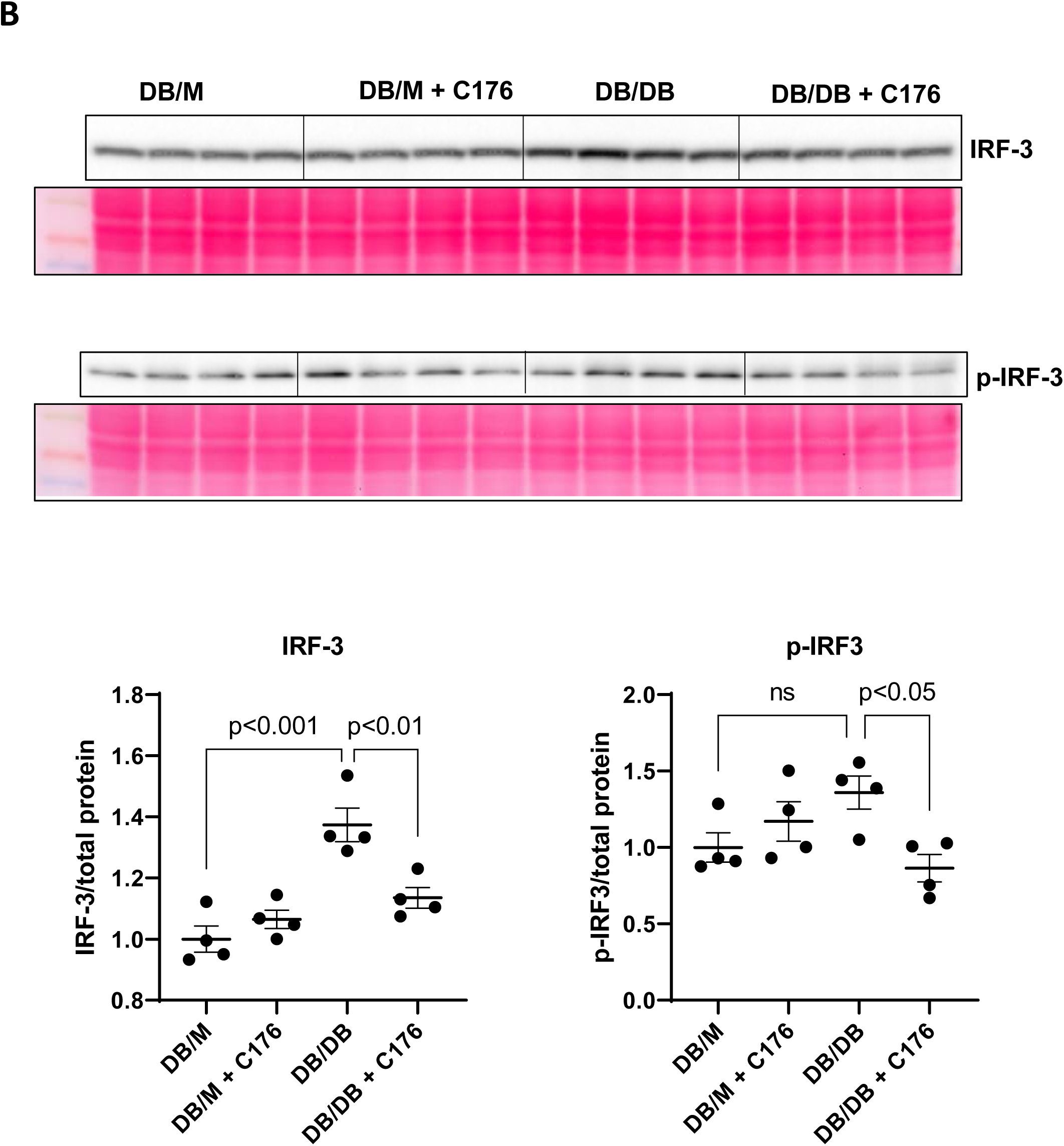
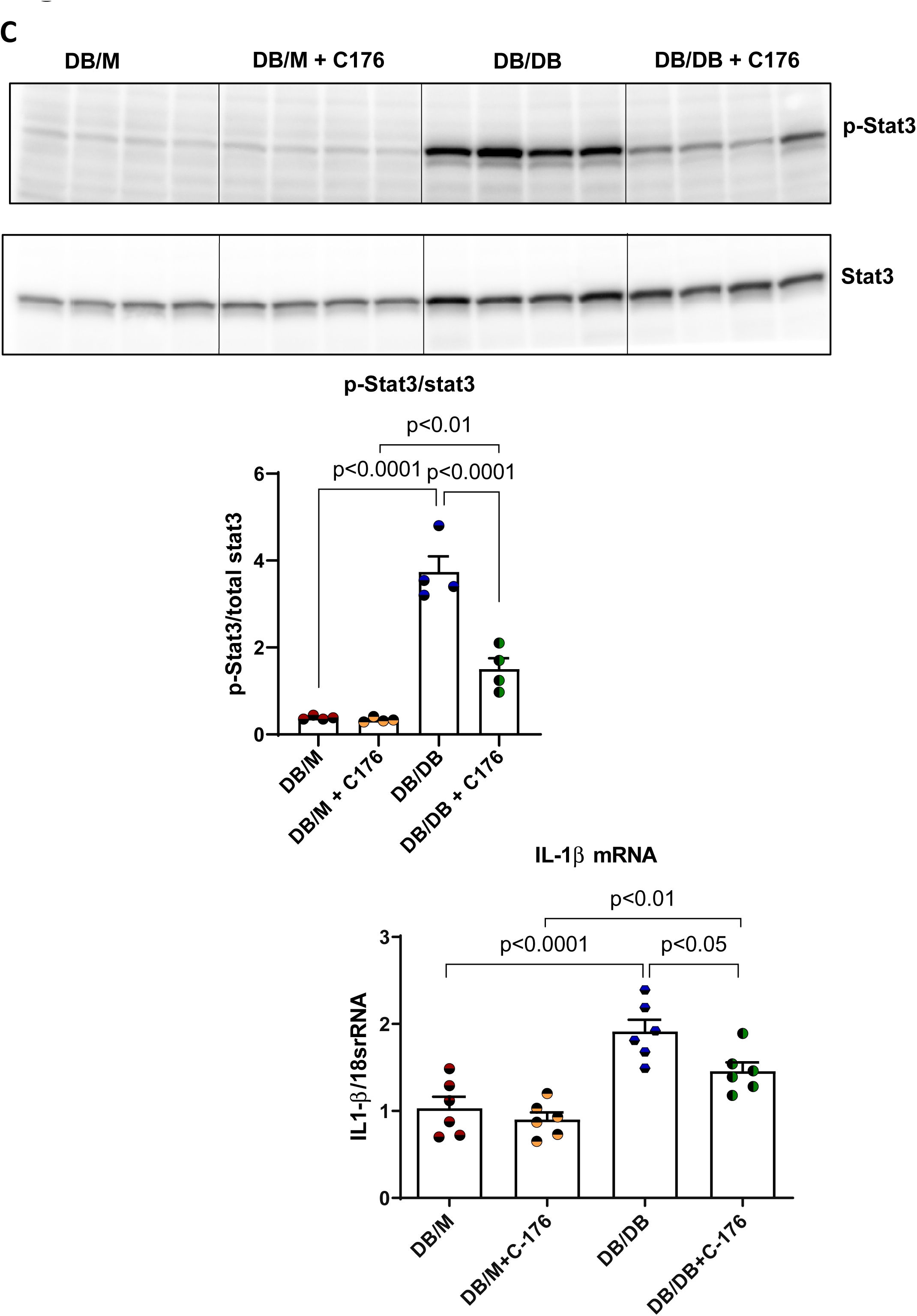
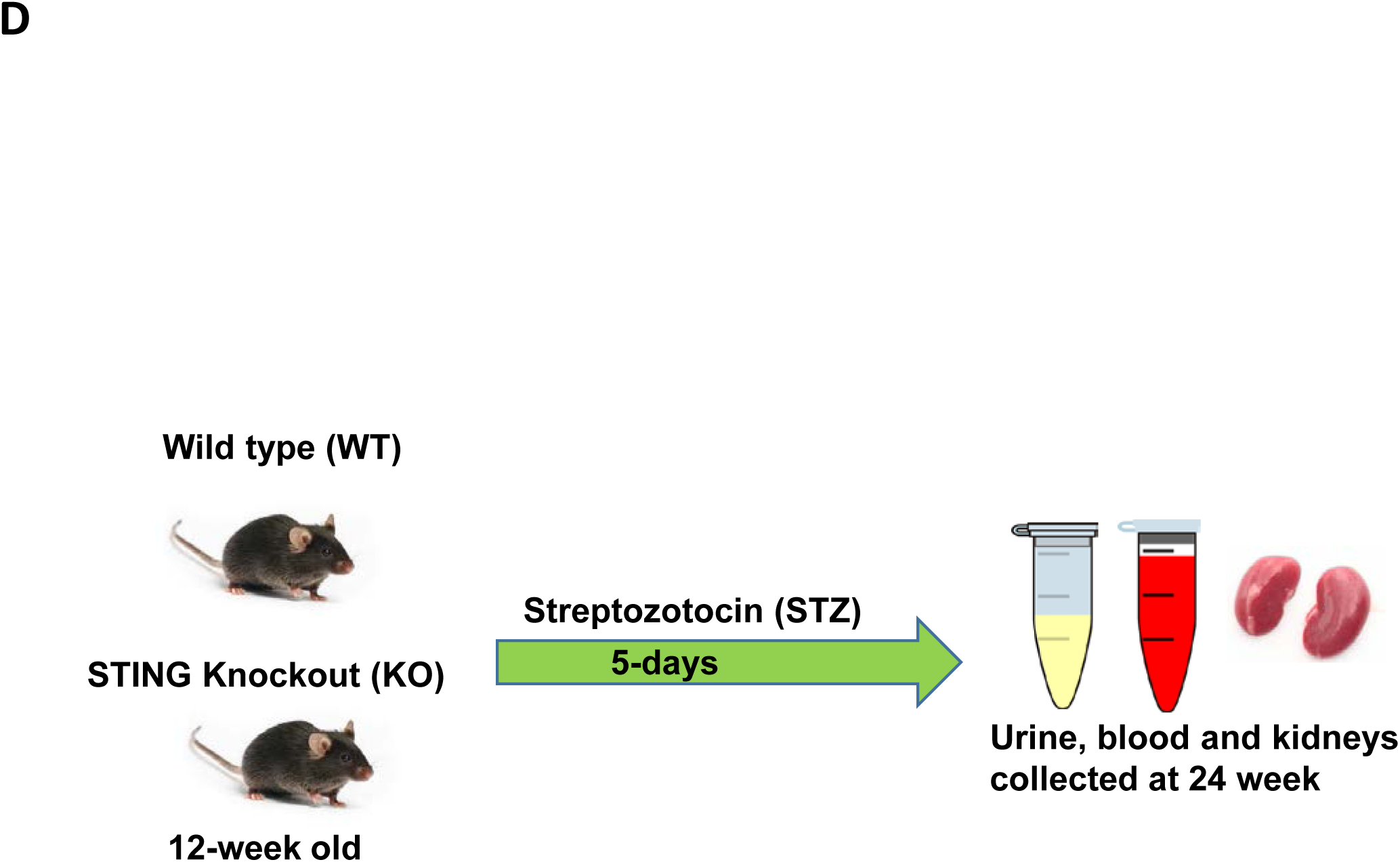
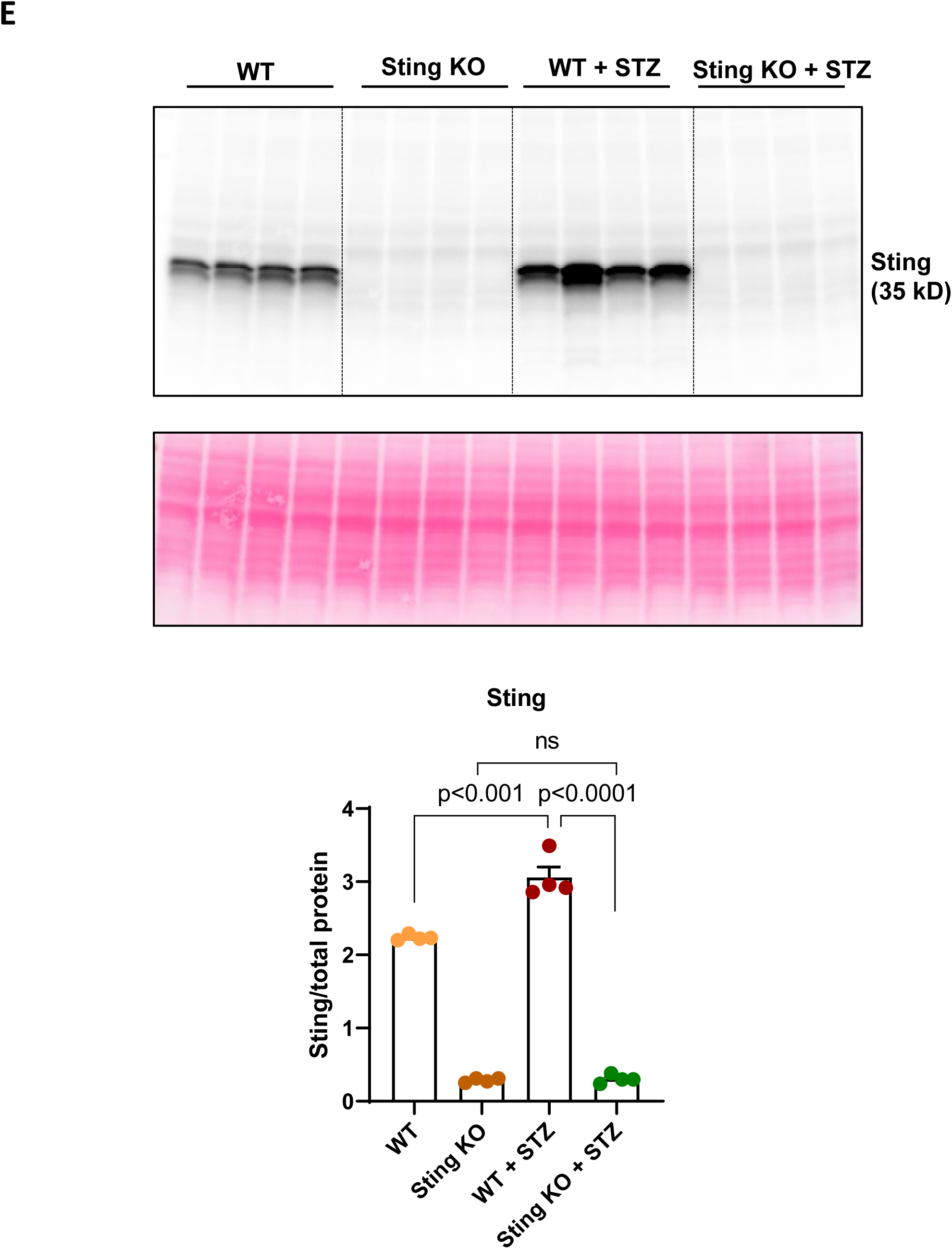
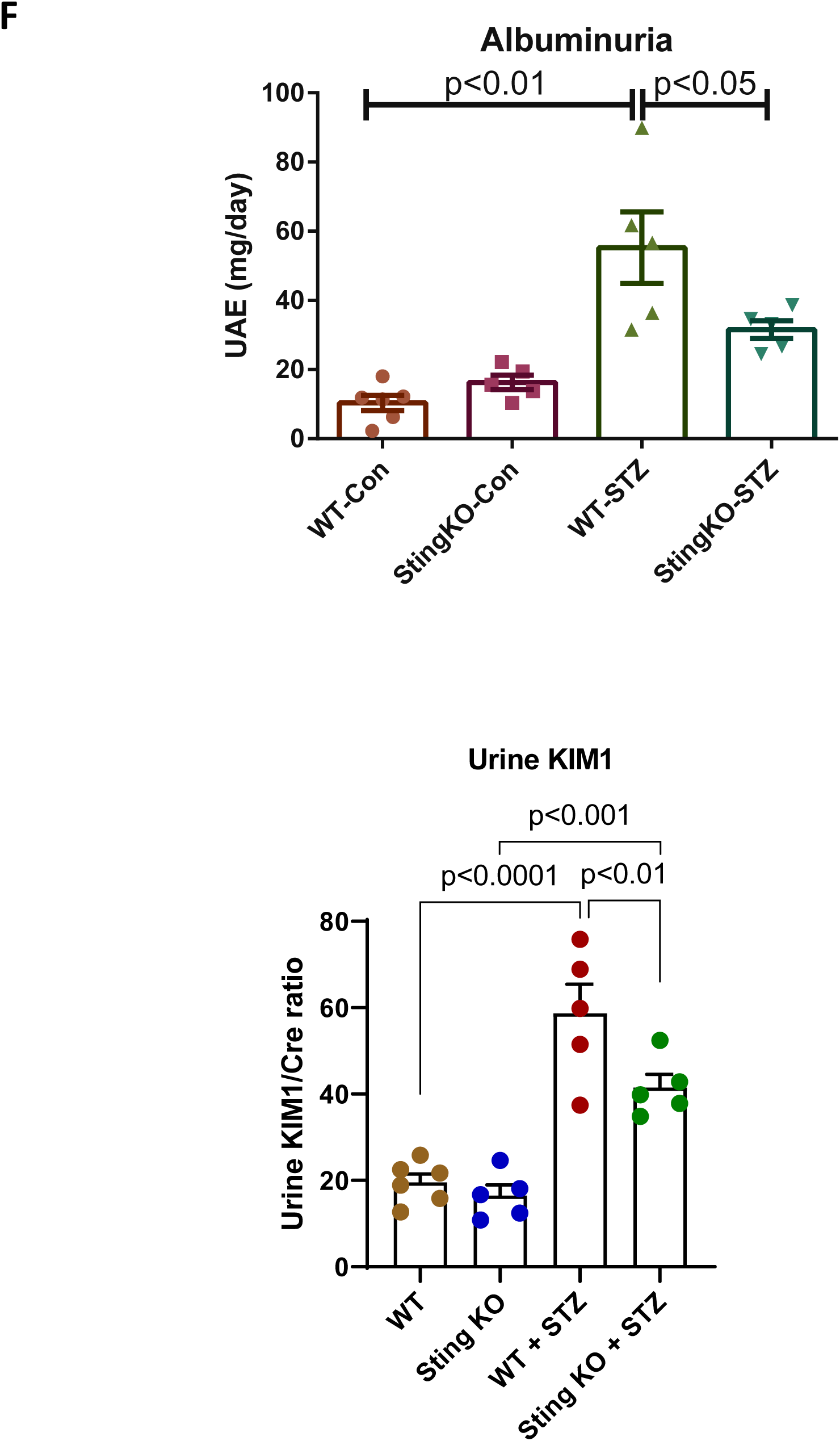
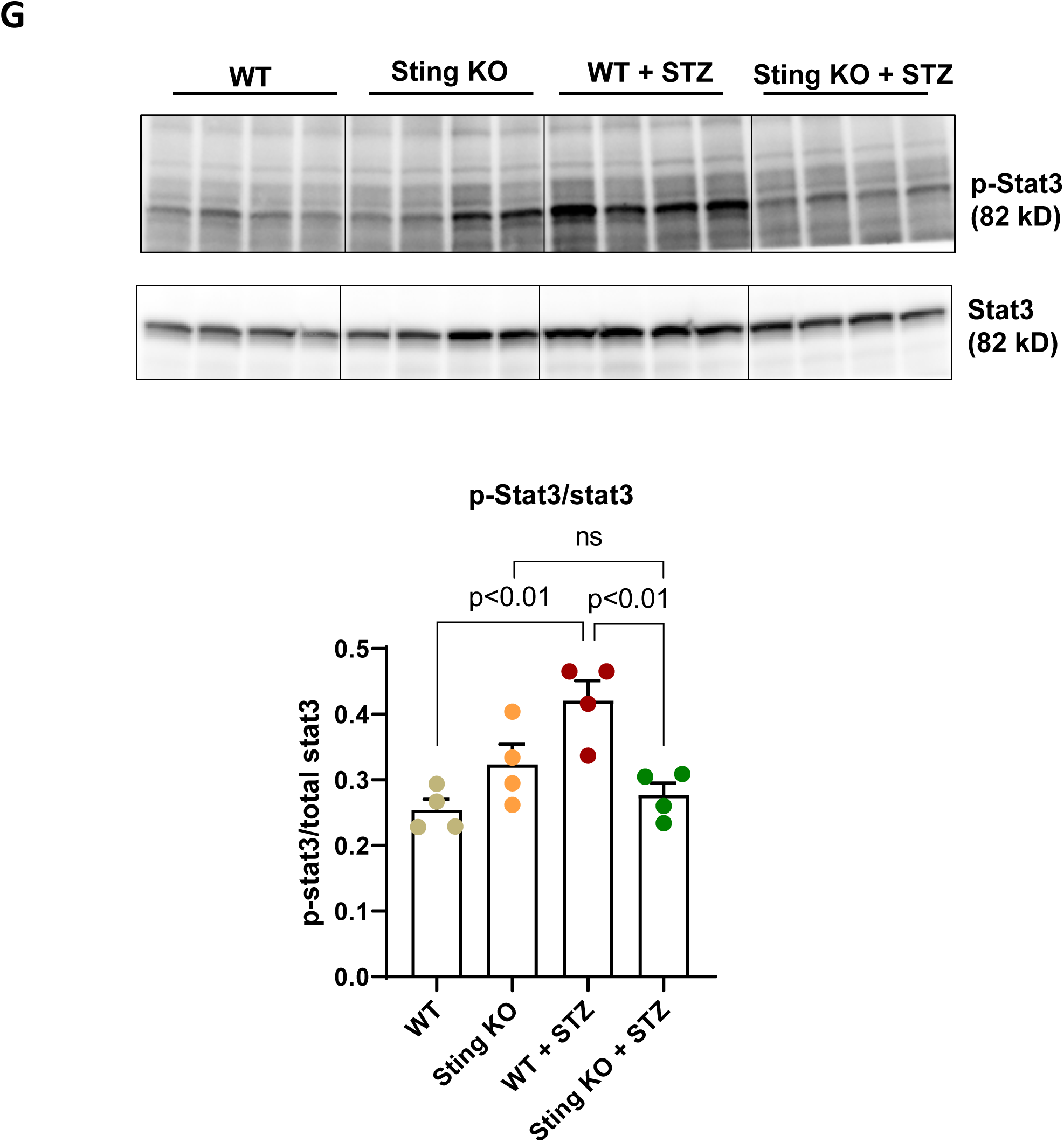
Effects of STING inhibitor C176 or STING knockout (KO) on the inflammatory response and kidney function in db/db and streptozotocin (STZ) induced diabetic mice. A) Experimental scheme for the study with C176 treatment. B) total and phosphorylated IRF3 protein levels were decreased significantly in db/db mice treated with C176. C) phosphorylated Stat3 protein levels were higher in db/db mice and C176 treatment decreased their levels. IL1-β mRNA levels increased in kidney and decreased with C176 treatment in db/db compared to control db/m mice. D) Experimental scheme for the study with STING KO mice. E) Sting protein levels were significantly increased in wild type (WT) mice with STZ induced diabetes and markedly decreased in Sting KO mice made diabetic with STZ. F) Urinary albumin/creatinine and urinary KIM1/creatinine ratio were increased significantly in wild type mice with STZ-induced diabetes and they were markedly reduced in the Sting KO mice made diabetic with STZ. G) Phosphorylation of stat3 protein levels were higher in kidney of WT mice with STZ-induced diabetes and normalized in STZ induced diabetic Sting KO mice. n=4-6 per group, values presented as mean ± SEM with variance is calculated using one-way ANOVA.

To determine a direct role for STING, independent of the potential off-target effects of C176, we performed studies with the STING KO mice **(Figure 5D)**. In these studies, wild type and STING KO mice were made diabetic with the administration of streptozotocin (STZ). In the STZ mice there was a significant increase in STING protein level, which was prevented in the STING KO mice made diabetic with STZ **(Figure 5E)**. STZ induced increases in urinary albumin and urinary KIM-1 which were significantly attenuated in STING KO mice **(Figure 5F)**. In the STZ mice there was increased protein abundance of phospho-Stat3 which was also significantly decreased in STING KO mice **(Figure 5G)**.

### Inflammation was associated with NAD+ reduction and mitochondrial dysfunction

To evaluate the injury mediated by inflammation *per se*, we used a LPS-induced acute kidney injury model. While LPS activated innate immune response, a cascade of downstream signaling triggered increased STING expression and decreased mitochondrial complex I activity. Other factors involved in the mitochondrial homeostasis were also found to be decreased in LPS kidneys, such as NAD+ level, and expression of PGC-1a, estrogen-receptor related protein-α (ERRα) and sirtuin (SIRT) 3 (**Supplementary Figure S3**).

### NR treatment increased NAD+ level and increased SIRT3 expression and activity

The complexity of interconnecting crosstalk between inflammatory response and mitochondria dysfunction prompted us to further assess the effects of NR treatment in the kidneys. We measured NAD+ levels in db/m and db/db kidneys. Although there was no change in baseline NAD+ levels between db/m and db/db kidneys, NR significantly increased NAD+ levels in both db/m and db/db kidneys (**Figure 6A**).

**Figure 6:**
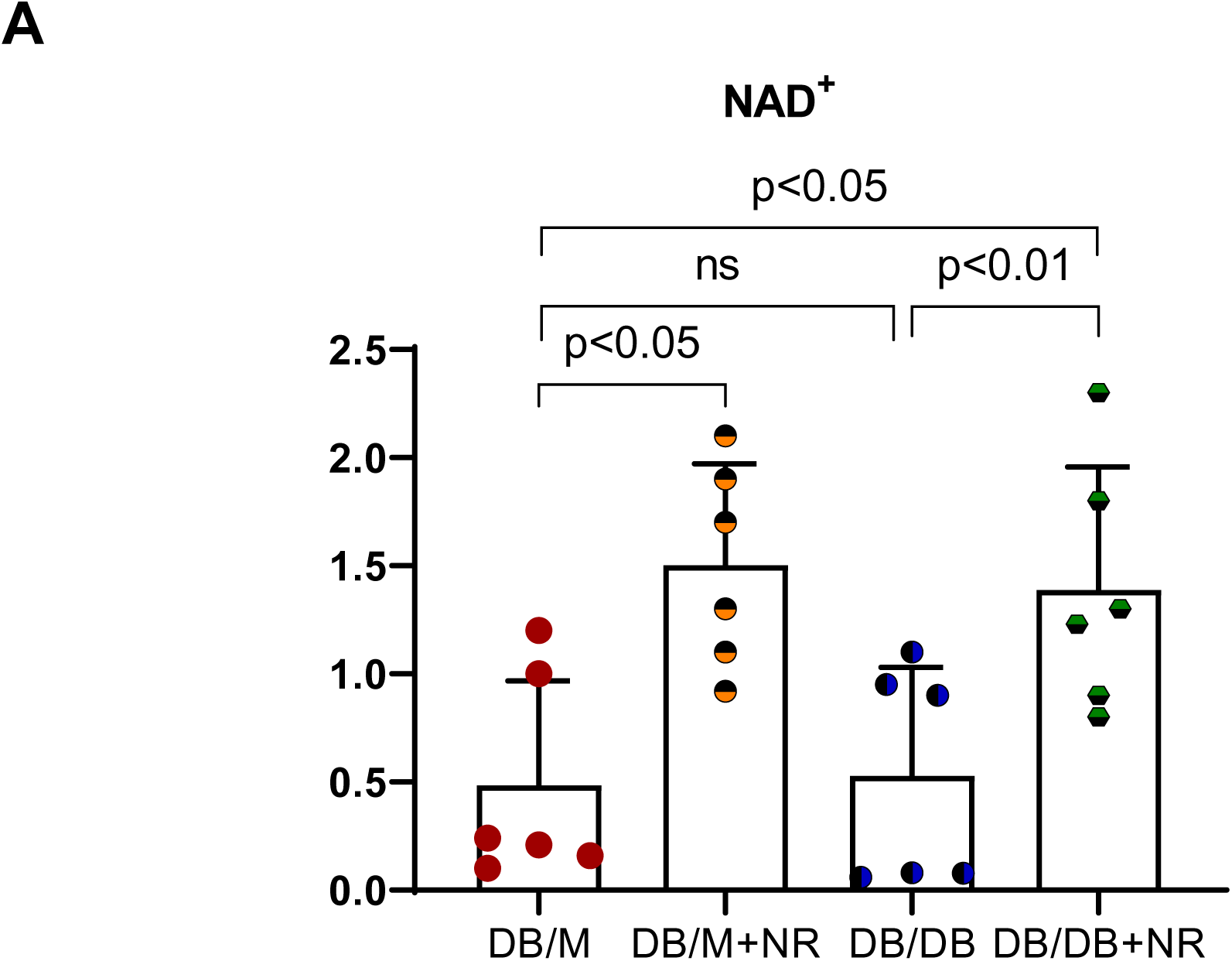
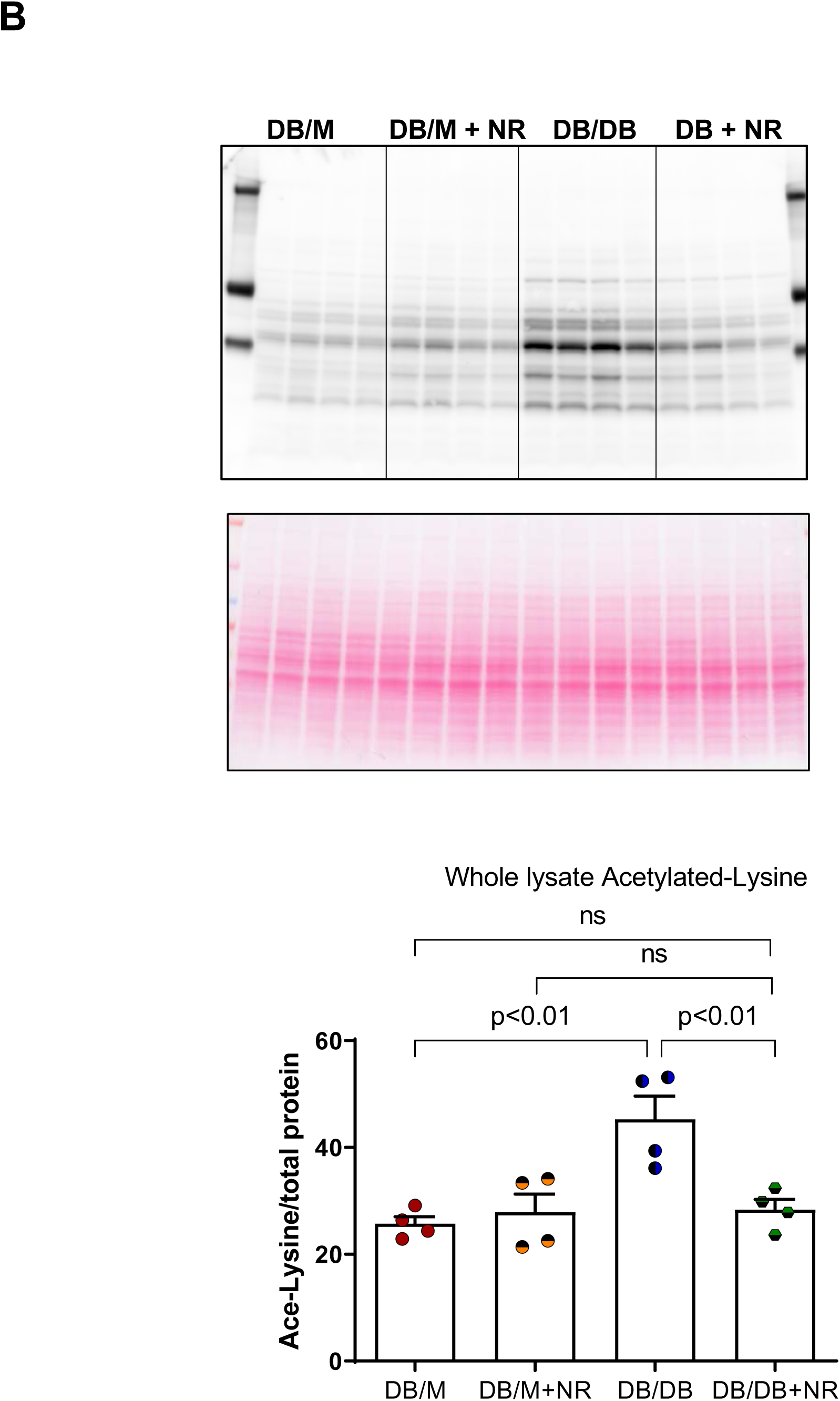
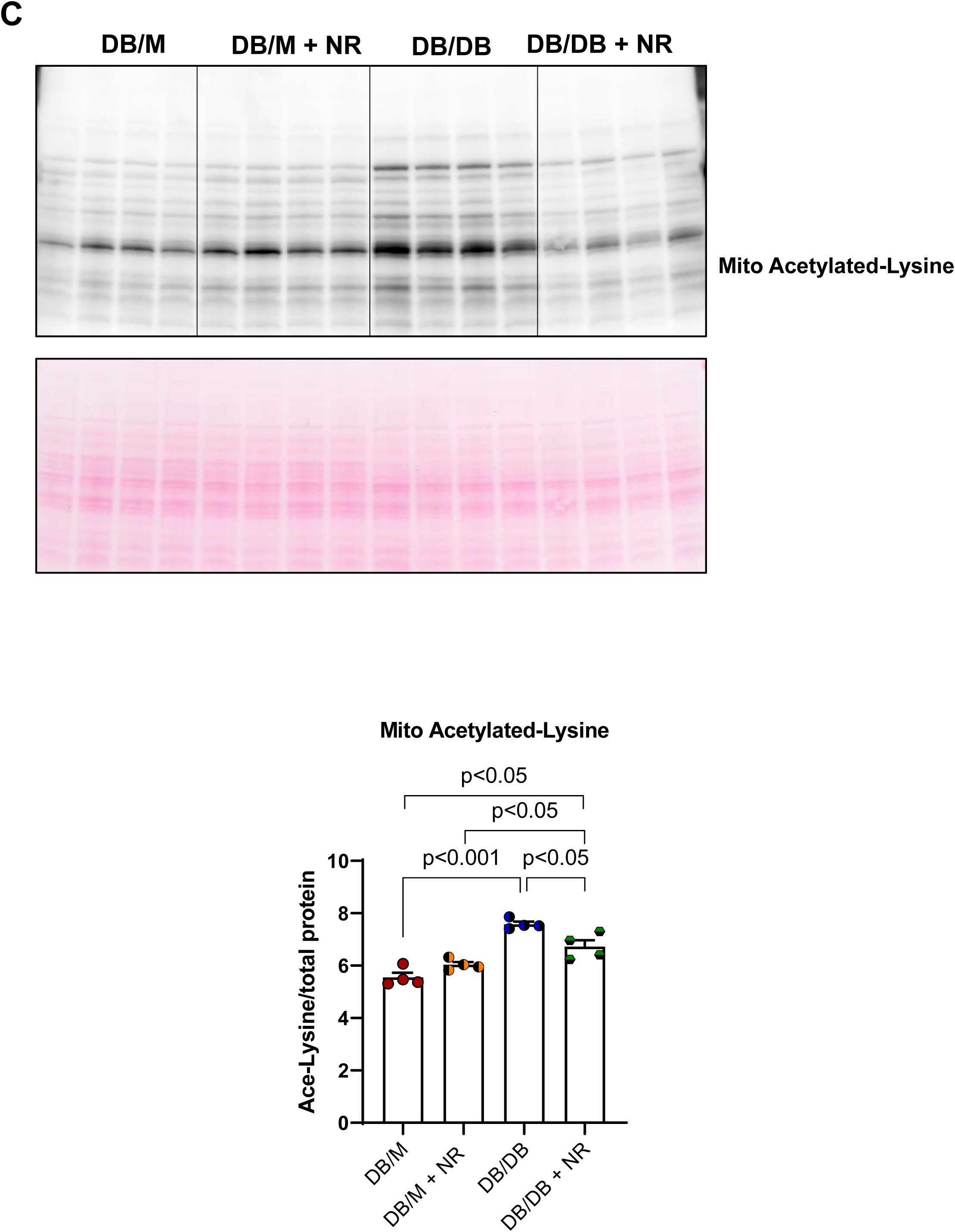
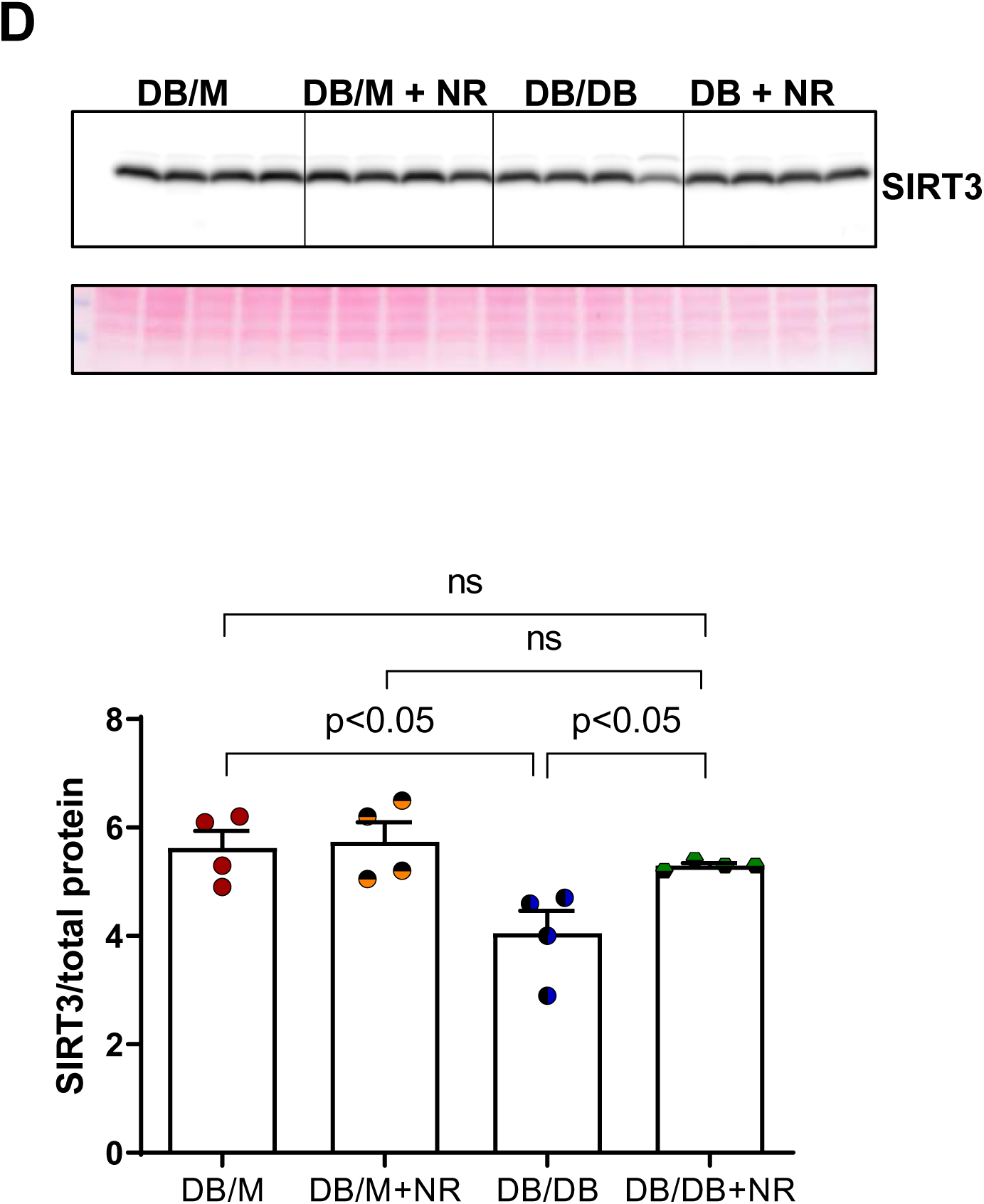
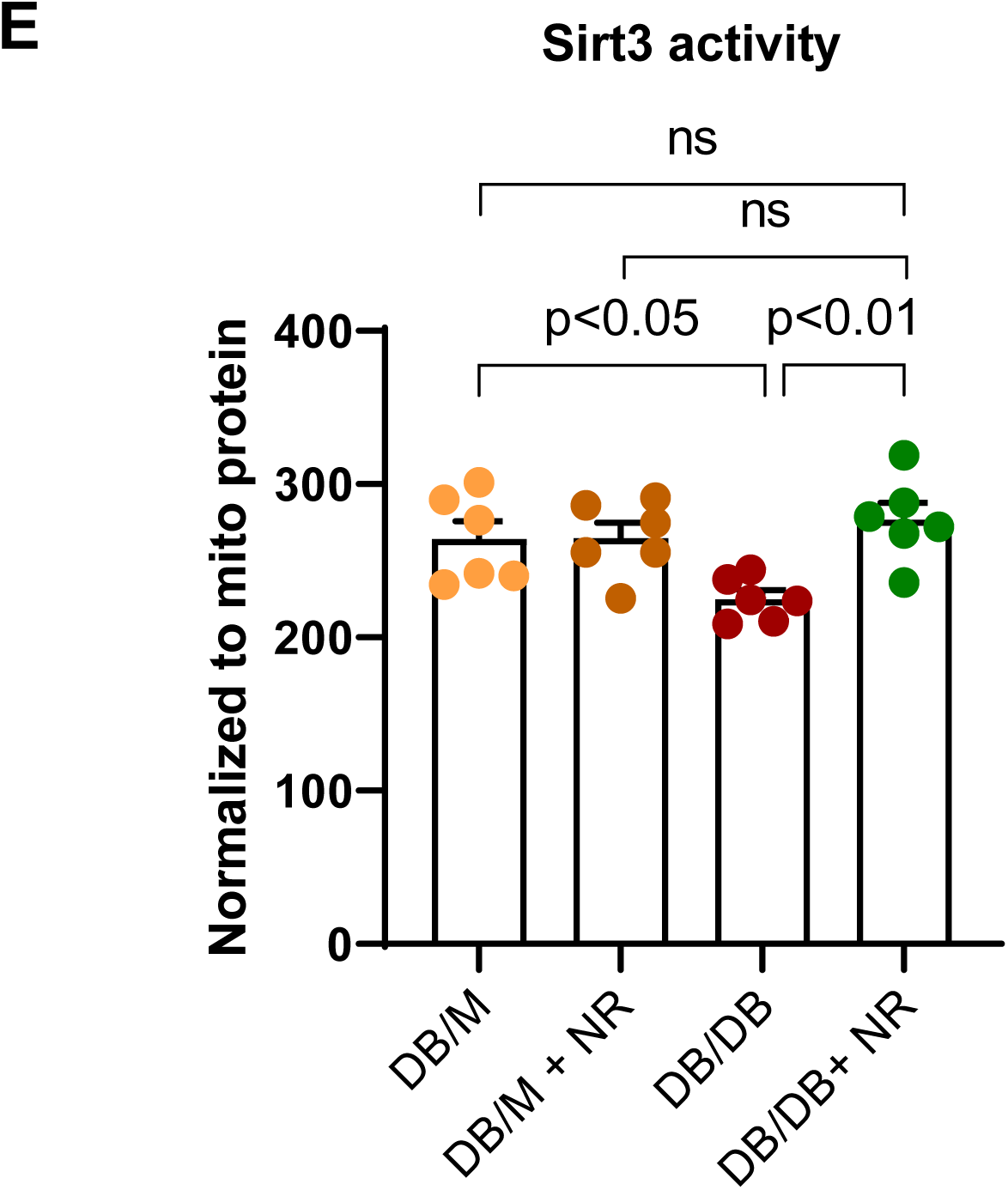
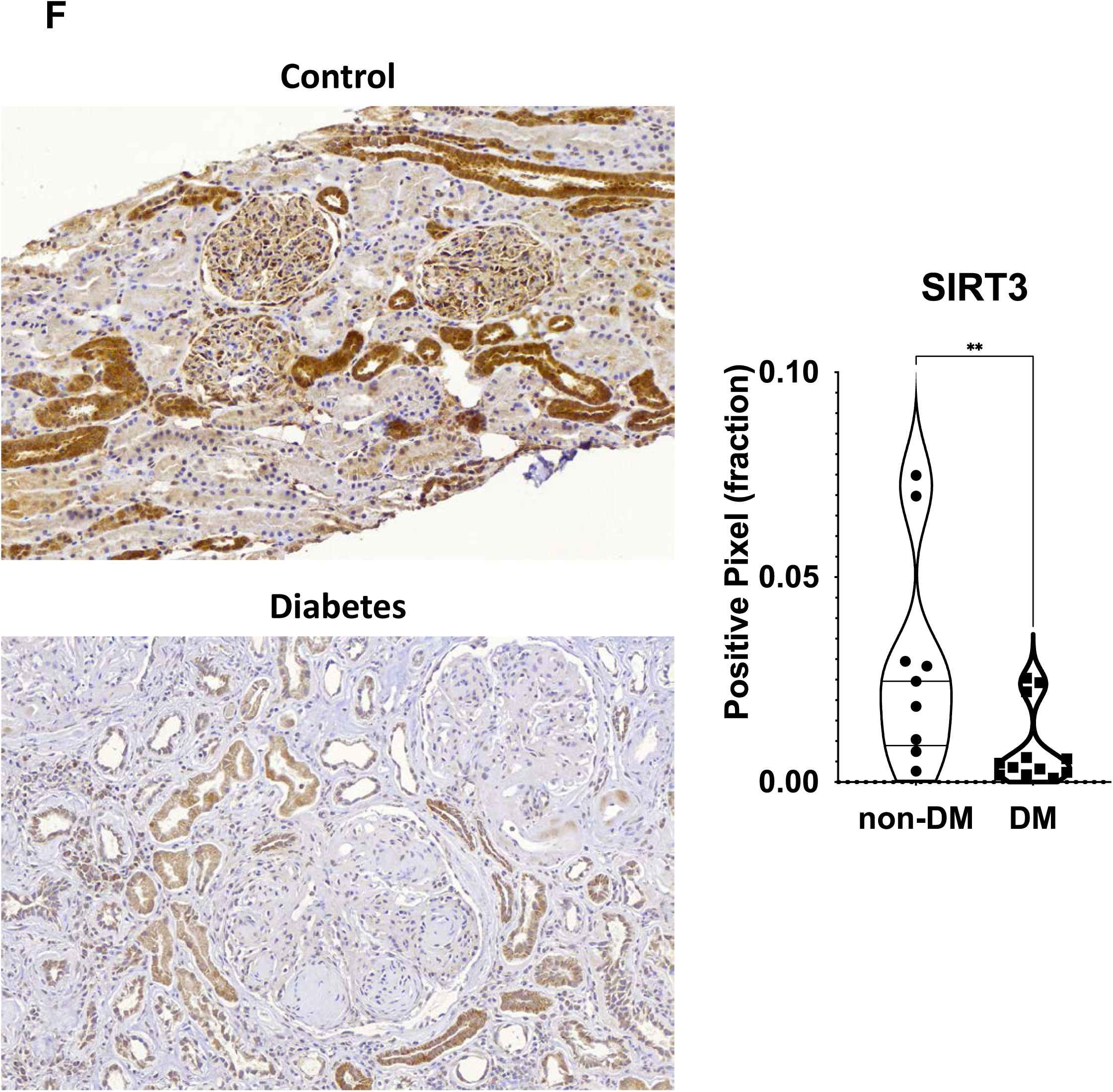
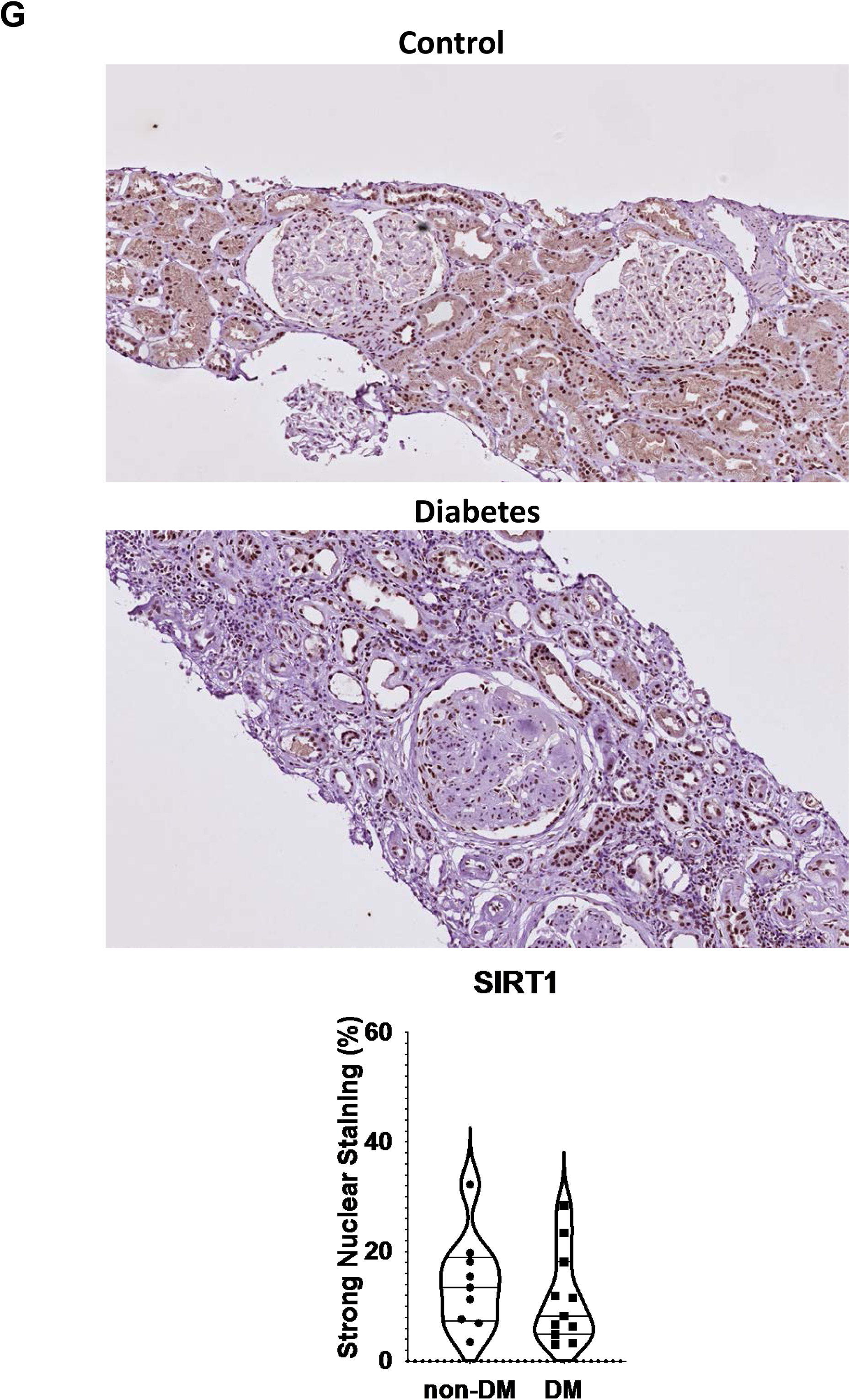
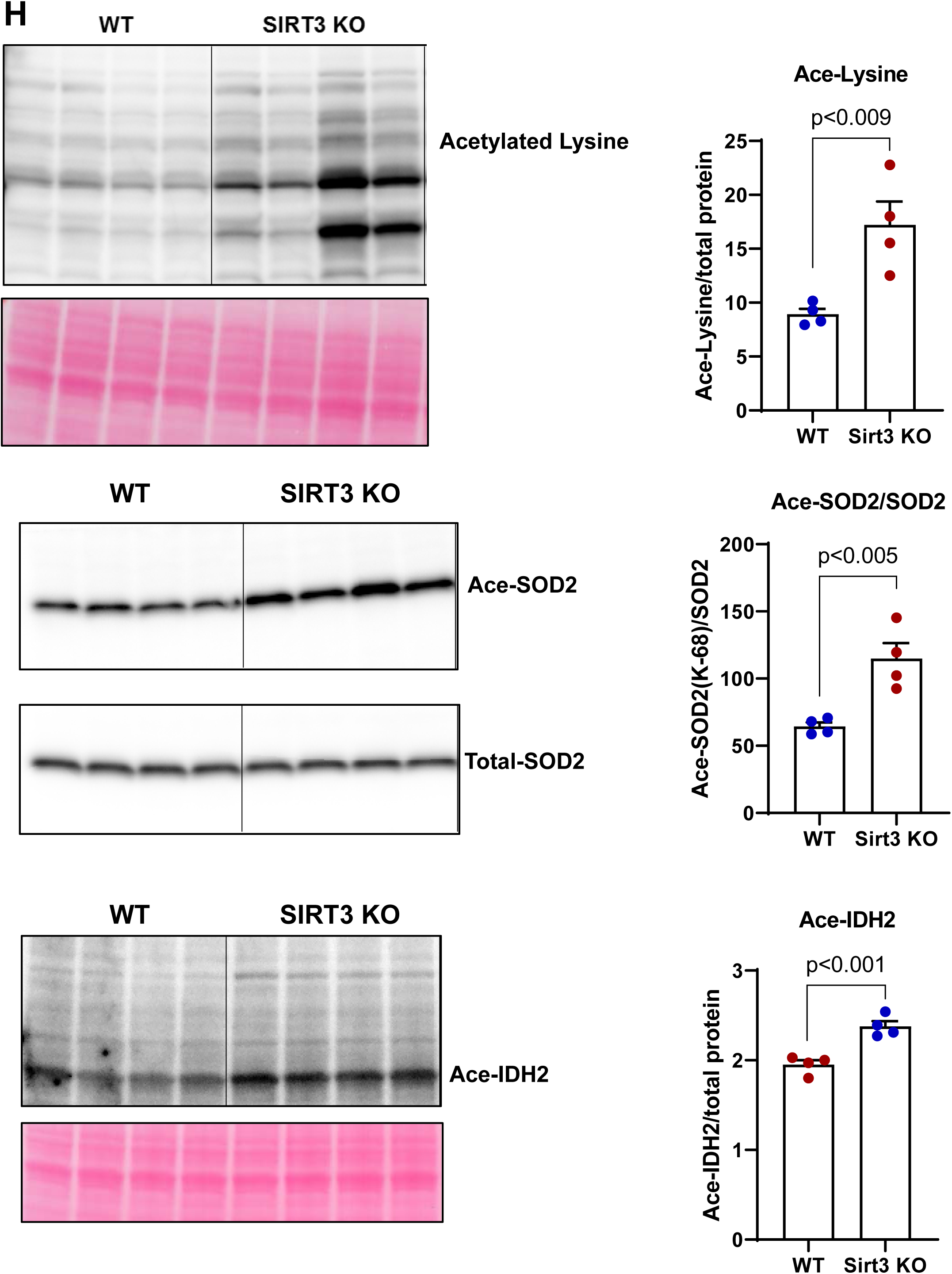
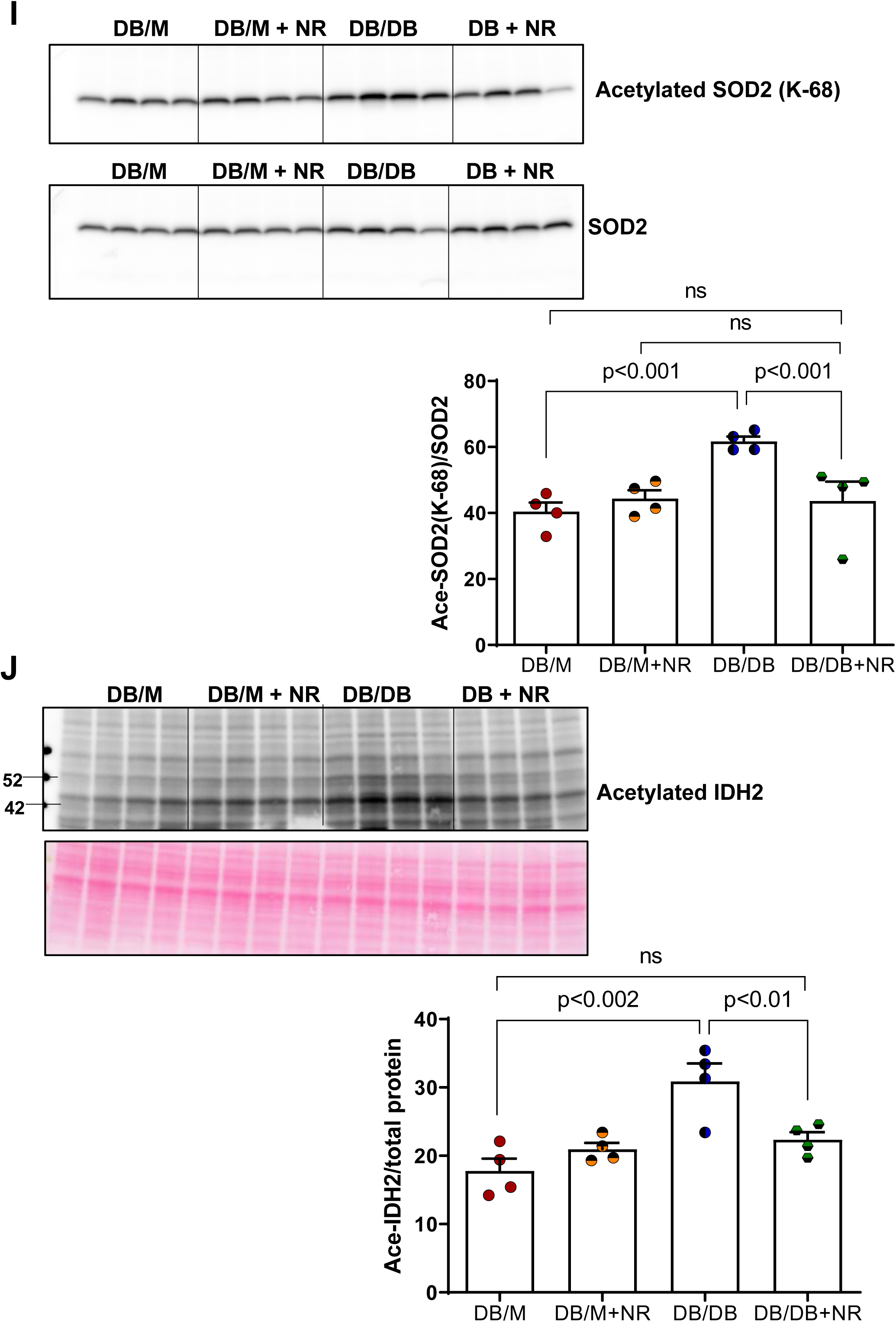
Effect of NR treatment on NAD^+^ levels, SIRT3 expression and activity in the kidney as well as expression of total acetylated proteins in the kidney of SIRT3 KO mice. A) NAD^+^ levels were measured in kidney and comparable in both db/m and db/db mice. 1-methyl-nicotinamide was measured with metabolomics. NR treatment increased NAD^+^ level and 1-methyl-nicotinamide level in both groups. B) & C) Acetylated-lysine protein expression levels in total kidney lysates and mitochondrial fractions were increased in db/db mice and NR supplementation decreased its abundance. D) SIRT3 protein abundance in whole tissue lysate and E) SIRT3 enzyme activity in mitochondrial fraction were reduced significantly in db/db kidney and reduction was prevented upon NR treatment. In human diabetic kidneys F) Immunohistochemical (IHC) staining showing the SIRT3 protein abundance is significantly lower compared to non-diabetic human kidneys G) Histology quantification of nuclear Sirt1 expression levels were unchanged between non-diabetic and diabetic human kidneys. Western blot analysis of kidney mitochondrial lysates showing increased H) protein expression levels of total acetylated lysine and acetylated superoxide dismutase 2 (SOD2) and isocitrate dehydrogenase 2 (IDH2) in kidney of SIRT3 KO mice. In total kidney lysates, the abundance of I) acetylated SOD2 and J) acetylated IDH2 were increased in db/db mice and prevented in mice treated with NR. n=6 per group, values presented as mean ± SEM with variance is calculated using one-way ANOVA.

There were increases in acetylated lysine levels in whole kidney lysate **(Figure 6B)** and in isolated mitochondrial fractions **(Figure 6C)** in db/db mice, an indication of decreased deacetylase activity. Treatment with NR restored acetylated lysine proteins to levels seen in db/m mice, suggesting the improved deacetylase activity in the mitochondria. SIRT3 is the main mitochondrial deacetylase and is involved in regulating mitochondrial functions, including fatty acid oxidation (FAO). To explore whether SIRT3 was involved in the NR effect, we examined SIRT3 expression and activity. SIRT3 protein abundance (**Figure 6D**) and activity (**Figure 6E**) were reduced in the diabetic kidney. NR treatment increased SIRT3 protein abundance **(Figure 6D)** and SIRT3 activity **(Figure 6E)**. In human diabetic kidneys, SIRT3 expression (**Figure 6F, Supplementary Table S2**) was also decreased. Another NAD+-dependent deacetylase SIRT1 expression did not change in human diabetic kidneys when compared with the non-diabetic controls (**Figure 6G, Supplementary Table S2**).

To further assess a role for SIRT3 in regulating acetylated lysine protein in the kidney, we studied SIRT3 KO mice ^34, 35^. In SIRT3 KO kidneys, total protein acetylation level was increased, as well as the acetylation of SOD2 and IDH2 (**Figure 6H)**, two proteins previously identified as SIRT3 targets ^36, 37^. In diabetic kidneys, acetylated SOD2 and IDH2 protein levels are increased, and NR treatment decreased acetylated SOD2 and IDH2 protein levels (**Figure 6I and 6J**). Acetylation of K68 and K122 residues of SOD2 has been reported to regulate SOD2 activity ^37^.

### NR treatment enhanced mitochondrial biogenesis in diabetic kidneys

NR treatment increased the mitochondrial DNA/nuclear DNA ratio in db/db kidneys, indicative of increased mitochondrial biogenesis (**Figure 7A**). NR treatment also increased PGC1α mRNA and protein abundance (**Figure 7B**). PGC1α is a master mitochondrial biogenesis regulator ^38^. As expected, the direct targets of PGC1α, *Nrf1* and the mitochondrial transcription factor *Tfam1,* expression were increased in the diabetic kidney following the NR treatment (**Figure 7C**).

**Figure 7:**
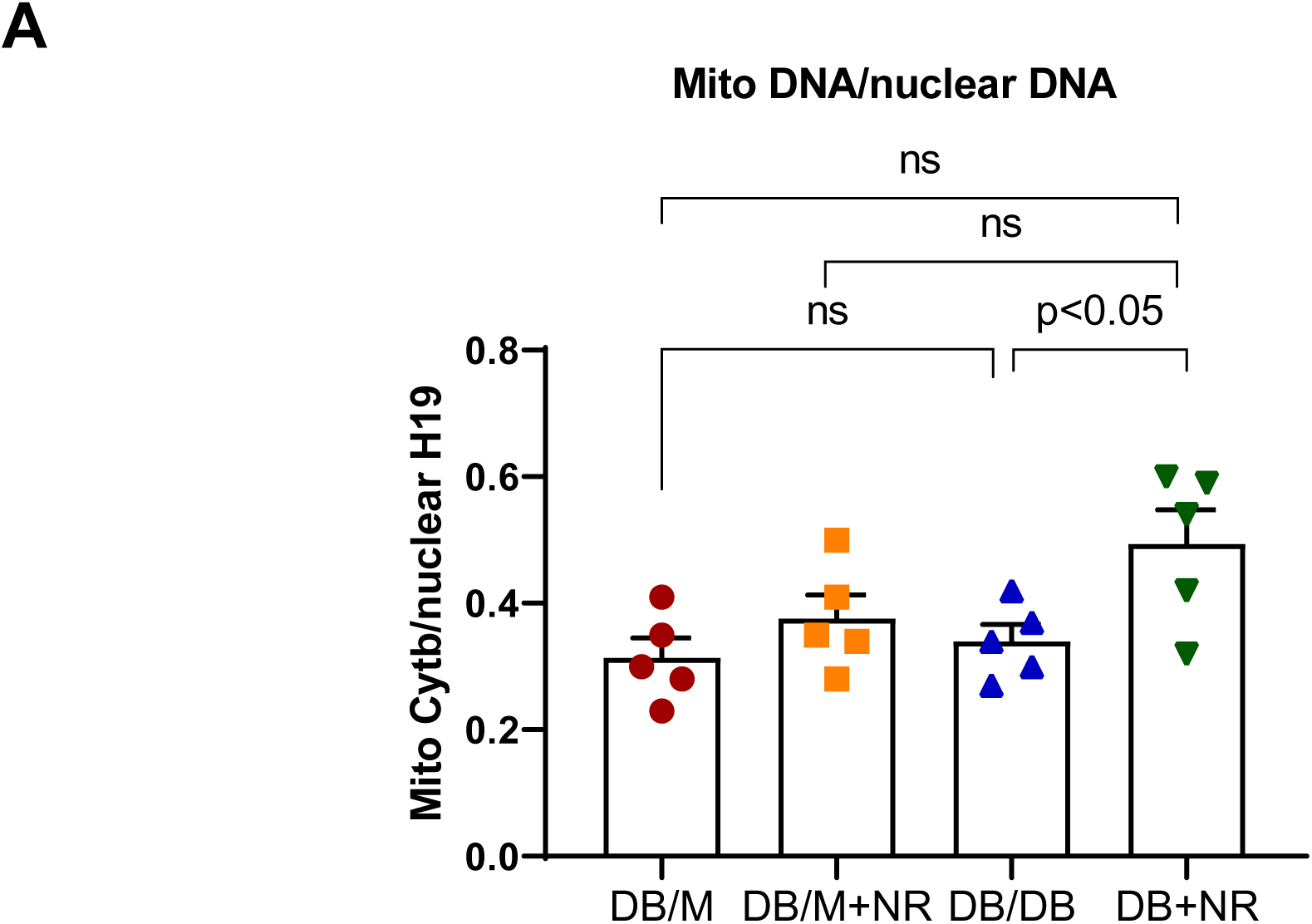
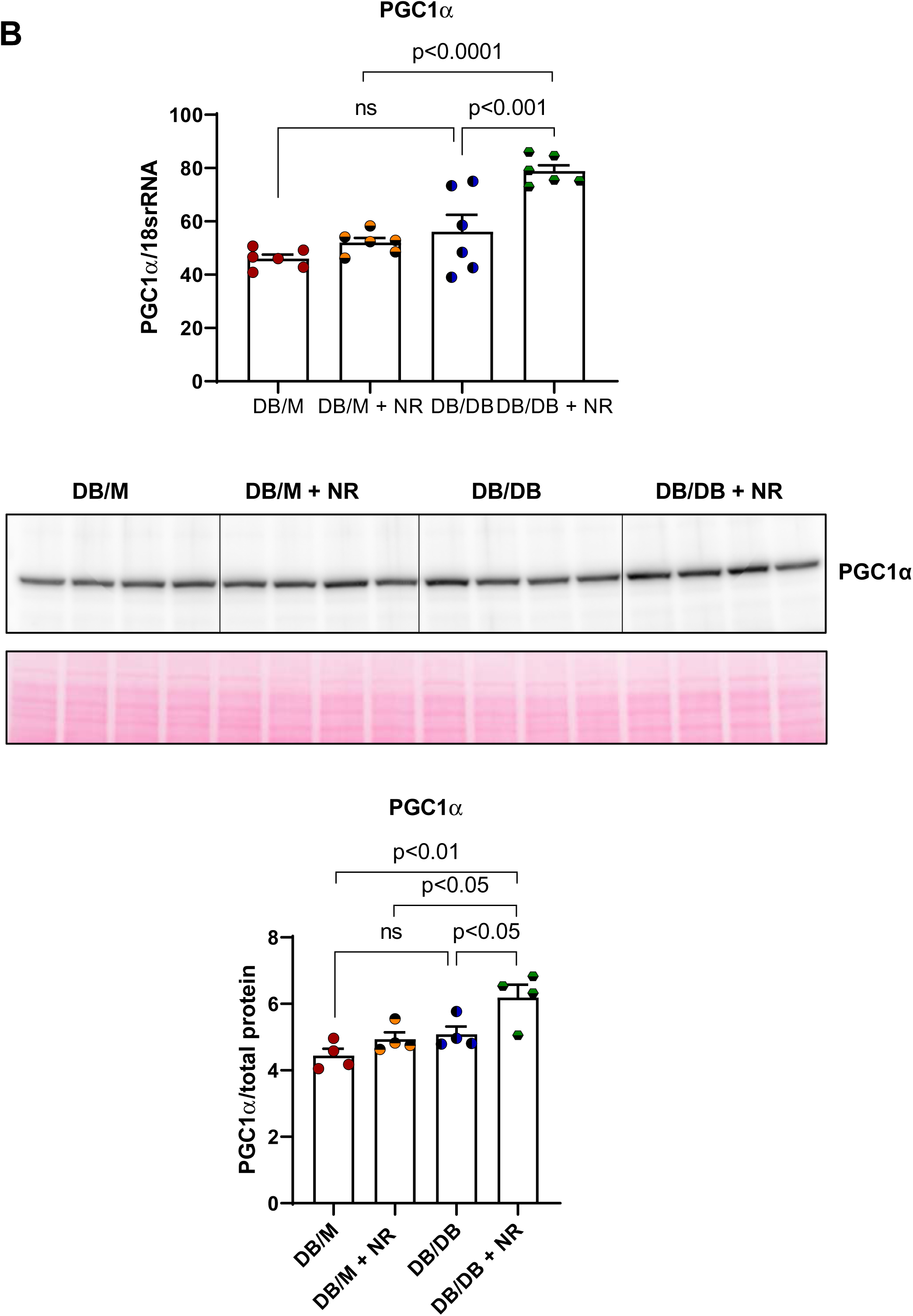
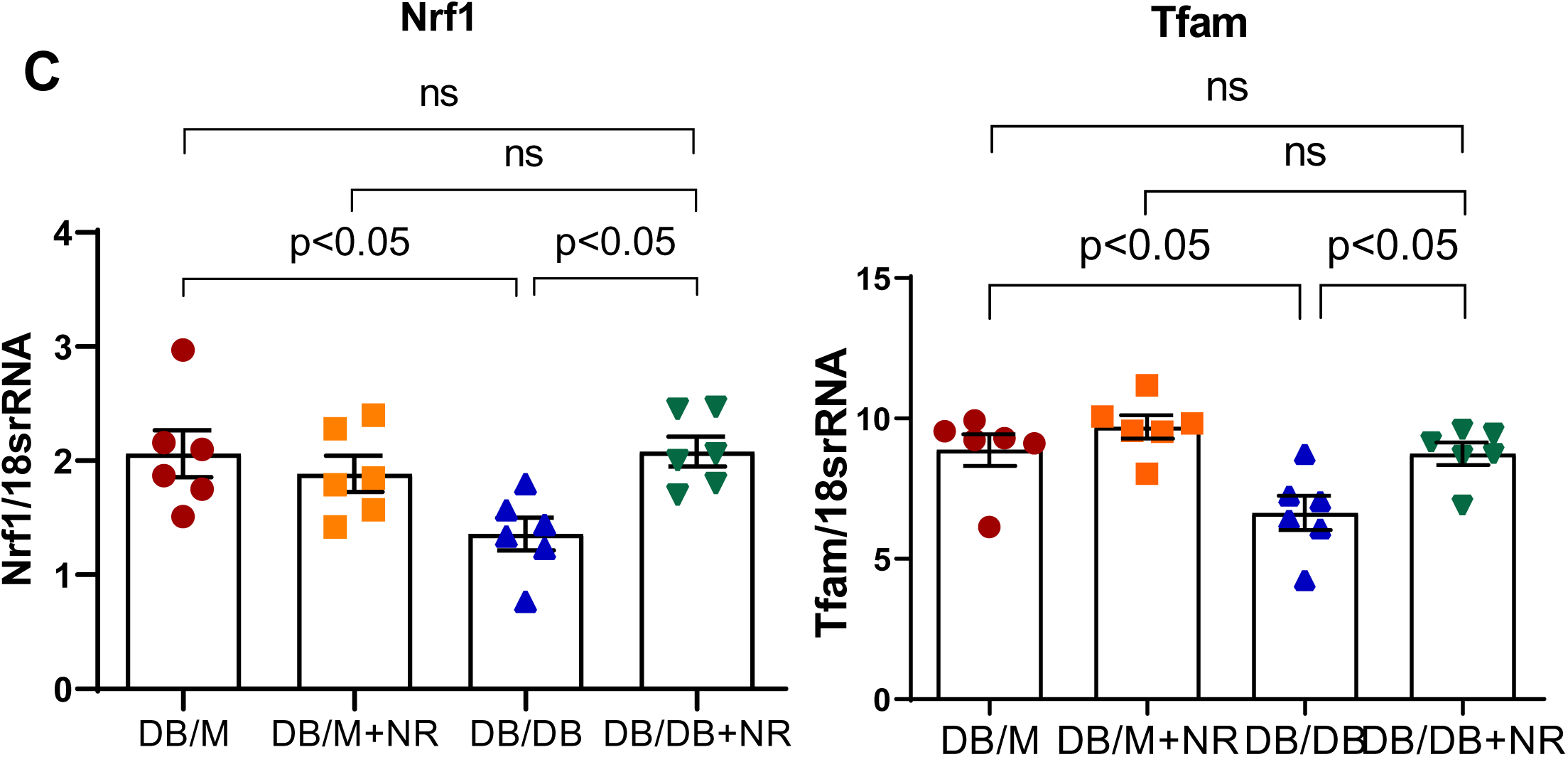
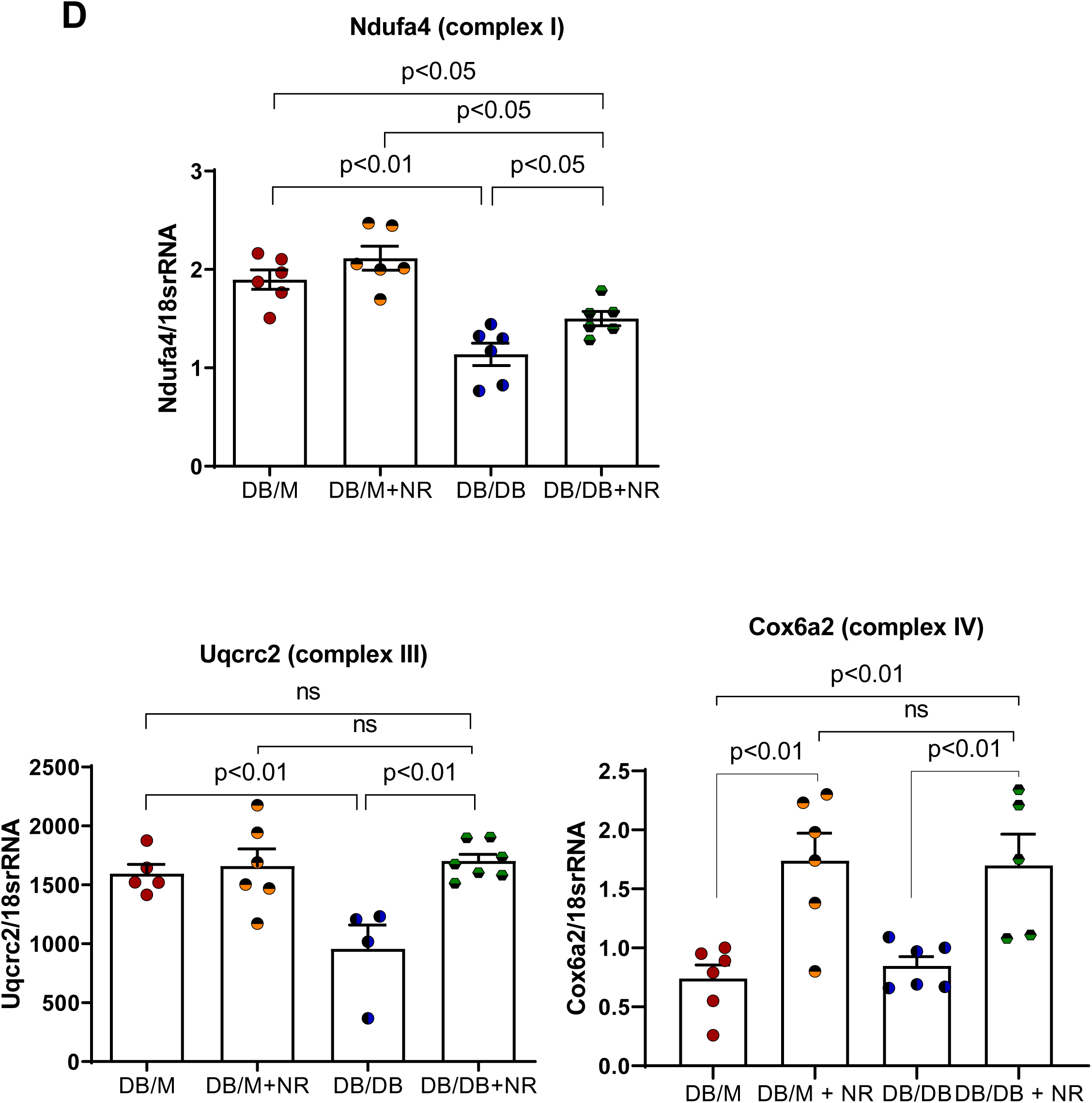
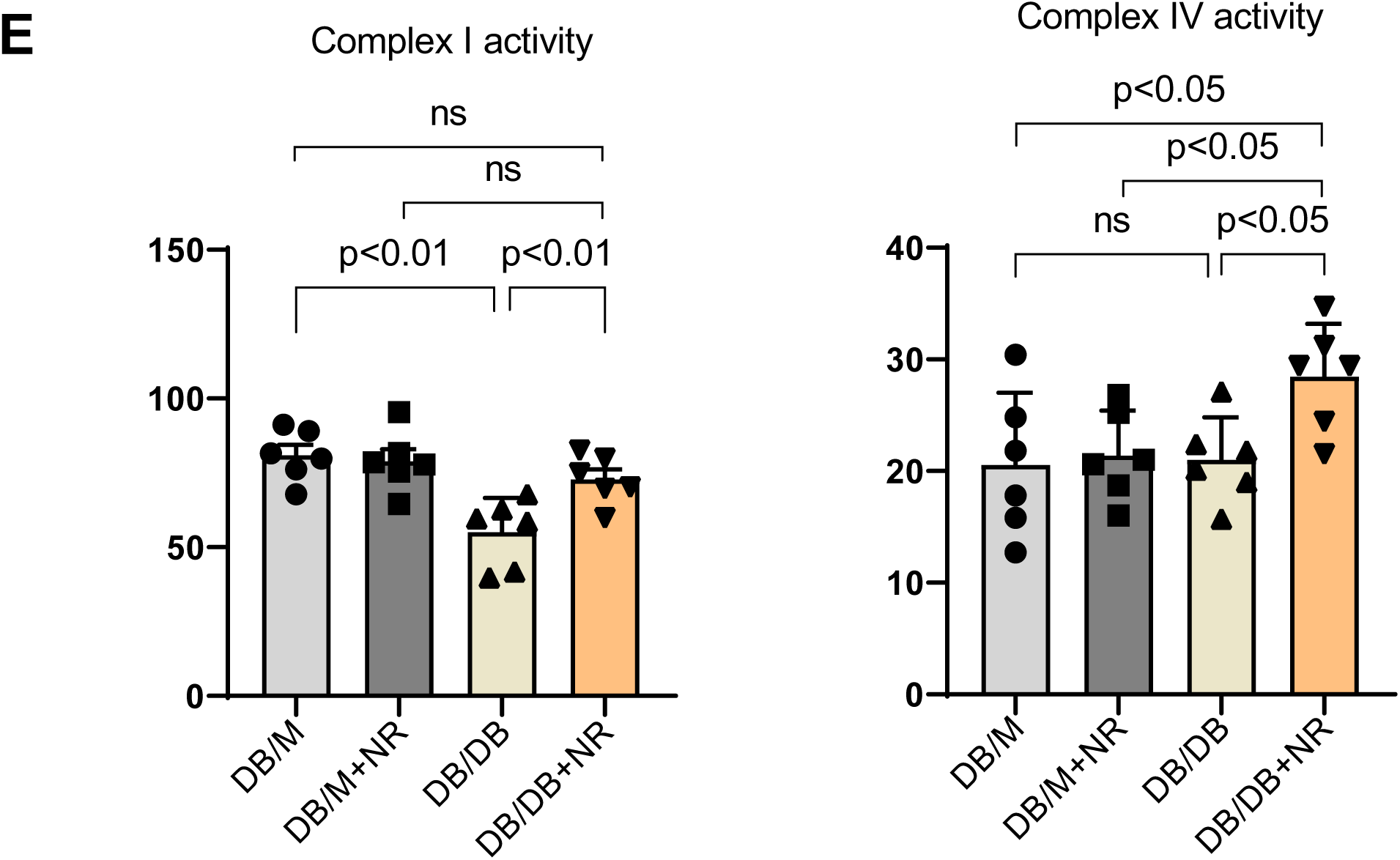
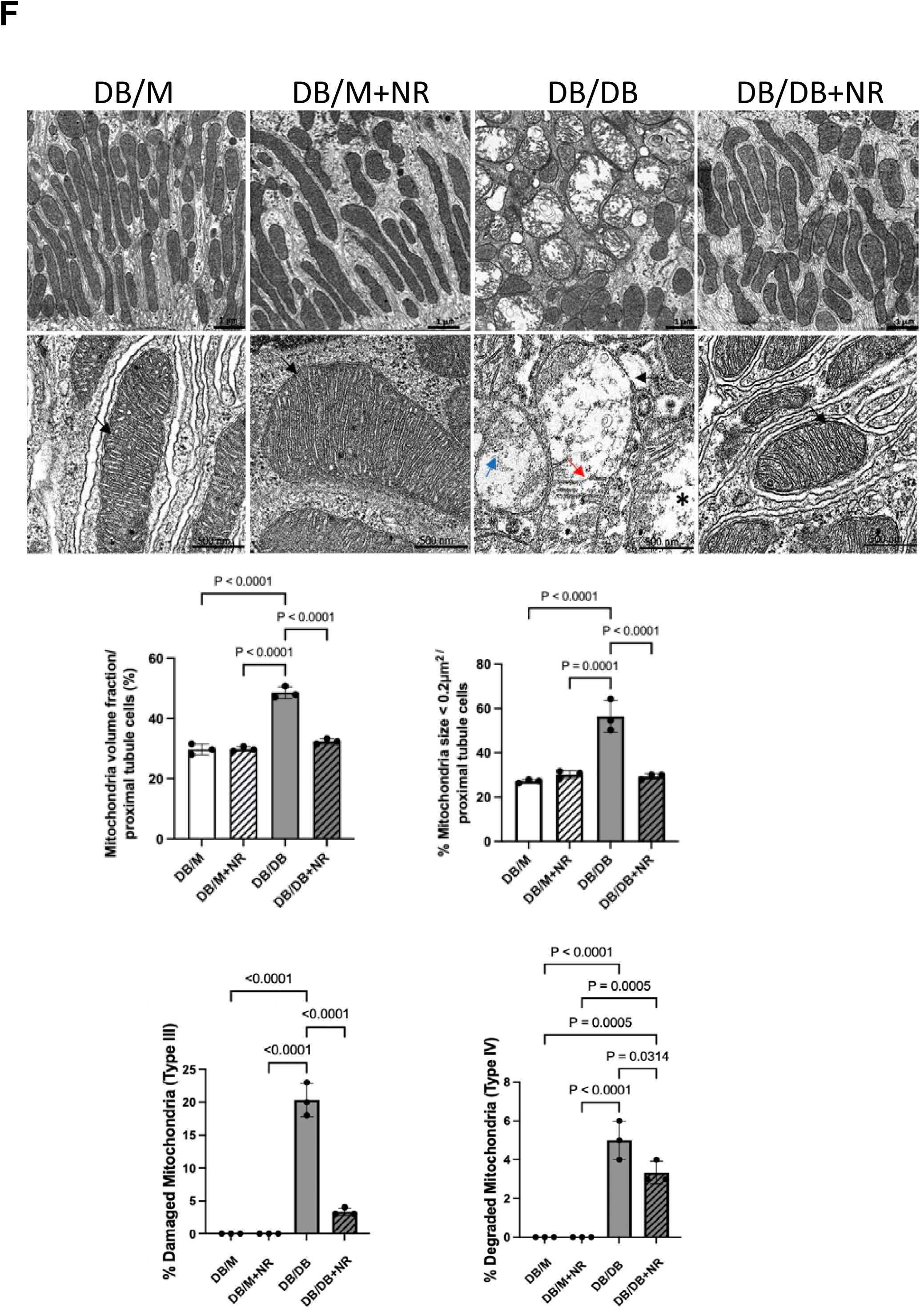
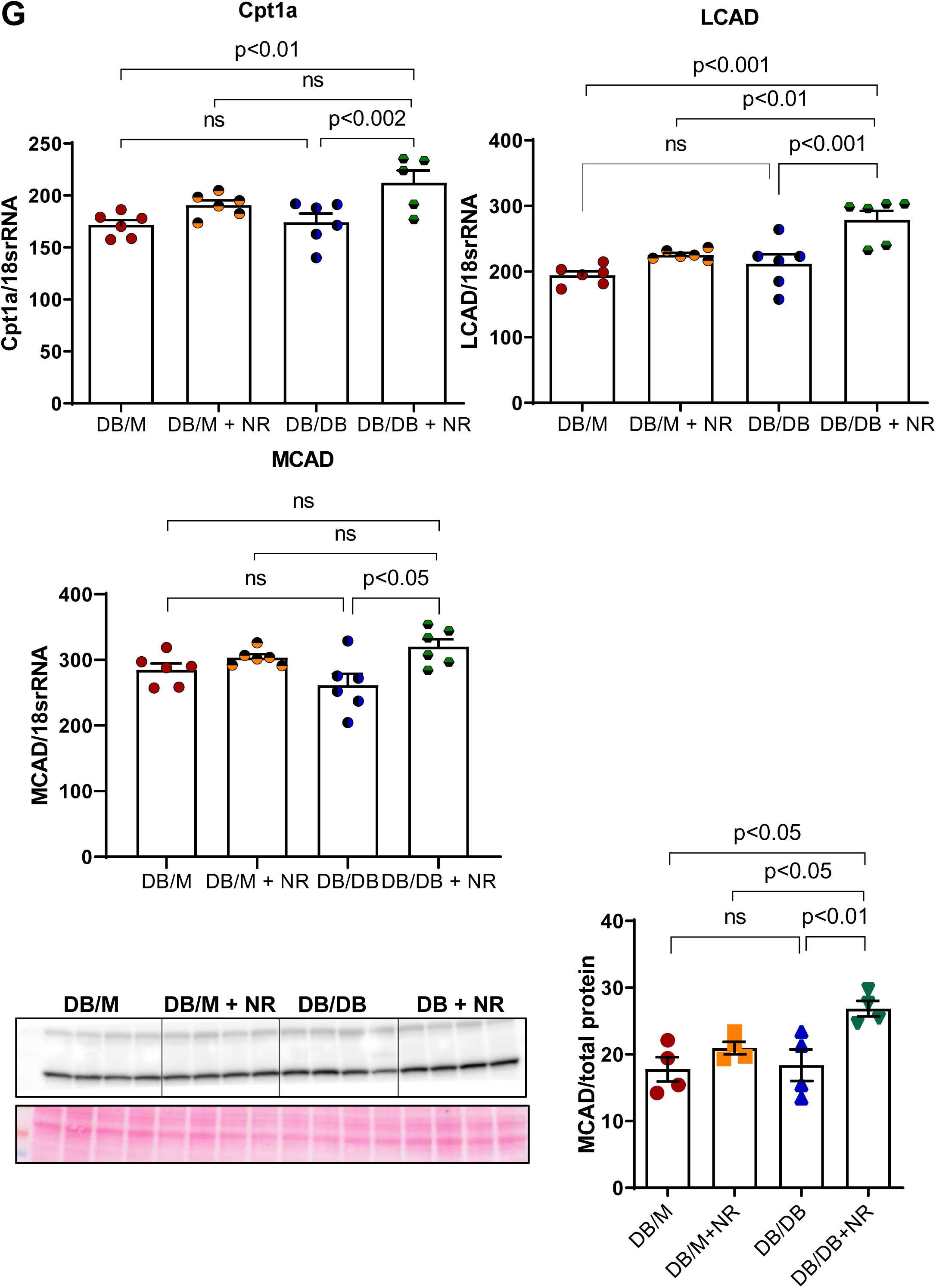
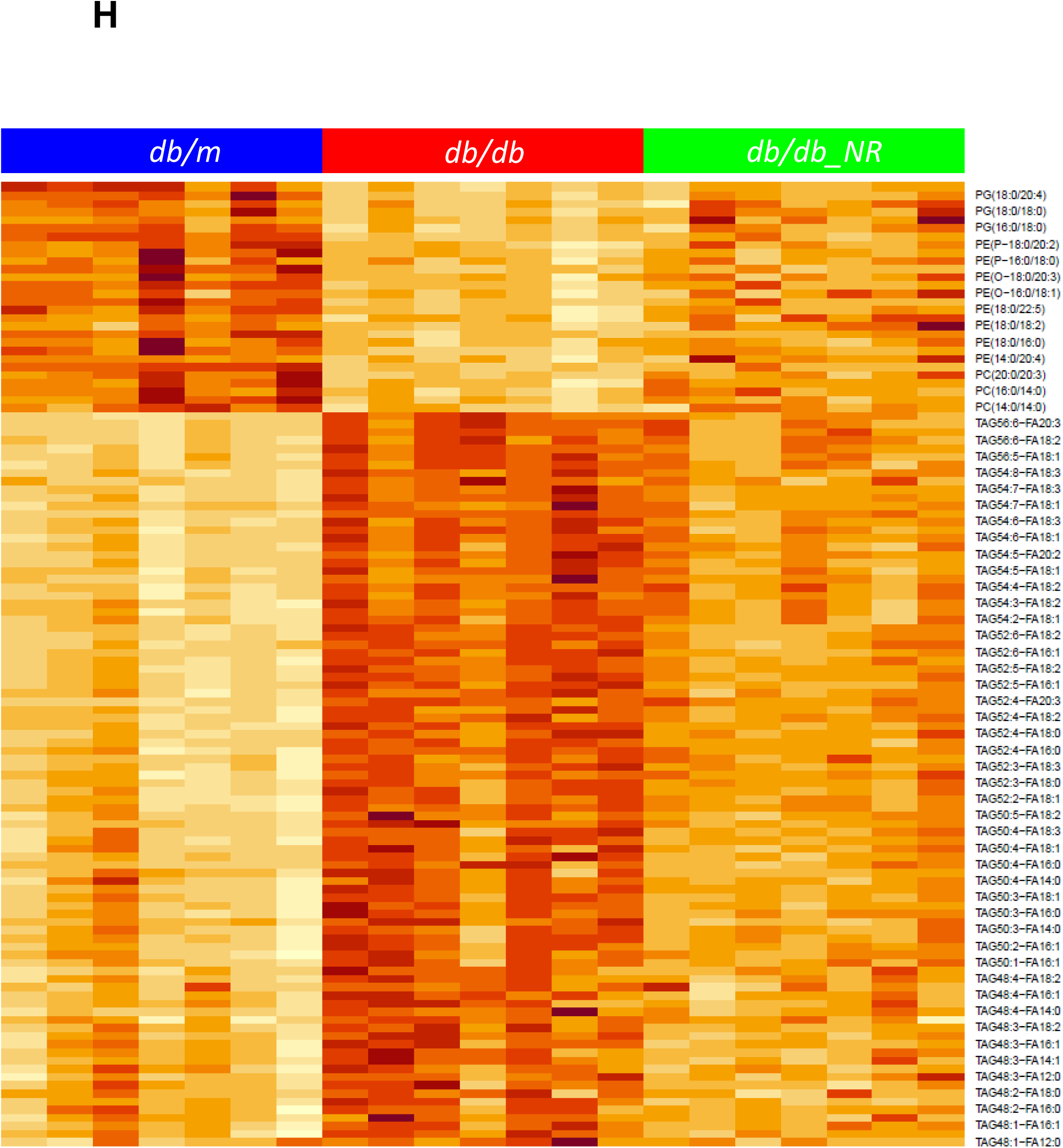
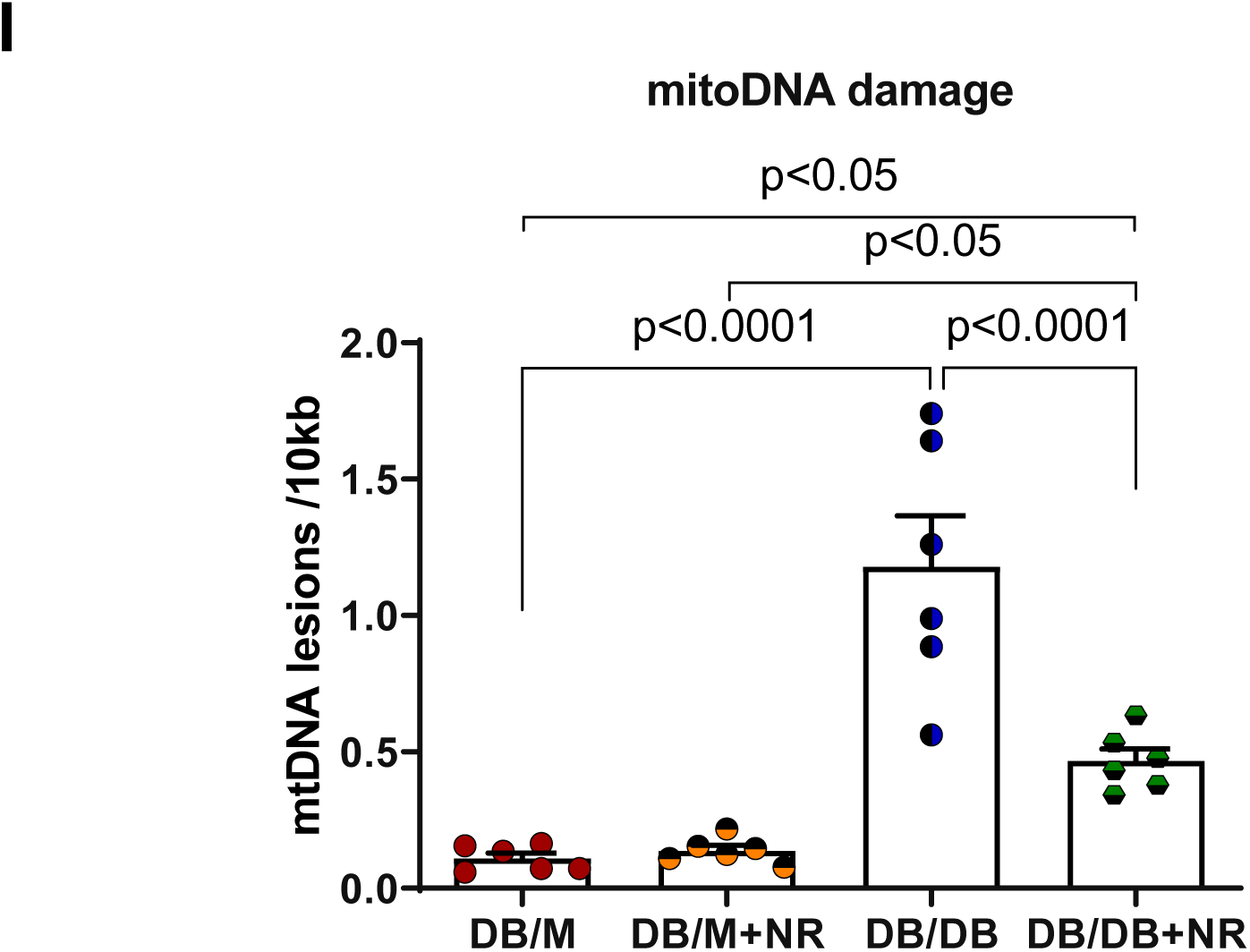
Effect of NR treatment on mitochondria in the kidney of diabetic db/db mice. A) mitochondrial DNA/nuclear DNA ratio was unchanged in db/db mice and NR supplementation increased the ratio of mito DNA. B) mRNA and protein expression levels of PGC1a a master regulator of mitochondrial biogenesis was increased with NR treatment in db/db mice though there is comparable differences between db/m and db/db mice. C) mRNA levels of *Nrf2* and *Tfam* were decreased in db/db mice and prevented the decreasing upon NR treatment. D) Complex 1 and III subunits *Ndufa4* and *Uqcrc2* mRNA levels were reduced significantly in db/db kidney and reduction was inhibited with NR treatment. Complex IV subunit *Cox6a2* mRNA levels were increased in both db/m and db/db kidney with NR supplementation. E) Complex 1 enzyme activity was markedly decreased in db/db mice and reversed in response to NR treatment. In addition, complex IV activity also increased by NR treatment in db/db mice. F) Representative electron microscopy images showing mitochondria morphology in proximal tubule cells. SEM images in db/db mice show a chaotic mitochondrial distribution and damaged mitochondria in db/db mice and an improvement with NR treatment. Scale bars 1 um. Magnifications 35,000. Higher magnification of TEM images show normal cristae organization (arrow) in db/m mice and damage to the mitochondria in db/db mice such as cristae fragmentation (red arrow), cristae homogenization (blue arrow) in swollen electron-lucent matrix (asterisk) and ruptured outer membrane (black arrow). NR treatment db/db mice resulted in normalization of mitochondrial stricture similar to the control (arrows). Scale bars 500 nm. Magnificationsx 50,000. The graphs show the percentage of mitochondria volume, mitochondria size < 0.2 um and mitochondrial types in db/m and db/db and NR treated mice. n=3 per group. G) The fatty acid β-oxidation enzymes *Cpt1a*, *Lcad* mRNA and *Mcad* mRNA and protein levels were increased with NR administration in db/db mice. H) Lipidomics data showed increased triglycerides species in db/db kidneys and NR reduced the increase. I) Mitochondrial DNA (mtDNA) damage markedly increased in db/db mice and damage was prevented by NR. n=5-6 per group, values presented as mean ± SEM with variance is calculated using one-way ANOVA (except in F where two-way ANOVA was used).

Increased mitochondrial biogenesis was accompanied by increased expression of genes related to the mitochondrial ETC complexes, including complex I subunit *Ndufa4*, complex III subunit *Uqcrc2* and complex IV subunit *Cox6a2* (**Figure 7D**). Complex I activity was decreased in the diabetic kidneys and treatment with NR restored complex I activity. NR treatment also restored complex IV activity (**Figure 7E**).

In db/db mice, mitochondria within proximal tubule epithelial cells exhibited severe polymorphic structural changes. Based on the type of mitochondrial restructuring, 4 categories were assigned (I-IV), as previously described ^39^. Type I mitochondria are oval-shaped, with longitudinally oriented and tightly packed cristae. Type II mitochondria are structurally abnormal, with indistinct shape and/or non-uniform size together with hypoplasia. The cristae are swollen and/or have signs of homogenization, irregular or whirling, which usually have lost the longitudinal orientation, tightness, regular spacing, and electron-lucent matrix. Type III mitochondria manifest hypoplasia and degenerative changes. The shape and size of these mitochondria vary, often with a discontinuous outer membrane. The focal disruption of the inner membrane leads to an uneven increase in the crista thickness, homogenization and fragmentation, with a swollen electron-lucent matrix. Type IV mitochondria exhibit disrupted and discontinuous outer membranes, deficiency in cristae and “myelin-like” cristae transformations (**Figure 7F**). In db/db mice, up to 20% of the total mitochondria corresponded to type III and up to 5% were found to have type IV structural damage. These manifestations were absent in all db/m mice. NR treated db/db mice exhibited a significant decrease in severely damaged mitochondria compared to untreated db/db mice (**Figure 7F**).

Morphometric analysis revealed that in db/m control mice the mitochondrial area ranged from 0.2 - 0.9 μm2 and mitochondria < 0.2 μm2 constituted up to 24% of all mitochondria. In diabetic db/db mice, mitochondria < 0.2μm2 were found up to 58% which was significantly higher than in control mice (**Figure 7F**). The increase in mitochondria < 0.2μm2 size indicates the enhancement of mitochondria fission. NR treated diabetic mice showed a significant reduction in number of mitochondria < μm2 size, and this may suggest the improvement of mitochondrial functional. In addition, renal expression of mitochondrial enzymes that mediate mitochondrial fatty acid β-oxidation, including *Cpt1a* mRNA, *Lcad* mRNA, and *Mcad* mRNA and MCAD protein, were all upregulated by NR treatment of db/db mice (**Figure 7G**), suggesting that NR treatment promotes mitochondrial fatty acid β-oxidation. Consistent with these effects, NR treatment prevented triglyceride accumulation in the kidney (**Figure 7H**).

Finally, we found increased mtDNA damage in the db/db kidneys and NR treatment protected the kidney from mtDNA damage (**Figure 7I**). mtDNA damage triggers the leakage of mtDNA into cytosol, activating the nucleic acid sensors such as cGAS and STING.

## DISCUSSION

Inflammation and mitochondrial dysfunction have been proposed to play an important role in the progression of diabetic kidney disease ^40–43^. It is not clear whether and how these two processes are linked with each other. We have found that treating diabetic mice with nicotinamide riboside (NR), an NAD+ precursor, improved inflammation as well as mitochondrial function.

The increases in the interferon-induced transmembrane proteins 1,2, and 3 (IFITMs) from RNAseq prompted us to find the regulation of cGAS/STING signaling in diabetic kidneys. A role for increased STING expression and activity per se in mediating inflammation is demonstrated in studies where we inhibited STING with a well-established inhibitor C-176 and also in STING knockout mice made diabetic with streptozotocin. In both studies we found that STING inhibition prevents inflammation. In addition, and importantly we determined that STING inhibition prevents diabetic kidney injury as demonstrated by preventing increases in urine albumin and urine KIM1 excretion.

While we failed to document a working phospho-STING antibody for the mouse tissue with STING knockout mice, total STING antibody has been found working in the staining of kidney tissue. We found that STING expression is also increased in the kidneys of human subjects with diabetes. STING staining of control kidney biopsies show endothelial staining in glomeruli, peritubular capillaries and larger vessels with sparse staining of interstitial inflammatory cells. In diabetic renal parenchyma, an expanded pattern of staining is observed. There is increased endovascular staining in glomeruli correlating with segments with prominent endothelium and endocapillary inflammatory cells including lymphocytes and monocyte/macrophages. Prominent parietal epithelial cells show increased expression. By far, the most prominent compartment with STING expression are interstitial inflammatory cells. Also noted is enhanced staining in two tubular elements: atrophic tubule and distal nephron. Although STING has been previously reported in other kidney injury models ^31, 33, 51^, this is the first time that the localization of STING activation in the kidney has been examined. In contrast to previous reports, the proximal tubules did not seem to be a major site for STING activation in diabetic kidneys. The infiltrated immune cells express most of the STING found in the kidney. The expression of STING in other cell types such as podocytes, in diabetic kidneys, may well be a result of co-purifying immune cells. This raises the question of how the mitochondrial DNA damage in the tubules or podocytes relates to STING activation in the immune cells, which warrants further investigation.

From STING inhibition studies we found that this inhibition did not achieve what NR treatment showed in db/db kidneys. This could be due to other players of nucleic acid sensors that we also found to be activated in diabetic kidneys, and blocked by NR, such as AIM2 and NLRP3. This indicates that NR can target to a mechanism upstream of other nucleic acid sensors besides cGAS/STING, making itself a better treatment than STING inhibition alone.

NR treatment improved several parameters of mitochondrial dysfunction including restoration of mitochondrial fatty acid β-oxidation. This may be mediated by increasing the mitochondrial sirtuin 3 activity. The increase in sirtuin 3 activity was associated with decreasing the acetylation of proteins important for mitochondrial function including SOD2 and IDH2. The increase in the mitochondrial antioxidant SOD2 is reflected by the ability of NR to decrease renal oxidative stress. NR treatment also induced an increase in mitochondrial DNA/nuclear DNA ratio, indicative of an increase in mitochondrial biogenesis, as well as increases in Tfam, Nrf1, and PGC-1α, complex I and complex IV activities, and enzymes that mediate mitochondrial fatty acid β-oxidation (FAO). Other NAD+ precursors have also been reported beneficial renal effects ^44, 45^. However, in this report, we further connected the NAD+ effects in diabetic kidney disease model to its direct target SIRT3 by showing the increase in SIRT3 deacetylase activity and the decrease of acetylation level in the mitochondrial targets of SIRT3. SIRT3 relies on the availability of NAD+ level to improve the mitochondrial functions including FAO ^46–48^. Restoring FAO is a critical step to reverse the kidney injury. Recently, increasing the FAO enzyme Cpt1a in renal tubules was shown to protect against kidney fibrosis ^49^.

How these mitochondrial functional changes relate to regulation of inflammation is not fully known, however in diabetes there is evidence for increased mitochondrial DNA damage which is prevented by NR treatment. The increase in mitochondrial DNA damage may be mediated by increased oxidative stress and also by decreases in Tfam and Nrf1 ^50^. The increase in mitochondrial DNA damage on the other hand can result in activation of nucleic acid sensor signaling, which are major mediators of inflammation via inducing the NFκB and STAT3 regulated inflammatory pathways. Treatment with NR prevents the decreases in Tfam and Nrf1, the increase in mitochondrial damage, and the increases in the interferon-induced transmembrane proteins 1,2, and 3, NFκB and STAT3. Overall, our data showed that NR supplementation boosted the NAD metabolism to modulate inflammation and mitochondrial function and prevent progression of diabetic kidney disease.

## METHODS

### Animal studies

All mouse experiments were conducted according with the Guide for Care and Use of Laboratory Animals, National Institute of Health, Bethesda, MD and were approved by Institutional Animal Care and Use Committee of Georgetown University, Washington, D.C.

10-week-old male db/m (non-diabetic controls) (catalog # 00662) and db/db (diabetic) (catalog # 00642) mice were obtained from Jackson Laboratories (Bar Harbor, ME). The mice were housed in animal care facility with 12/12-hour light-dark cycle and fed for 20 weeks on regular chow diet (TD 190694, Envigo, Madison, WI) or chow diet supplemented with 500 mg/kg body weight nicotinamide riboside (NR) (ChromaDex, Irvine, CA) (TD 190695, Envigo, Madison, WI). One week prior to the end of the study, the mice were placed in metabolic cages for a 24-hour urine collection. At the end of the study, heparinized plasma and kidneys were harvested for further processing.

To determine the role of SIRT3, whole body SIRT3 knockout mice on C57BL/6 background were obtained from Matthew Hirschey (Duke University) ^35^ to compare their kidneys with the age, gender matched wild-type control kidneys.

To determine the role of STING inhibition in the diabetic kidneys, 20-week-old male db/m and db/db mice were treated with C-176 ^32^ (Focus Biomolecules, Plymouth Meeting, PA) at 1.2mg/kg body weight dose with daily i.p. injection for 4 weeks.

To further determine the role of STING in the diabetic kidneys, 12-week-old male wild type and whole-body STING knockout mice on C57BL/6J background (JAX catalog #025805) were treated with streptozotocin at 50mg/kg body weight as 5-day daily consecutive i.p. injection to induce diabetes. These mice were sacrificed after 12 weeks on diabetes.

### Blood and urine biochemical analysis

Blood glucose levels were assessed with a glucometer (Elite XL, Bayer, Tarrytown, NY). Blood urea nitrogen (BUN) was measured with colorimetric QuantiChrom assay kit (Bioassay systems, Hayward, CA). Total cholesterol and triglycerides levels in plasma were determined with calorimetric assay kit provided by Pointe Scientific (Canton, MI, USA). Urine assays followed the instructions from the kits listed in **Supplementary Table S1.**

### NAD^+^ measurement

20 mg of kidney tissue was homogenized in extraction buffer provided by the kit (E2ND-100, Bioassay systems) and NAD^+^ was determined immediately according to manufacturer’s protocol.

### Immunoblotting

Total protein was quantified using the BCA protein assay kit (Thermo Scientific, Rockford, IL). Western blotting was done as previously described ^16^. Antibody information can be found in **Supplementary Table S1.**

### Quantitative Real-Time PCR

Total RNA from kidneys were isolated according to manufacturer’s protocol with Qiagen RNeasy mini kit (Qiagen, Gaithersburg, MD) and cDNA was made using reverse transcript reagents from Thermo Scientific (Catalog # 4374967). qRT–PCR was performed using Quant Studio Real-Time PCR machine (Thermo Fisher Scientific). Expression levels of target genes were normalized to 18S level. Primer sequences are listed in **Supplementary Table S1.**

### Mitochondrial enzymatic complex activity assay and sirtuin 3 activity assay

Mitochondrial fraction was isolated according to previously described protocol ^16^. Isolated mitochondria were assayed for the complex I (Catalog # ab109729), complex IV (Catalog # ab109911) and sirtuin 3 activity (Catalog # ab156067) with the kits purchased from Abcam, Boston, MA.

### Analysis of mitochondrial DNA damage

Total DNA from kidney tissue was isolated using Genomic Tip (Qiagen, Valencia, CA), and its concentration was measured by PicoGreen dye (Invitrogen, Carlsbad, CA). 15 ng DNA was used to amplify a long 10kb mitochondria DNA target followed by real-time PCR based quantification to determine mitochondria DNA lesions, as previously described ^52^.

### Immunohistochemical (IHC) and PAS staining

Immunohistochemical staining was performed on formalin fixed and paraffin embedded 5 µm kidney sections. Following deparaffinization and rehydration, the slides were subjected to heat mediated antigen retrieval in citrate buffer pH 6 and blocked with 3% BSA. The sections were probed with Sirt1 (Abcam catalogue # ab110304), Sirt3 (Sigma catalogue # S4072) or STING (Cell Signaling, catalogue # 13647S) antibody and incubated in room temperature for 1.5 hours. Mouse/Rabbit PolyDetector reagent (Bio SB, Catalog No. BSB 0269) or UnoVue HRP secondary antibody detection reagent (Diagnostic BioSystems, Pleasanton, CA) was applied followed by DAB chromogen. The Periodic Acid-Schiff (PAS) staining was performed with a PAS stain kit (Thermo Scientific, Catalog No. 87007). Imaging was done with Nanozoomer (Hamamatsu Photonics, Japan) and Motic Digital Slide Scanner (Richmond, BC, Canada).

### Quantification of Morphology

Glomeruli were extracted from images of PAS staining and the PAS components of each glomerulus were segmented as described before ^53^. To quantify mesangial expansion, the ratio of PAS positive pixels to detected glomerular pixels was used.

### Immunofluorescence microscopy

The kidney tissue was snap frozen by embedding in to optimum cutting temperature (OCT) medium (Thermo Scientific, CA). The tissues were sectioned at 5-µm in thickness and transferred over the superfrost slides. Immunofluorescence staining performed as descried previously ^16, 54^. Antibody information can be found in **Supplementary Table S1.**

### Electron Microscopy

Renal cortex tissues were fixed in the 2.5% glutaraldehyde/2% paraformaldehyde/ 0.05M cacodylate solution, post-fixed with 1% osmium tetroxide, and embedded in EmBed812. For imaging acquisition, ultrathin sections (70 nm) were post-stained with uranyl acetate and lead citrate and examined in the Talos F200X FEG *transmission electron microscope* (FEI, Hillsboro, OR) at 80 kV located at the George Washington University Nanofabrication and Imaging Center. Digital electron micrographs were recorded with the TIA software (FEI). Ultrathin sections (120 nm) were mounted in silicon wafers and observed with a Teneo LV FEG *scanning electron microscope* (FEI, ThermoFischer Scientific). For optimal results, we used the optiplan mode (high-resolution) equipped with an in-lens T1 detector (Segmented A+B, working distance of 8 mm). Low-magnification images (600×) were first taken for the observation then we performed high magnification tile images of our regions of interest (35,000) using 2LkV and 0.4 current landing voltage.

Morphometric analysis was performed under blinded conditions by systematic uniform random sampling with the Fiji Software using 20 randomly selected images. In EM images, volume fraction of mitochondria was determined using the morphometric technique with a dot grid. The size of each individual mitochondria was calculated by using the FIJI ImageJ software. Mitochondrial types were determined using the point counting method ^55^.

### Bulk RNA-seq

One microgram of total RNA samples was sent to Novogene (Sacramento, CA) for mRNA sequencing. RNA-seq fastQ files were filtered and trimmed from adaptors using Trimmomatic algorithm ^56^. The reads were aligned to *Mus musculus* genome assembly and annotation file GRCM38:mm10 using STAR algorithm ^57^. Gene expression was estimated in FPKM counts using RSEM algorithm ^58^. Differential expression was quantified with DeSeq2 algorithm ^59^. Absolute fold change of 1.5 and Bonferroni adjusted p-value of less than 0.05 was considered as significant change. All bioinformatics analysis was performed on T-BioInfo Platform (http://tauber-data2.haifa.ac.il:3000/). DAVID Bioinformatics ^60, 61^ and PANTHER Classification System (http://PANTHERdb.org/) were used to classify the differentially expressed genes into functional groups. To identify proteins that are localized to mitochondria, we used a curated database of mitochondrial localized proteins – MitoCarta3 database ^62^.

### Proteomic analysis

Frozen mouse kidney samples were lysed in 8 M urea and 50 mM triethylammonium bicarbonate (TEAB). Proteins were then reduced and alkylated followed by digestion with LysC (Fujifilm Wako Pure Chemical, Osaka, Japan) in the ratio of 1:100 (enzyme-to-protein, w/w) at 37°C for 3 hours. Subsequently, the proteins were further digested with trypsin (Promega, Fitchburg, WI) in the ratio of 1:50 (enzyme-to-protein, w/w) at 37°C overnight after diluting the urea concentration from 8 M to 2 M. Proteins were then acidified, desalted, and lyophilized sequentially. The dried peptides were labeled with 16-plex TMT reagents (Thermo Scientific). The labeled peptides was fractionated using basic pH RPLC for total proteome analysis as described previously ^63^. The LC-MS/MS was analyzed on an Orbitrap Fusion Lumos Tribrid Mass Spectrometer coupled with an Ultimate3000 RSLCnano nano-flow liquid chromatography system (Thermo Scientific). The resulting spectra were analyzed by Proteome Discoverer (version 2.4.1.15 software package, Thermo Scientific) following standard procedures. Downstream analysis was performed with Perseus ^64^ using log2 normalized intensities of protein abundance. Absolute fold change of 1.5 and Bonferroni adjusted p-value of less than 0.05 was considered as significant change.

### Lipidomics

Kidney samples were pulverized in liquid nitrogen and dissolved in 300 μL of isopropanol extraction buffer containing internal standard for lipid classes. The samples were vortexed for 30 seconds and homogenized for 1-2 min on ice and incubated on ice for 20 min followed by incubation at -20 ℃ for 20 min. Samples were centrifuged at 13,000 rpm for 20 min at 4 ℃. The supernatant was transferred to MS vial for LC-MS analysis using QTRAP® 5500 LC-MS/MS System (Sciex, Framingham, MA).

### Statistical Analysis

All the resulted data sets were calculated and presented as meanL±LSEM. One-way ANOVA fallowed by Student-Newman-Keuls post hoc analysis were used to analyze the variance among multiple groups and between two groups. The statistically significant differences were designated as a *P* values of <0.05. GraphPad prism 8.1.2 software package was used for statistical analysis (www.graphpad.com).

### Data and Resource Availability

All reagents and data from this article are available from the corresponding author upon request.

## Acknowledgements

**Funding.** This study was funded by NIH R01 Grants DK116567 (ML), DK127830 (ML), F30 Fellowship DK129003 (BAJ), AHA Postdoctoral Fellowship 19POST34381041 (KM), and National Center for Advancing Translational Sciences of NIH under Award Number TL1TR001431 (BAJ). We also thank Chromadex (Los Angeles, CA) for supplying us NR.

**Duality of Interest.** No potential conflicts of interest relevant to this article were reported.

**Author Contributions.** ML and XXW conceived and designed research; KM, XXW performed most experiments; BAJ, MDH, XY, AZR, BG and PS performed the histology work; LB, YJ, CHN, YQ, XZ, UG, PL, CW, JM, AC, and JP conducted the –omics work and analysis. KM and XXW analyzed data and interpreted results of experiments; KM, XXW and JP prepared figures; XXW, KM, JP and ML wrote the manuscript with input from all the co-authors; ML is the guarantor of this work and, as such, had full access to all the data in the study and takes responsibility for the integrity of the data and the accuracy of the data analysis.

---

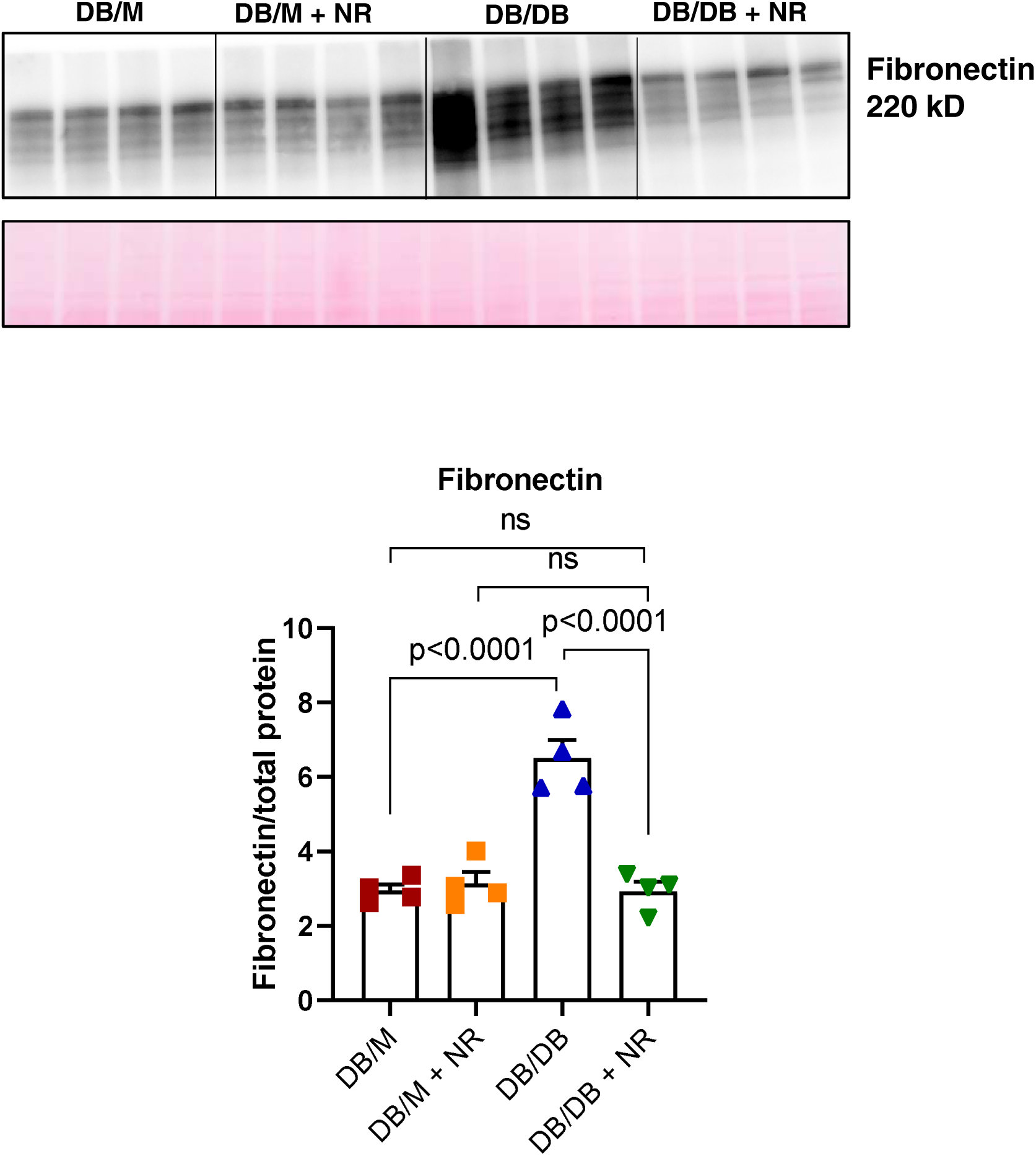
Figure S1

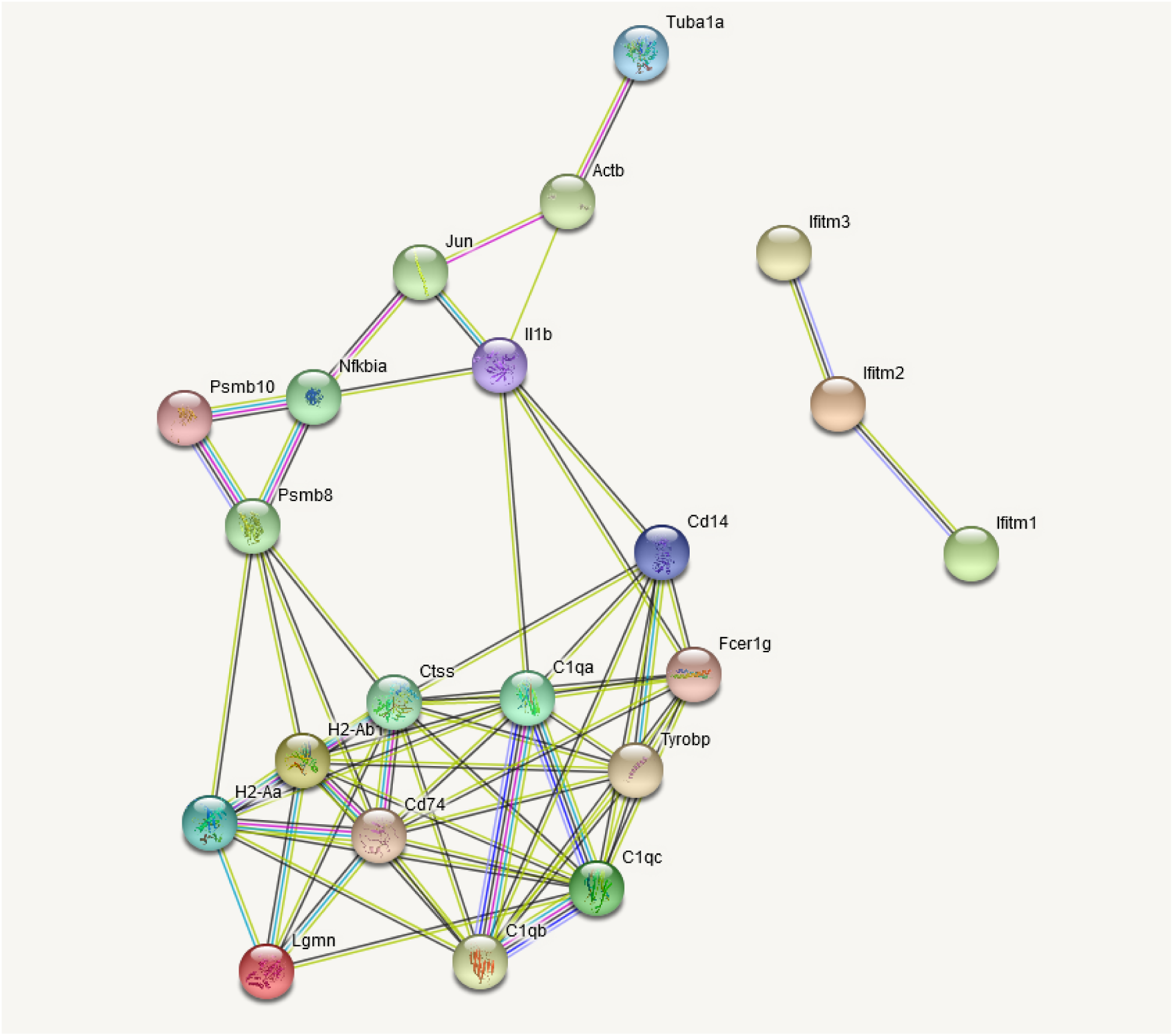
Figure S2

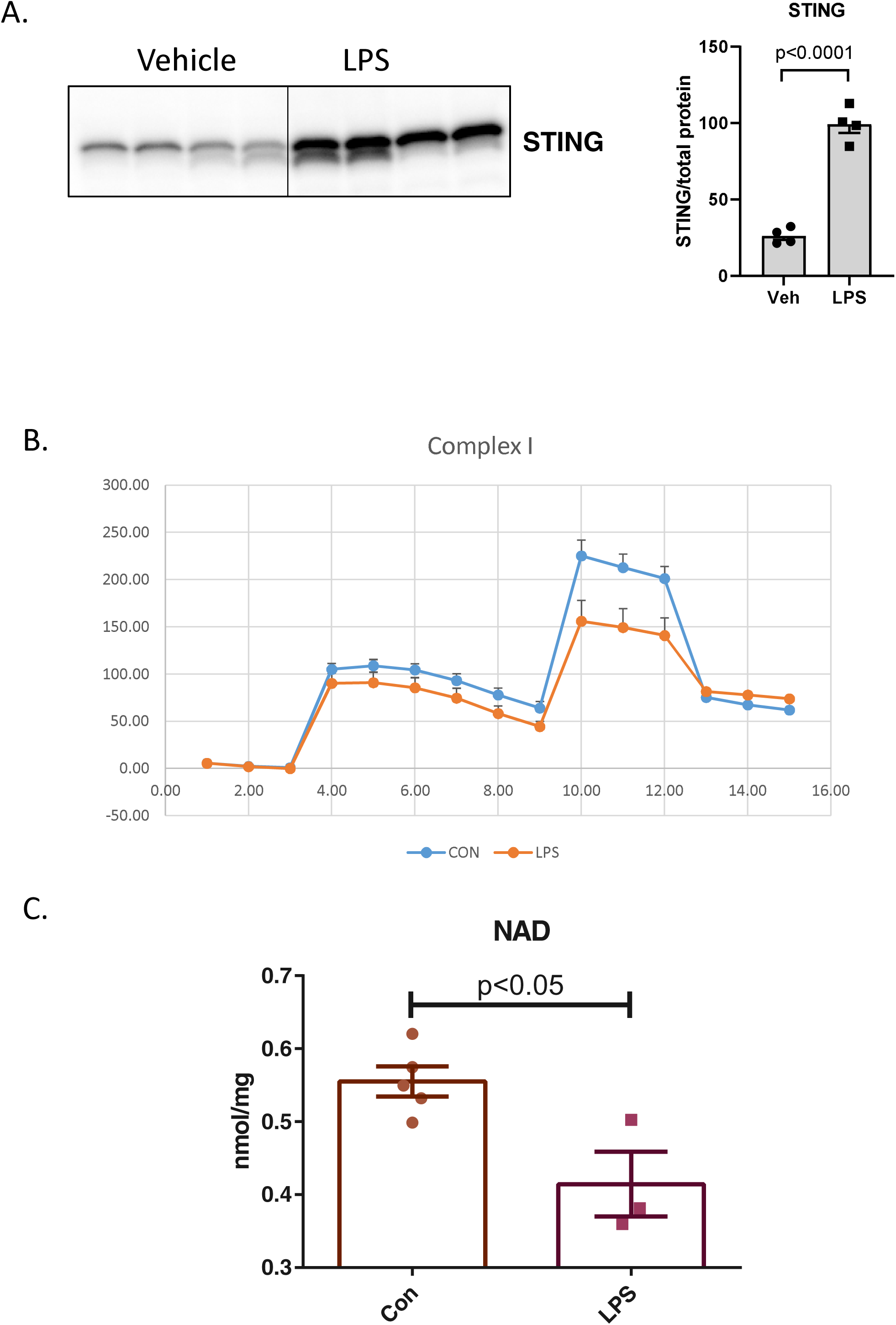
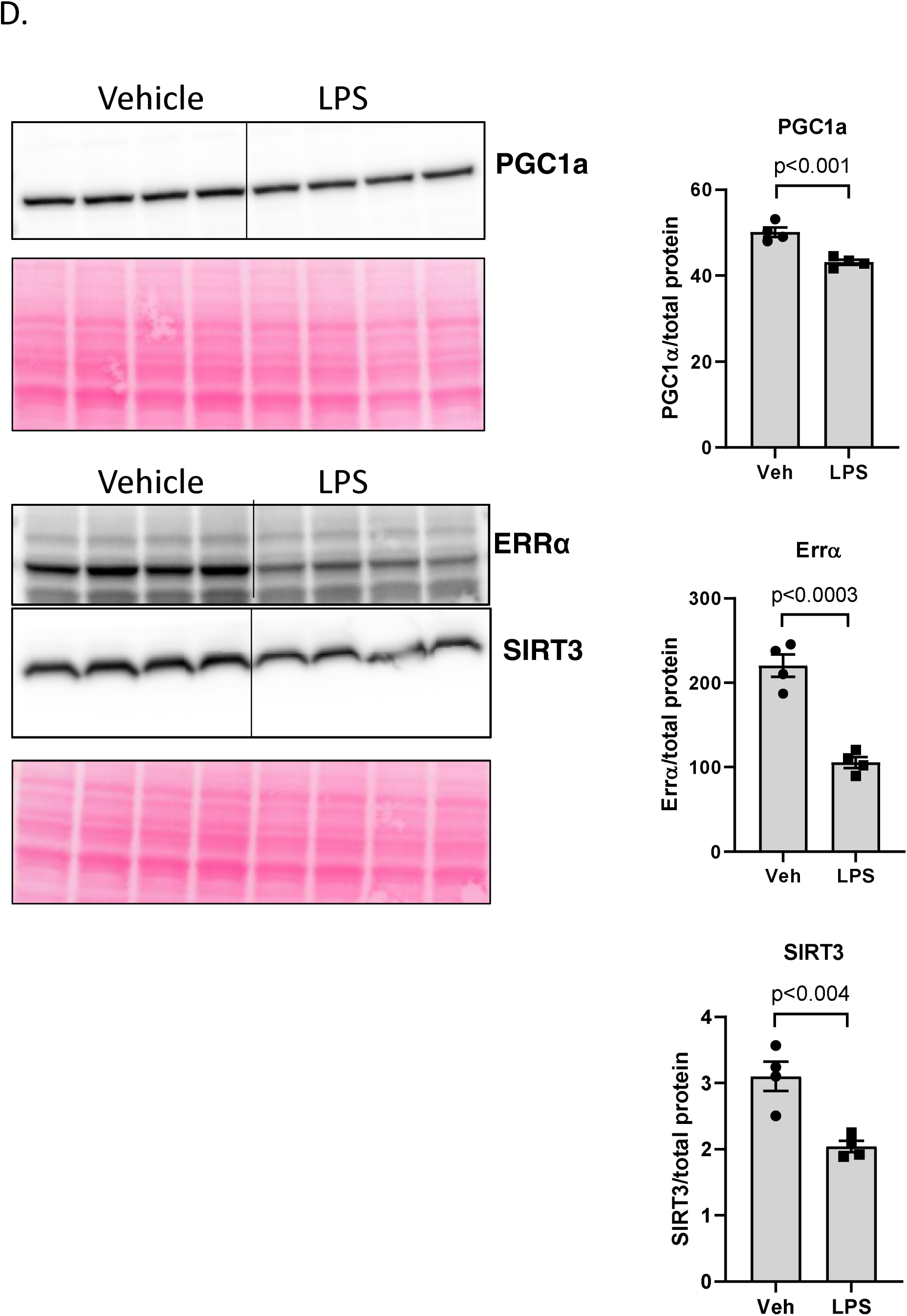
Figure S3

## Notes

### Competing Interest Statement

The authors have declared no competing interest.

### Summary of Updates

More data have been added.

## REFERENCES

1. Sedor, JR, Freedman, BI: Biologic Underpinnings of Type 1 Diabetic Kidney Disease. J Am Soc Nephrol, 30: 1782–1783, 2019.

2. Thomas, MC, Cooper, ME, Zimmet, P: Changing epidemiology of type 2 diabetes mellitus and associated chronic kidney disease. Nat Rev Nephrol, 12: 73–81, 2016.

3. Breyer, MD, Coffman, TM, Flessner, MF, Fried, LF, Harris, RC, Ketchum, CJ, Kretzler, M, Nelson, RG, Sedor, JR, Susztak, K, Kidney Research National, D: Diabetic nephropathy: a national dialogue. Clin J Am Soc Nephrol, 8: 1603–1605, 2013.

4. Boyle, JP, Thompson, TJ, Gregg, EW, Barker, LE, Williamson, DF: Projection of the year 2050 burden of diabetes in the US adult population: dynamic modeling of incidence, mortality, and prediabetes prevalence. Popul Health Metr, 8: 29, 2010.

5. Reidy, K, Kang, HM, Hostetter, T, Susztak, K: Molecular mechanisms of diabetic kidney disease. J Clin Invest, 124: 2333–2340, 2014.

6. Gregg, EW, Li, Y, Wang, J, Burrows, NR, Ali, MK, Rolka, D, Williams, DE, Geiss, L: Changes in diabetes-related complications in the United States, 1990-2010. N Engl J Med, 370: 1514–1523, 2014.

7. Barrera-Chimal, J, Jaisser, F: Pathophysiologic mechanisms in diabetic kidney disease: A focus on current and future therapeutic targets. Diabetes Obes Metab, 22 Suppl 1: 16–31, 2020.

8. Matthews, DR, Paldanius, PM, Proot, P, Chiang, Y, Stumvoll, M, Del Prato, S, group, Vs: Glycaemic durability of an early combination therapy with vildagliptin and metformin versus sequential metformin monotherapy in newly diagnosed type 2 diabetes (VERIFY): a 5-year, multicentre, randomised, double-blind trial. Lancet, 394: 1519–1529, 2019.

9. Davies, MJ, D’Alessio, DA, Fradkin, J, Kernan, WN, Mathieu, C, Mingrone, G, Rossing, P, Tsapas, A, Wexler, DJ, Buse, JB: Management of Hyperglycemia in Type 2 Diabetes, 2018. A Consensus Report by the American Diabetes Association (ADA) and the European Association for the Study of Diabetes (EASD). Diabetes Care, 41: 2669–2701, 2018.

10. Alicic, RZ, Rooney, MT, Tuttle, KR: Diabetic Kidney Disease: Challenges, Progress, and Possibilities. Clin J Am Soc Nephrol, 12: 2032–2045, 2017.

11. Perkovic, V, Jardine, MJ, Neal, B, Bompoint, S, Heerspink, HJL, Charytan, DM, Edwards, R, Agarwal, R, Bakris, G, Bull, S, Cannon, CP, Capuano, G, Chu, PL, de Zeeuw, D, Greene, T, Levin, A, Pollock, C, Wheeler, DC, Yavin, Y, Zhang, H, Zinman, B, Meininger, G, Brenner, BM, Mahaffey, KW, Investigators, CT: Canagliflozin and Renal Outcomes in Type 2 Diabetes and Nephropathy. N Engl J Med, 380: 2295–2306, 2019.

12. Michos, ED, Tuttle, KR: GLP-1 Receptor Agonists in Diabetic Kidney Disease. Clin J Am Soc Nephrol, 2021.

13. Winiarska, A, Knysak, M, Nabrdalik, K, Gumprecht, J, Stompor, T: Inflammation and Oxidative Stress in Diabetic Kidney Disease: The Targets for SGLT2 Inhibitors and GLP-1 Receptor Agonists. Int J Mol Sci, 22, 2021.

14. Andrade-Oliveira, V, Foresto-Neto, O, Watanabe, IKM, Zatz, R, Camara, NOS: Inflammation in Renal Diseases: New and Old Players. Front Pharmacol, 10: 1192, 2019.

15. Wang, XX, Wang, D, Luo, Y, Myakala, K, Dobrinskikh, E, Rosenberg, AZ, Levi, J, Kopp, JB, Field, A, Hill, A, Lucia, S, Qiu, L, Jiang, T, Peng, Y, Orlicky, D, Garcia, G, Herman-Edelstein, M, D’Agati, V, Henriksen, K, Adorini, L, Pruzanski, M, Xie, C, Krausz, KW, Gonzalez, FJ, Ranjit, S, Dvornikov, A, Gratton, E, Levi, M: FXR/TGR5 Dual Agonist Prevents Progression of Nephropathy in Diabetes and Obesity. J Am Soc Nephrol, 29: 118–137, 2018.

16. Myakala, K, Jones, BA, Wang, XX, Levi, M: Sacubitril/valsartan treatment has differential effects in modulating diabetic kidney disease in db/db mice and KKAy mice compared with valsartan treatment. Am J Physiol Renal Physiol, 320: F1133–F1151, 2021.

17. Marchi, S, Guilbaud, E, Tait, SWG, Yamazaki, T, Galluzzi, L: Mitochondrial control of inflammation. Nat Rev Immunol, 2022.

18. Ji, D, Yin, JY, Li, DF, Zhu, CT, Ye, JP, Pan, YQ: Effects of inflammatory and anti-inflammatory environments on the macrophage mitochondrial function. Sci Rep, 10: 20324, 2020.

19. Ahmad, AA, Draves, SO, Rosca, M: Mitochondria in Diabetic Kidney Disease. Cells, 10, 2021.

20. Galvan, DL, Mise, K, Danesh, FR: Mitochondrial Regulation of Diabetic Kidney Disease. Front Med (Lausanne), 8: 745279, 2021.

21. Rajman, L, Chwalek, K, Sinclair, DA: Therapeutic Potential of NAD-Boosting Molecules: The In Vivo Evidence. Cell Metab, 27: 529–547, 2018.

22. Nacarelli, T, Zhang, R: NAD(+) metabolism controls inflammation during senescence. Mol Cell Oncol, 6: 1605819, 2019.

23. Hou, Y, Wei, Y, Lautrup, S, Yang, B, Wang, Y, Cordonnier, S, Mattson, MP, Croteau, DL, Bohr, VA: NAD(+) supplementation reduces neuroinflammation and cell senescence in a transgenic mouse model of Alzheimer’s disease via cGAS-STING. Proc Natl Acad Sci U S A, 118, 2021.

24. Zhou, B, Wang, DD, Qiu, Y, Airhart, S, Liu, Y, Stempien-Otero, A, O’Brien, KD, Tian, R: Boosting NAD level suppresses inflammatory activation of PBMCs in heart failure. J Clin Invest, 130: 6054–6063, 2020.

25. Minhas, PS, Liu, L, Moon, PK, Joshi, AU, Dove, C, Mhatre, S, Contrepois, K, Wang, Q, Lee, BA, Coronado, M, Bernstein, D, Snyder, MP, Migaud, M, Majeti, R, Mochly-Rosen, D, Rabinowitz, JD, Andreasson, KI: Macrophage de novo NAD(+) synthesis specifies immune function in aging and inflammation. Nat Immunol, 20: 50–63, 2019.

26. Chatterjee, S, Balram, A, Li, W: Convergence: Lactosylceramide-Centric Signaling Pathways Induce Inflammation, Oxidative Stress, and Other Phenotypic Outcomes. Int J Mol Sci, 22, 2021.

27. Schneider, WM, Chevillotte, MD, Rice, CM: Interferon-stimulated genes: a complex web of host defenses. Annu Rev Immunol, 32: 513–545, 2014.

28. Decout, A, Katz, JD, Venkatraman, S, Ablasser, A: The cGAS-STING pathway as a therapeutic target in inflammatory diseases. Nat Rev Immunol, 2021.

29. Ablasser, A, Hur, S: Regulation of cGAS- and RLR-mediated immunity to nucleic acids. Nat Immunol, 21: 17–29, 2020.

30. Ablasser, A, Chen, ZJ: cGAS in action: Expanding roles in immunity and inflammation. Science, 363, 2019.

31. Chung, KW, Dhillon, P, Huang, S, Sheng, X, Shrestha, R, Qiu, C, Kaufman, BA, Park, J, Pei, L, Baur, J, Palmer, M, Susztak, K: Mitochondrial Damage and Activation of the STING Pathway Lead to Renal Inflammation and Fibrosis. Cell Metab, 30: 784–799 e785, 2019.

32. Haag, SM, Gulen, MF, Reymond, L, Gibelin, A, Abrami, L, Decout, A, Heymann, M, van der Goot, FG, Turcatti, G, Behrendt, R, Ablasser, A: Targeting STING with covalent small-molecule inhibitors. Nature, 559: 269–273, 2018.

33. Maekawa, H, Inoue, T, Ouchi, H, Jao, TM, Inoue, R, Nishi, H, Fujii, R, Ishidate, F, Tanaka, T, Tanaka, Y, Hirokawa, N, Nangaku, M, Inagi, R: Mitochondrial Damage Causes Inflammation via cGAS-STING Signaling in Acute Kidney Injury. Cell Rep, 29: 1261–1273 e1266, 2019.

34. Shimazu, T, Hirschey, MD, Hua, L, Dittenhafer-Reed, KE, Schwer, B, Lombard, DB, Li, Y, Bunkenborg, J, Alt, FW, Denu, JM, Jacobson, MP, Verdin, E: SIRT3 deacetylates mitochondrial 3-hydroxy-3-methylglutaryl CoA synthase 2 and regulates ketone body production. Cell Metab, 12: 654–661, 2010.

35. Hirschey, MD, Shimazu, T, Goetzman, E, Jing, E, Schwer, B, Lombard, DB, Grueter, CA, Harris, C, Biddinger, S, Ilkayeva, OR, Stevens, RD, Li, Y, Saha, AK, Ruderman, NB, Bain, JR, Newgard, CB, Farese, RV, Jr., Alt, FW, Kahn, CR, Verdin, E: SIRT3 regulates mitochondrial fatty-acid oxidation by reversible enzyme deacetylation. Nature, 464: 121–125, 2010.

36. Someya, S, Yu, W, Hallows, WC, Xu, J, Vann, JM, Leeuwenburgh, C, Tanokura, M, Denu, JM, Prolla, TA: Sirt3 mediates reduction of oxidative damage and prevention of age-related hearing loss under caloric restriction. Cell, 143: 802–812, 2010.

37. Tao, R, Vassilopoulos, A, Parisiadou, L, Yan, Y, Gius, D: Regulation of MnSOD enzymatic activity by Sirt3 connects the mitochondrial acetylome signaling networks to aging and carcinogenesis. Antioxid Redox Signal, 20: 1646–1654, 2014.

38. Fernandez-Marcos, PJ, Auwerx, J: Regulation of PGC-1alpha, a nodal regulator of mitochondrial biogenesis. Am J Clin Nutr, 93: 884S–890, 2011.

39. Shults NV, Kanovka SS, Ten Eyck JE, Rybka V, Suzuki YJ. Ultrastructural Changes of the Right Ventricular Myocytes in Pulmonary Arterial Hypertension. J Am Heart Assoc, 5;8(5), 2019.

40. Mise, K, Galvan, DL, Danesh, FR: Shaping Up Mitochondria in Diabetic Nephropathy. Kidney 360, 1: 982–992, 2020.

41. Tang, SCW, Yiu, WH: Innate immunity in diabetic kidney disease. Nat Rev Nephrol, 16: 206–222, 2020.

42. Forbes, JM, Thorburn, DR: Mitochondrial dysfunction in diabetic kidney disease. Nat Rev Nephrol, 14: 291–312, 2018.

43. Sharma, K, Karl, B, Mathew, AV, Gangoiti, JA, Wassel, CL, Saito, R, Pu, M, Sharma, S, You, YH, Wang, L, Diamond-Stanic, M, Lindenmeyer, MT, Forsblom, C, Wu, W, Ix, JH, Ideker, T, Kopp, JB, Nigam, SK, Cohen, CD, Groop, PH, Barshop, BA, Natarajan, L, Nyhan, WL, Naviaux, RK: Metabolomics reveals signature of mitochondrial dysfunction in diabetic kidney disease. J Am Soc Nephrol, 24: 1901–1912, 2013.

44. Tran, MT, Zsengeller, ZK, Berg, AH, Khankin, EV, Bhasin, MK, Kim, W, Clish, CB, Stillman, IE, Karumanchi, SA, Rhee, EP, Parikh, SM: PGC1alpha drives NAD biosynthesis linking oxidative metabolism to renal protection. Nature, 531: 528–532, 2016.

45. Yasuda, I, Hasegawa, K, Sakamaki, Y, Muraoka, H, Kawaguchi, T, Kusahana, E, Ono, T, Kanda, T, Tokuyama, H, Wakino, S, Itoh, H: Pre-emptive Short-term Nicotinamide Mononucleotide Treatment in a Mouse Model of Diabetic Nephropathy. J Am Soc Nephrol, 32: 1355–1370, 2021.

46. Srivastava, SP, Li, J, Takagaki, Y, Kitada, M, Goodwin, JE, Kanasaki, K, Koya, D: Endothelial SIRT3 regulates myofibroblast metabolic shifts in diabetic kidneys. iScience, 24: 102390, 2021.

47. Srivastava, SP, Li, J, Kitada, M, Fujita, H, Yamada, Y, Goodwin, JE, Kanasaki, K, Koya, D: SIRT3 deficiency leads to induction of abnormal glycolysis in diabetic kidney with fibrosis. Cell Death Dis, 9: 997, 2018.

48. Morigi, M, Perico, L, Rota, C, Longaretti, L, Conti, S, Rottoli, D, Novelli, R, Remuzzi, G, Benigni, A: Sirtuin 3-dependent mitochondrial dynamic improvements protect against acute kidney injury. J Clin Invest, 125: 715–726, 2015.

49. Miguel, V, Tituana, J, Herrero, JI, Herrero, L, Serra, D, Cuevas, P, Barbas, C, Puyol, DR, Marquez-Exposito, L, Ruiz-Ortega, M, Castillo, C, Sheng, X, Susztak, K, Ruiz-Canela, M, Salas-Salvado, J, Gonzalez, MAM, Ortega, S, Ramos, R, Lamas, S: Renal tubule Cpt1a overexpression protects from kidney fibrosis by restoring mitochondrial homeostasis. J Clin Invest, 131, 2021.

50. Xu, W, Boyd, RM, Tree, MO, Samkari, F, Zhao, L: Mitochondrial transcription factor A promotes DNA strand cleavage at abasic sites. Proc Natl Acad Sci U S A, 116: 17792–17799, 2019.

51. Wu, J, Raman, A, Coffey, NJ, Sheng, X, Wahba, J, Seasock, MJ, Ma, Z, Beckerman, P, Laczko, D, Palmer, MB, Kopp, JB, Kuo, JJ, Pullen, SS, Boustany-Kari, CM, Linkermann, A, Susztak, K: The key role of NLRP3 and STING in APOL1-associated podocytopathy. J Clin Invest, 131, 2021.

52. Furda, A, Santos, JH, Meyer, JN, Van Houten, B: Quantitative PCR-based measurement of nuclear and mitochondrial DNA damage and repair in mammalian cells. Methods Mol Biol, 1105: 419–437, 2014.

53. Ginley, B, Lutnick, B, Jen, KY, Fogo, AB, Jain, S, Rosenberg, A, Walavalkar, V, Wilding, G, Tomaszewski, JE, Yacoub, R, Rossi, GM, Sarder, P: Computational Segmentation and Classification of Diabetic Glomerulosclerosis. J Am Soc Nephrol, 30: 1953–1967, 2019.

54. Wang, XX, Jiang, T, Shen, Y, Adorini, L, Pruzanski, M, Gonzalez, FJ, Scherzer, P, Lewis, L, Miyazaki-Anzai, S, Levi, M: The farnesoid X receptor modulates renal lipid metabolism and diet-induced renal inflammation, fibrosis, and proteinuria. Am J Physiol Renal Physiol, 297: F1587–1596, 2009.

55. Weibel, ER, Kistler, GS, Scherle, WF: Practical stereological methods for morphometric cytology. J Cell Biol, 30: 23–38, 1966.

56. Bolger, AM, Lohse, M, Usadel, B: Trimmomatic: a flexible trimmer for Illumina sequence data. Bioinformatics, 30: 2114–2120, 2014.

57. Dobin, A, Davis, CA, Schlesinger, F, Drenkow, J, Zaleski, C, Jha, S, Batut, P, Chaisson, M, Gingeras, TR: STAR: ultrafast universal RNA-seq aligner. Bioinformatics, 29: 15–21, 2013.

58. Li, B, Dewey, CN: RSEM: accurate transcript quantification from RNA-Seq data with or without a reference genome. BMC Bioinformatics, 12: 323, 2011.

59. Love, MI, Huber, W, Anders, S: Moderated estimation of fold change and dispersion for RNA-seq data with DESeq2. Genome Biol, 15: 550, 2014.

60. Huang da, W, Sherman, BT, Lempicki, RA: Systematic and integrative analysis of large gene lists using DAVID bioinformatics resources. Nat Protoc, 4: 44–57, 2009.

61. Huang, DW, Sherman, BT, Tan, Q, Collins, JR, Alvord, WG, Roayaei, J, Stephens, R, Baseler, MW, Lane, HC, Lempicki, RA: The DAVID Gene Functional Classification Tool: a novel biological module-centric algorithm to functionally analyze large gene lists. Genome Biol, 8: R183, 2007.

62. Rath, S, Sharma, R, Gupta, R, Ast, T, Chan, C, Durham, TJ, Goodman, RP, Grabarek, Z, Haas, ME, Hung, WHW, Joshi, PR, Jourdain, AA, Kim, SH, Kotrys, AV, Lam, SS, McCoy, JG, Meisel, JD, Miranda, M, Panda, A, Patgiri, A, Rogers, R, Sadre, S, Shah, H, Skinner, OS, To, TL, Walker, MA, Wang, H, Ward, PS, Wengrod, J, Yuan, CC, Calvo, SE, Mootha, VK: MitoCarta3.0: an updated mitochondrial proteome now with sub-organelle localization and pathway annotations. Nucleic Acids Res, 49: D1541–D1547, 2021.

63. Tahir, R, Renuse, S, Udainiya, S, Madugundu, AK, Cutler, JA, Nirujogi, RS, Na, CH, Xu, Y, Wu, X, Pandey, A: Mutation-Specific and Common Phosphotyrosine Signatures of KRAS G12D and G13D Alleles. J Proteome Res, 20: 670–683, 2021.

64. Tyanova, S, Cox, J: Perseus: A Bioinformatics Platform for Integrative Analysis of Proteomics Data in Cancer Research. Methods Mol Biol, 1711: 133–148, 2018.

