## Supplemental figure legend for "NAD metabolism modulates inflammation and mitochondria function in diabetic kidney disease"

**Supplementary Figure S1:** Effect of nicotinamide riboside (NR) treatment on expression of fibronectin in kidney of db/db mice. Western blot analysis indicating fibronectin protein expression levels were higher in db/db mice and the levels were normalized upon NR treatment. n=6 per group, values presented as mean ± SEM with variance is calculated using one-way ANOVA.

**Supplementary Figure S2:** Protein-protein-interaction network analysis of immune-related genes that are corrected by NR treatment.

**Supplementary Figure S3:** Effect of LPS (5mg/kg body weight) on inflammation and mitochondria. A) expression of STING protein by western blot. B) Oxygen consumption analysis of isolated mitochondria from kidneys. C) NAD+ level measured with colorimetric assay kit. D) Western blot analysis of PGC1α, ERRα, and Sirt3 expression in the kidneys. n=4 per group, values presented as mean ± SEM with variance is calculated using student t-test.
