## supplemental table 1 for "NAD metabolism modulates inflammation and mitochondria function in diabetic kidney disease"

**Supplementary Table S1: List of resources.**

**List of primers**

| **Name** | **Forward** | **Reverse** |
| --- | --- | --- |
| MCP-1 | TTAAAAACCTGGATCGGAACCAA | GCATTAGCTTCAGATTTACGGGT |
| TNF-α | CCCTCACACTCAGATCATCTTCT | GCTACGACGTGGGCTACAG |
| PAI1 | GACTCTGGATGAGTGGAAAGC | GGCGGCCTAAGTCTCCAAAAT |
| Tgfβ | CTCCCGTGGCTTCTAGTGC | GCCTTAGTTTGGACAGGATCTG |
| CTGF | GGGCCTCTTCTGCGATTTC | ATCCAGGCAAGTGCATTGGTA |
| Nox4 | GTGAAGATTTGCCTGGAAGAA | GATGATTGATGACTGAGATGATGG |
| mtDNA | ATAACCGAGTCGTTCTGCCAAT | TTTCAGAGCATTGGCCATAGAA |
| PGC1α | GTCAGAGTGGATTGGAGTTG | AAGTCATTCACATCAAGTTCAG |
| Nrf1 | AGCACGGAGTGACCCAAAC | AGGATGTCCGAGTCATCATAAGA |
| Tfam | AACACCCAGATGCAAAACTTTCA | GACTTGGAGTTAGCTGCTCTTT |
| Ndufa4 | TCCCAGCTTGATTCCTCTCTT | GGGTTGTTCTTTCTGTCCCAG |
| Uqcrc2 | AAAGTTGCCCCGAAGGTTAAA | GAGCATAGTTTTCCAGAGAAGCA |
| Cox6a2 | CTGCTCCCTTAACTGCTGGAT | GATTGTGGAAAAGCGTGTGGT |
| Cpt1a | CTCCGCCTGAGCCATGAAG | CACCAGTGATGATGCCATTCT |
| LCAD | ACTACTGTGCTTCAGGGACAA | GCAAAGGACTTCGATTCTGCC |
| MCAD | AACACTTACTATGCCTCGATTGCA | CCATAGCCTCCGAAAATCTGAA |
| Il6 | TAGTCCTTCCTACCCCAATTTCC | TTGGTCCTTAGCCACTCCTTC |
| Timp1 | GCAACTCGGACCTGGTCATAA | CGGCCCGTGATGAGAAACT |
| TLR2 | GCAAACGCTGTTCTGCTCAG | AGGCGTCTCCCTCTATTGTATT |
| cGAS | GAGGCGCGGAAAGTCGTAA | TTGTCCGGTTCCTTCCTGGA |
| TMEM173 | GGTCACCGCTCCAAATATGTAG | CAGTAGTCCAAGTTCGTGCGA |
| CD68 | TGTCTGATCTTGCTAGGACCG | GAGAGTAACGGCCTTTTTGTGA |

| **Reagent Name** | **Source** | **Identifier** |
| --- | --- | --- |
| Collagen IV | Rockland | 600-401-106 |
| Fibronectin | Sigma | F3648 |
| P57 | Abcam | ab75974 |
| 4-HNE | Abcam | ab46545 |
| Acetylated Lysine | Cell Signaling | 9441 |
| SIRT3 | Cell Signaling | D22A3 |
| SIRT3 IHC | Sigma Aldrich | S4072 |
| Acetylated SOD2 (k68) | Abcam | ab137037 |
| SOD2 | Cell signaling | 13141 |
| Acetylated IDH2 | Genetel Lab | AC0004 |
| PGC1a | Millipore | AB3242 |
| MCAD | Abcam | ab110296 |
| CD45 | BD Biosciences | 550539 |
| CD68 | Bio-Rad | MCA1957 |
| Sting (mouse) | Cell Signaling | 50494s |
| Sting (human) | Cell Signaling | 13647s |
| p-TBK1 | Cell Signaling | 5483 |
| TBK1 | Cell Signaling | 3504 |
| p-IRF3 | Cell Signaling | 29047 |
| IRF3 | Cell Signaling | 4302 |
| p-Stat3 | Cell Signaling | 9145 |
| Stat3 | Cell Signaling | 4904 |
| p-p65 | Cell signaling | 3036 |
| P65 | Cell Signaling | 8242 |

**List of antibodies**

**List of assays**

| **Name** | **Source** | **Identifier** |
| --- | --- | --- |
| Albumin | Exocell | 1011 |
| Creatinine | BioAssay Systems | DICT-500 |
| SIRT3 assay | Abcam | Ab156067 |
| TBARS | BioAssay Systems | DTBA-100 |
| Mitochondrial isolation kit | Sigma | MITOISO1 |
| Complex I enzyme activity assay kit | Abcam | ab109721 |
| Complex IV enzyme activity assay kit | Abcam | ab109911 |
| BCA protein assay | Thermo Fisher Scientific | 23225 |
| Kim1 assay kit | Abcam | ab119596 |
