## supplemental table 2 for "NAD metabolism modulates inflammation and mitochondria function in diabetic kidney disease"

**Supplementary Table S2: Human subject’s demography.**

| **Characteristics** | **Controls**  **(*n* = 9-11)** | **Diabetic**  **(*n* = 11-13)** |
| --- | --- | --- |
| Age (yr.) | 47.6 (8 - 89) | 59 (42 - 79) |
| Sex (%M) | 54.5 | 61.5 |
| HbA1c | NA | 6.7 (5.8 - 7.5) |
| Proteinuria (UPCR) | 1.0 (0.1480 - 2.790) | 7.3 (2.9 - 22.0) |
| Serum Creatinine | 1.8 (0.4 - 3.7) | 2.9 (1.1 - 8.9) |
| Global Sclerosis (%) | 11.4 (0 - 34) | 30.2 (3 - 57) |
| IFTA (%) | 9 (0 - 25) | 53.5 (10 - 80) |
